# Proteome of the secondary plastid of *Euglena gracilis* reveals metabolic quirks and colourful history

**DOI:** 10.1101/573709

**Authors:** Anna M. G. Novák Vanclová, Martin Zoltner, Steven Kelly, Petr Soukal, Kristína Záhonová, Zoltán Füssy, ThankGod E. Ebenezer, Eva Lacová Dobáková, Marek Eliáš, Julius Lukeš, Mark C. Field, Vladimír Hampl

## Abstract

*Euglena gracilis* is a well-studied biotechnologically exploitable phototrophic flagellate harbouring secondary green plastids. Here we describe its plastid proteome obtained by high-resolution proteomics. We identified 1,345 candidate plastid proteins and assigned functional annotations to 774 of them. More than 120 proteins are affiliated neither to the host lineage nor the plastid ancestor and may represent horizontal acquisitions from various algal and prokaryotic groups. Reconstruction of plastid metabolism confirms both the presence of previously studied/predicted enzymes/pathways and also provides direct evidence for unusual features of its metabolism including uncoupling of carotenoid and phytol metabolism, a limited role in amino acid metabolism and the presence of two sets of the SUF pathway for FeS cluster assembly. Most significantly, one of these was acquired by lateral gene transfer (LGT) from the chlamydiae. Plastidial paralogs of membrane trafficking-associated proteins likely mediating a poorly understood fusion of transport vesicles with the outermost plastid membrane were identified, as well as derlin-related proteins that potentially act as protein translocases of the middle membrane, supporting an extremely simplified TIC complex. The proposed innovations may be also linked to specific features of the transit peptide-like regions described here. Hence the Euglena plastid is demonstrated to be a product of several genomes and to combine novel and conserved metabolism and transport processes.

## Introduction

Euglenids are a diverse group of flagellates belonging to the phylum Euglenozoa, which, together with heteroloboseans and jakobids, comprise the Discoba (Leander et al., 2017). Photosynthetic euglenids, also known as euglenophytes, represent a clade of distinctive, aesthetically pleasing organisms with relatively large cells (up to 200 μm in length), which made them one of the first protists to be discovered and described. The first documented observation of a euglenophyte, presumably *Euglena viridis*, and description of its unique euglenoid movement called metaboly, was carried out by Harris at the end of the 17^th^ century (1695) and the first species of euglenophytes were described in the first half of the 19^th^ century by Ehrenberg (1830). Nutritional strategies amongst euglenids include bacteriovory, eukaryovory, photoautotrophy, and both primary and secondary osmotrophy, the latter being the case for several species that have lost the ability to photosynthesize but at least in some cases retain colourless plastids (Leander et al. 2001; Leander 2004). The biology of heterotrophic euglenids remains relatively unexplored, but the ease of collection and cultivation of photosynthetic members has made them one of the most widely studied protist groups.

Euglenophytes harbour green, triple membrane-bound plastids that evolved approx. 500 Mya from a secondary endosymbiont belonging to the green algal lineage Pyramimonadales (Turmel et al. 2009; Jackson et al. 2018). However, it is possible that this symbiont was not the only one to share living space and genes with the common ancestors of euglenophytes, as a significant proportion of euglenophyte genes are related to various algal groups, suggesting extensive lateral, possibly endosymbiotic, gene transfer prior to establishment of the current plastid (Maruyama et al. 2011; Markunas and Triemer 2016; Lakey and Triemer 2017; Ponce-Toledo et al. 2018, Ebenezer et al., 2019). This concept, referred to as the shopping bag hypothesis, explains the presence of green algae-derived genes in red algal plastid-containing lineages and *vice versa* (Larkum et al. 2007).

Secondary and higher-order endosymbioses are highly intriguing evolutionary phenomena that have improved our understanding of the molecular basis of symbiotic relationships and organellogenesis in general. Transfer of genes from the endosymbiont to the host nucleus and establishment of new routes for targeting proteins to the organelle are crucial steps for endosymbiont transformation into a fully integrated organelle. The features of protein targeting systems of secondary plastids are remarkably similar in otherwise unrelated lineages (Durnford and Gray 2006; Sommer et al. 2007; Hempel et al. 2009; Spork et al. 2009; Minge et al. 2010; Felsner et al. 2011; Lau et al. 2016) which indicates general cytological constraints and convergent evolution rather than shared evolutionary history. In general, components of the secretory pathway, including the signal-recognition particle, endoplasmic reticulum (ER) and/or Golgi-derived vesicles, have been recruited for protein translocation across the additional (one or two outermost) membranes of secondary plastids. In multiple lineages of “chromalveolates” whose plastids are presumed of a single origin but not inherited vertically but rather via a series of endosymbiotic events (Stiller et al. 2014), a duplicated version of the ER-associated protein degradation (ERAD) pathway is present. Meanwhile, the major protein translocases of the primary plastids, the TOC/TIC complexes, retain their functions at the inner two membranes of secondary plastids, homologous to the envelope of the primary plastid (van Dooren and Striepen 2013; Sheiner and Striepen 2013; Archibald 2015; Gould et al. 2015; Maier et al. 2015; Bölter and Soll 2016).

In euglenophytes, vesicular transport between the ER, Golgi, and plastids has been observed (Sulli et al. 1999) and even reconstituted *in vitro* (Sláviková et al. 2005), but the molecular machinery mediating vesicle docking and fusion with the outer plastid membrane is uncharacterized. In addition, a plant-like plastid targeting signal (transit peptide) was identified and experimentally shown to be essential for the *in vitro* import into the plastid of at least some proteins, which implies the presence of a TOC/TIC-like pathway in the inner membranes (Sláviková et al. 2005). However, a recent bioinformatic analysis of available euglenophyte transcriptomic data identified homologs of only a few TIC subunits, while the key components of the complex as well as homologs of all TOC subunits are apparently missing (Záhonová et al. 2018, Ebenezer et al., 2019). This suggests the existence of a highly divergent or alternative translocon mediating protein import into the euglenophyte plastid.

*E. gracilis* is a well-established laboratory model and its ultrastructure and metabolism, as well as its plastid and mitochondrial genomes and their evolution have been studied in great detail (Hallick et al. 1993; Doetsch et al. 2001; Geimer et al. 2009; Dobáková et al. 2015; Schwartzbach and Shigeoka 2017). However, the sequence of the nuclear genome has remained incomplete due to its relatively large size and high proportion of non-coding and repetitive DNA. Nevertheless, analyses of available data revealed peculiar features of its molecular genetics including unusual introns and extensive *cis*- and *trans*-splicing of transcripts (Tessier et al. 1991; Muchhal and Schwartzbach 1994; Jenkins et al. 1995; Mateášiková-Kováčová et al. 2012; Kuo et al. 2013; Gumińska et al. 2018). Three transcriptome-derived datasets have been generated in recent years by O’Neill, Trick, Hill, et al. (2015), Yoshida et al. (2016), and Ebenezer et. al. (2019), the latter coupled with a draft genome and used as the main reference dataset in this study. These sequence data and the characteristic features of the plastid-targeting presequences enabled bioinformatic predictions of protein constituents of the *E. gracilis* plastid (Ebenezer et al. 2019; Záhonová et al., 2018). However, direct biochemical approaches are needed to evaluate the accuracy of these *in silico* inferences and to define the *E. gracilis* plastid proteome with confidence.

We present a proteomic analysis of the *E. gracilis* plastid based on high performance liquid chromatography/mass spectrometry analysis of purified organellar fractions. These data provide a new level of insight into the functions, evolution and protein import machinery of this unique organelle. Our analysis underscores the manner in which the *Euglena* plastid differs from its counterparts in other secondary algae and suggests that it holds unexpected or unique characteristics and evolutionary stories.

## Results

### The plastid proteome of *E. gracilis*: general features

The plastid proteome of *E. gracilis* determined here includes 1,345 proteins, 48 of which are encoded by the plastid genome (Hallick et al. 1993). Functional annotation was assigned to 774 proteins, while 571 (42.5%) have unknown function and 109 of these (8%) appear to have no BLAST-identifiable homologs (list of all identified proteins and their detailed annotations are given in supplementary-dataset-1.xlsx). The 774 proteins with functional annotations were sorted into 18 categories and the distribution of log_10_ CP/MT ratios (chloroplast to mitochondrion peptide intensity ratio; see Materials and Methods) in each category was determined (Fig. 1).

**Fig. 1:**
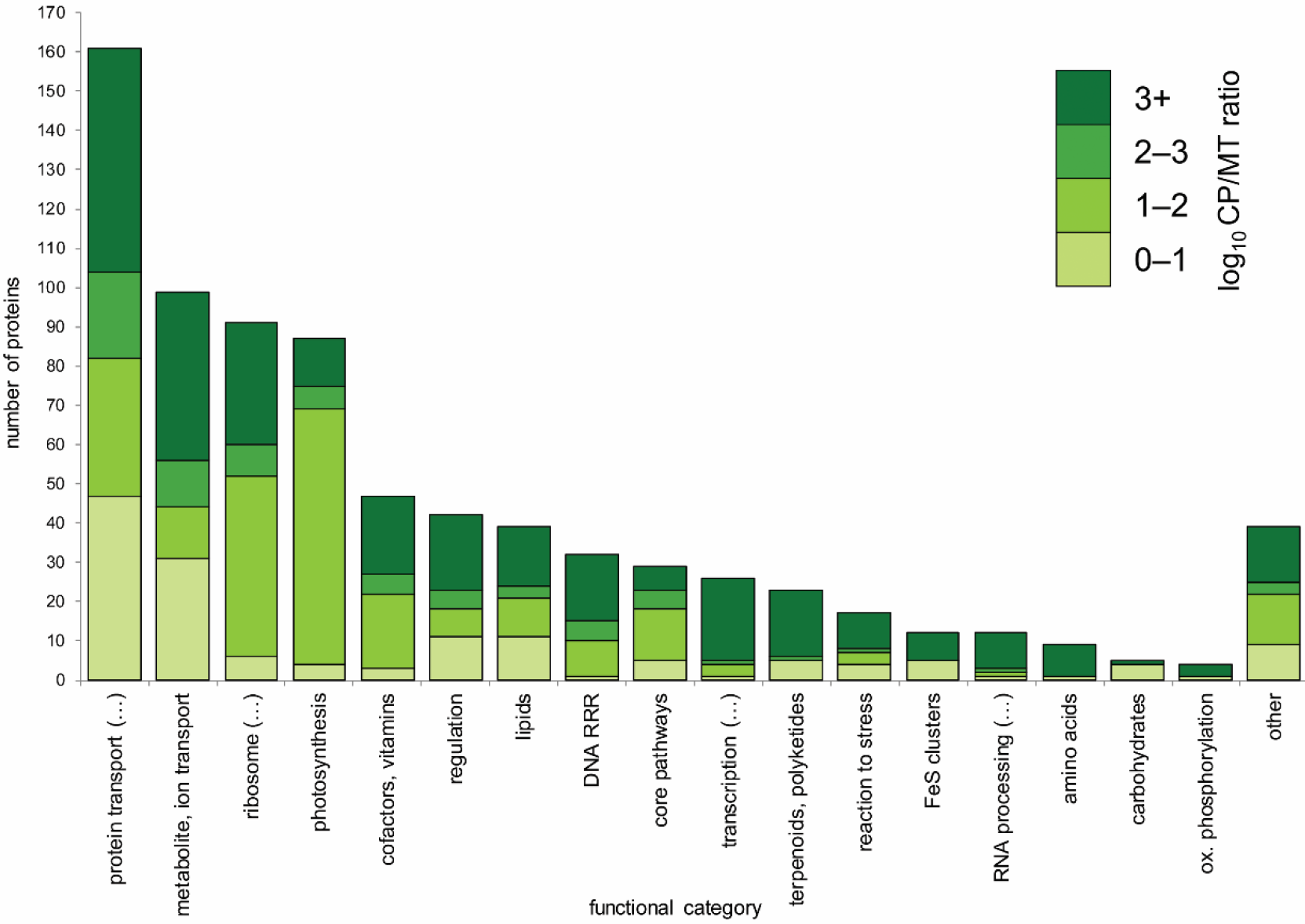
Distribution of functional categories among 774 plastid proteins with predicted function (57.5% of the whole proteome) and log_10_ CP/MT ratio distribution in each category, represented by shades of green (0-1 meaning 1-10× higher amount of the protein in plastid fraction compared to the mitochondrial one, 1-2 for 10-100×, 2-3 for 100-1000×; and >3 for more than 1000× or “infinite” value in case the protein was detected in plastid fraction only; the full category names are listed in Materials and Methods, for detailed description with examples see table S3).

To assess the completeness of the dataset, we looked at the representation of the 67 plastid genome encoded proteins and detected 48 (72%). We further compared the proteome to previous *in silico* predictions of nucleus-encoded plastid-targeted proteins in *E. gracilis*. Using EST data and careful manual analyses, Durnford and Gray (2006) identified 83 non-redundant high-confidence plastid-targeted candidates, of which 74(89%) are present in our proteome. The recent study by Záhonová et al. (2018) focusing on selected functional categories of plastid proteins defined 131 candidates, of which 113 (86%), including some particularly notable examples such as the translation termination factor Rho, not previously identified in the plastid of any eukaryote, were found in our dataset (For details of these comparisons, see supplementary-dataset-2.xlsx). Given that most of the missing proteins perform essential chloroplast functions (components of the photosynthetic apparatus, subunits of the plastidial ribosome etc.), most, if not all, likely represent false negative identifications. Finally, the predicted plastid proteome was compared to a global *in silico* prediction of putative plastid proteins reported recently (Ebenezer et al. 2019), which consists of (1) proteins containing N-terminal sequences adopting the structure typical for the bi-partite plastid-targeting sequences, (2) proteins of clearly plastidial function, or (3) proteins with orthologs in *A. thaliana* plastid proteome. Of 1902 such proteins, only 474 (25%) were detected in the organelle by proteomics. In contrast, 871 proteins identified by proteomics were missing in the *in silico* defined set. The difference reflects the presence of false positives and false negatives in both sets, and perhaps highlights the poor nature of *in silico* predictions in this context.

Using the set of proteins identified in the proteome, we have reconstructed a map of major plastidial metabolic pathways and complexes (see Fig. 2 for overall schematic and Figs. S5.1-11 for detailed pathway reconstructions). Furthermore, we have attempted to reconstruct the set of proteins, putatively involved in the highly derived protein import pathway (Fig. 3). The detailed features of these pathways and systems are discussed in the Discussion section.

**Fig. 2:**
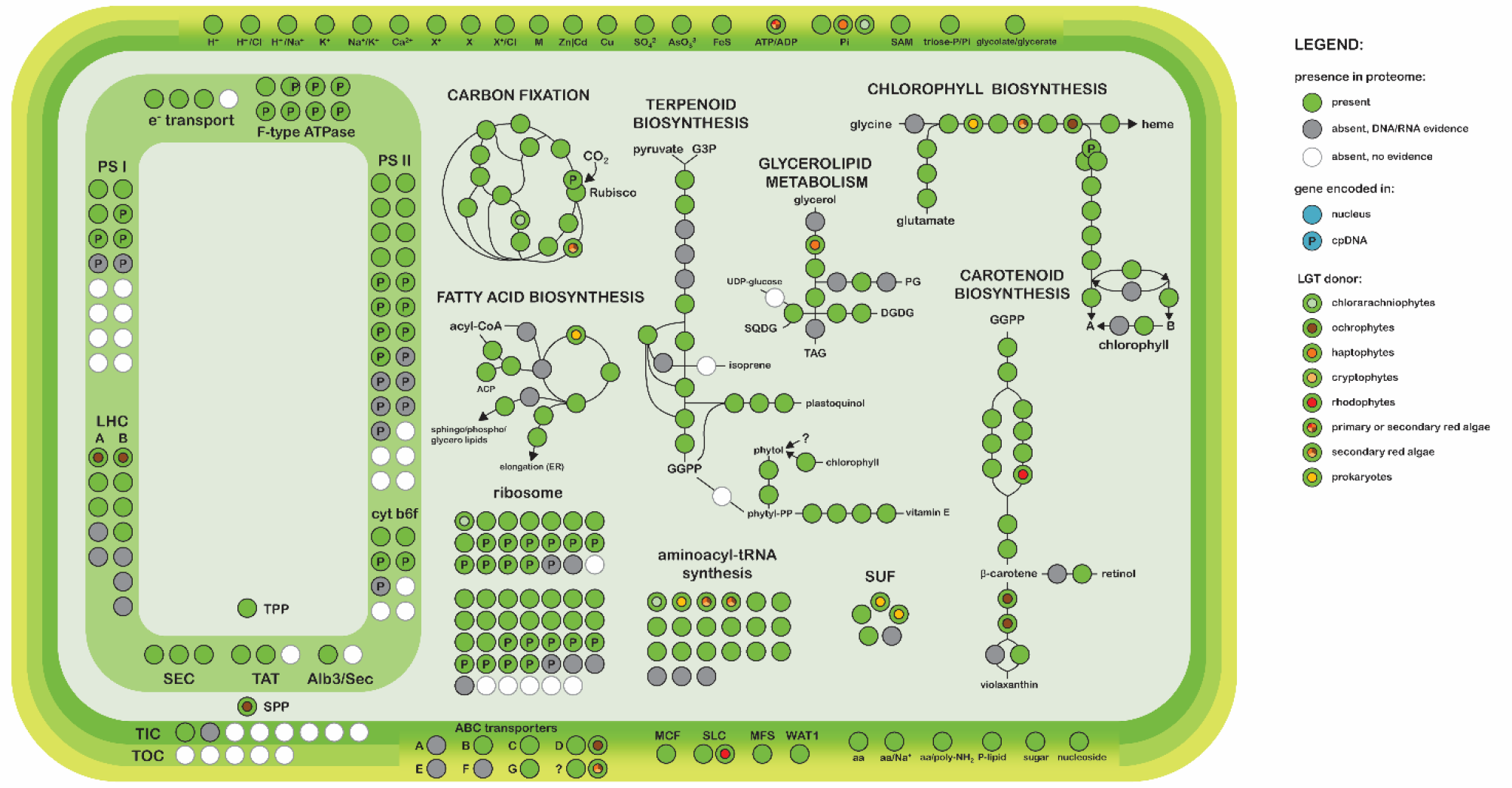
Overview of the *E. gracilis* plastid metabolism as reconstructed from mass spectrometry-based proteome. Enzymes present in the plastid proteome in at least one isoform are marked as green circles, grey circles represent enzymes which were identified on the RNA or DNA level (in this study or previously) but are absent from the proteome; white circles represent genes completely absent in *Euglena*; circles marked by the letter P represent genes coded in the plastid genome while the rest of the circles represent genes coded in the nucleus. Circles with colored dots in the middle represent genes with at least one of their isoforms presumably gained via lateral transfer from a donor group other than green algae and “miscellaneous” algae: pale green for chlorarachniophytes, brown for ochrophytes, dark orange for haptophytes, pale orange for cryptophytes, red for rhodophytes, and a combination of the former in case of proteins of “mixed red” origin (see Fig. S4 for larger and colorblind-friendly version). Multiple overlapping circles represent enzymes composed of multiple subunits with different characteristics. Note that a single protein may be represented by multiple circles due to its role in multiple pathways or reactions. A more detailed schematic including additional pathways and transcript identifiers for each enzyme is available as Figs. S6.1-11.

**Fig. 3:**
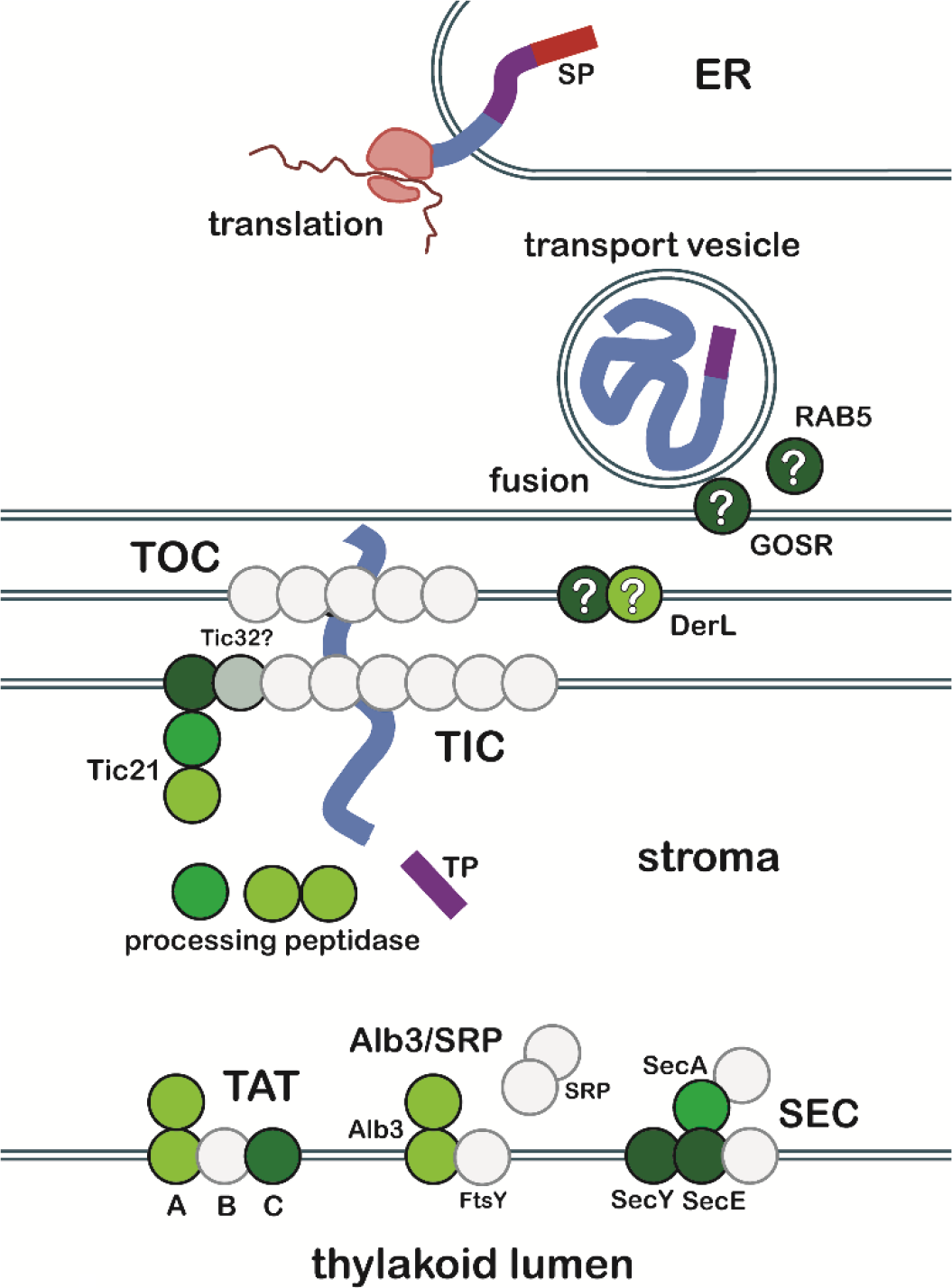
Reconstruction of *E. gracilis* plastid protein import machinery: The N-terminal signal-peptide (SP, red) bearing nuclear-encoded plastid-targeted protein (blue chain) is synthesized on the rough endoplasmic reticulum (RER). If the N-terminus does not contain stop transfer signal (STS), it is co-translationally imported into the ER lumen as indicated. Otherwise, the major part of the protein remains in the cytosol while being anchored in the ER membrane. The SP is cleaved by signal peptidase in the ER lumen. The protein then passes through Golgi and is loaded into/onto a vesicle which eventually fuses with the outermost plastid membrane. Plastidial homologs of GOSR and Rab5 GTPase might mediate the fusion. The protein passes the middle membrane via unknown mechanism dependent on transit peptide (TP, purple) but not TOC complex, possibly employing derlin-like proteins (DerL). Then it passes the inner membrane via highly reduced TIC translocase, possibly consisting of multiple isoforms of a single subunit. The protein is folded and has its transit peptide cleaved in the plastid stroma and, in case additional signal is revealed, enter thylakoid membrane or lumen via TAT, SEC, or SRP/Alb3 pathway. Each circle represents a putative subunit described in model plastids, green circles represent proteins identified in the plastid proteome, different color shades code CP/MT ratio, the darkest shade depicting the most credible evidence for plastid localization, grey circles represent subunits with transcript-level evidence only. Finally, white circles represent subunits which are completely absent.

### Evolutionary origin of plastid proteins

To understand the origin of the genes encoding the plastid proteome, the phylogenetic affiliation of the whole set of plastid proteins was calculated. For 1,152 of these, at least three homologs with an e-value < 10^−3^ were identified in an in-house protein sequence database composed of 208 genomes and transcriptomes from both prokaryotic and eukaryotic taxa. 89 proteins were rejected from further analysis as automatic trimming resulted in an alignment of less than four unique sequences and/or length < 75 aa. A total of 1,063 phylogenetic trees were constructed, but only those in which the branching of an E. gracilis with a taxonomically homogeneous clan (see Materials and Methods) with bootstrap support (BS) ≥ 75% were considered further. In these trees 37 *E. gracilis* proteins were related to sequences from other discobids (kinetoplastids or heteroloboseans) and most probably represent vertical descent. However, many of these proteins are suspected to be contamination from other cellular compartments as suggested by their functional annotations and low log_10_ CP/MT ratios.

On the other hand, 346 genes were related to phototrophic eukaryotes (algae and plants), 20 to other eukaryotes, and 13 to prokaryotes, and were therefore considered gene transfer (GT) candidates (Fig. 4). We use this neutral designation to acknowledge the fact that the direction of the transfer usually cannot be objectively established. In case a sequence was related to a clan composed of more than one higher algal taxon, it was assigned to the category “miscellaneous algae” or one of its narrower sub-categories (supplementary-dataset-1.xlsx). It should be noted that a fraction of horizontal relationships may result from phylogenetic artefacts, taxon sampling bias or electronic contamination of the datasets composing the in-house database.

**Fig. 4:**
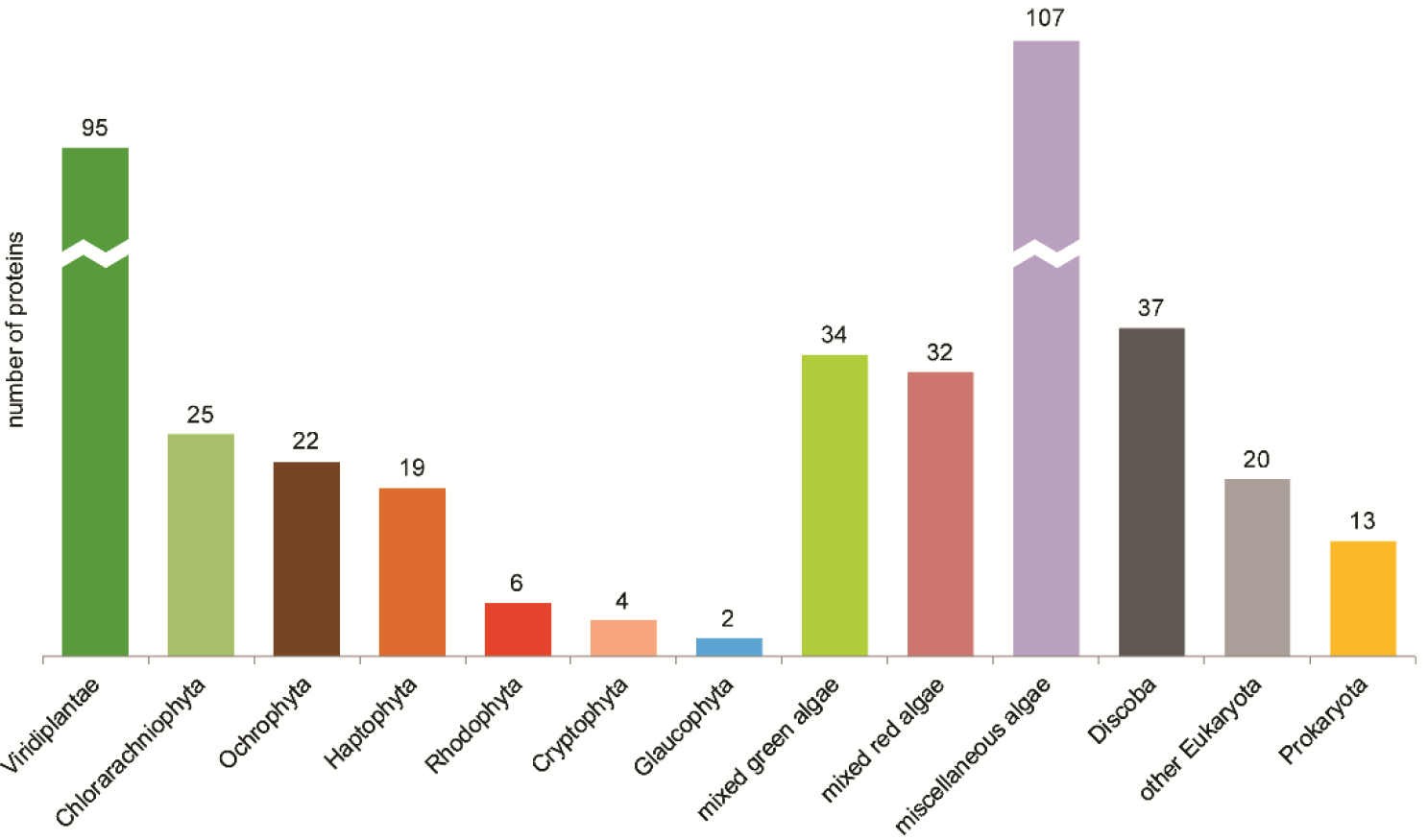
Evolutionary affiliations of 416 nuclear-encoded plastid proteins presumably representing lateral gene transfer into or from *E. gracilis*. Protein trees are available at https://drive.google.com/drive/folders/13zdxl2CjdXhzB-OgrXqIJdq7YaQ1V9h2.

The majority of GT candidates are related to green algae/plants (95) and most likely can be explained as the result of endosymbiotic gene transfer accompanying the origin of the euglenid plastid. The affiliation of the remaining GT candidates was more or less equally distributed among three groups of secondary algae: chlorarachniophytes, ochrophytes, and haptophytes (25, 22, and 19 genes, respectively; Fig. 4), with red algal, cryptophyte, and glaucophyte affiliations predicted for six, four and two genes, respectively. Some genes were related to a mixed group of phototrophs. Out of these, 34 proteins were related to mixed green lineage algae while 32 proteins were related to mixed red algae lineages, 23 of which were predicted as related specifically to secondary plastid-bearing organisms.

Some of the 13 proteins related to prokaryotes provide interesting new features to the organelle. For example, this set contains two components (SufS and SufB) of bacterial SUF system of FeS cluster assembly related to Chlamydiae. Manual search recovered presence of two other components (SufC and SufD) present in the proteome and additional one (SufE) present in the transcriptome with similar affiliation. There proteins form together a second set of enzymes for FeS cluster assembly next to vertically inherited green algae-related SUF system homologues, which are also present (Fig. 5, Table S7 and Fig. S8).

**Fig. 5:**
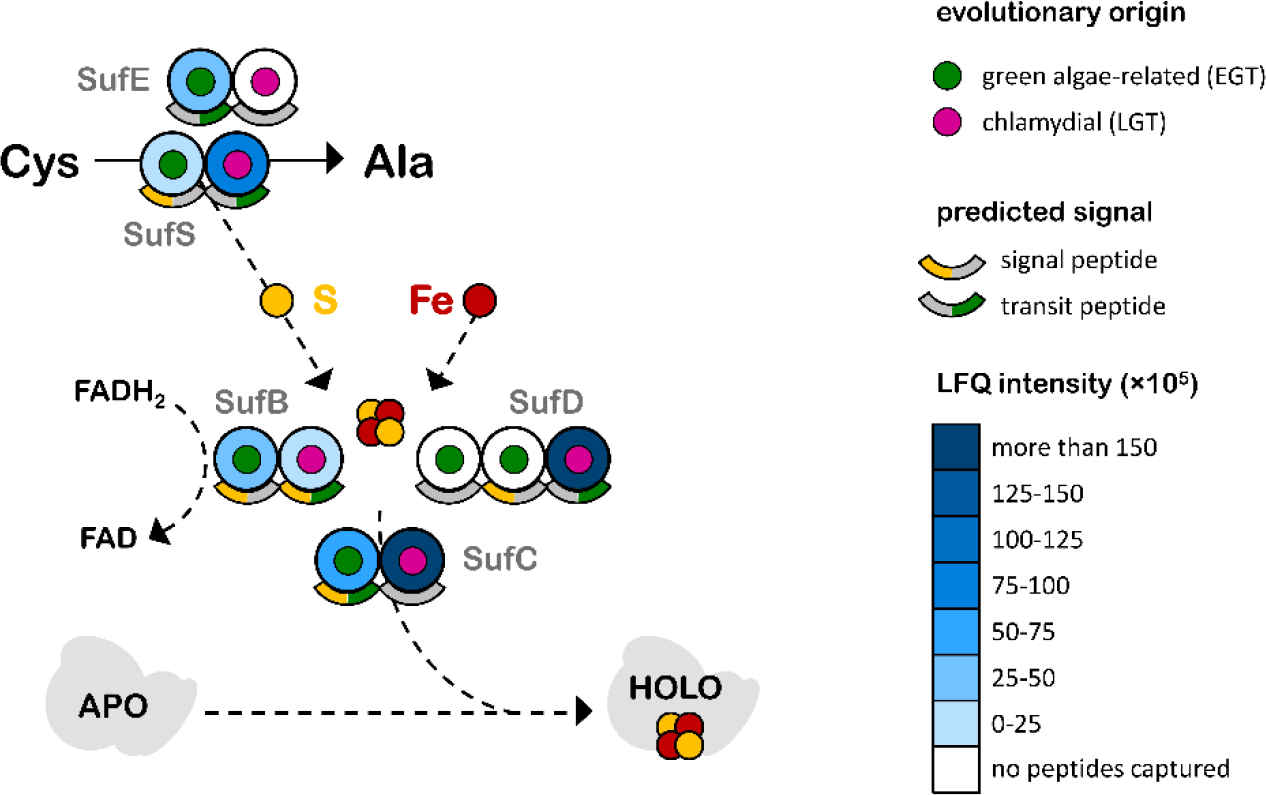
Plastidal SUF pathway of *E. gracilis*. Multiple copies of each subunit are represented by circles colored in shades of blue based on their average LFQ intensity per replicate which indirectly corresponds to the relative protein abundance. Proteins captured in only one replicate are also included. The evolutionary origin of the particular subunit copy is represented by the dot in the middle: green for green algae-related, i.e. gained along with the plastid via endosymbiotic gene transfer (EGT), or magenta for chlamydial-like, i.e. gained via LGT. The presence of predicted signal and transit peptides is also indicated.

### Re-evaluation of the N-terminal plastid-targeting presequences *in E. gracilis*

We took advantage of the large set of confidently identified plastid-targeted proteins to re-evaluate the characteristics of *E. gracilis* plastid-targeting presequences as the major reorganization of protein importing machinery may imply notable changes in these signals. The N-terminal sequences of 375 preproteins with well-supported plastid localization (log_10_ CP/MT ratio > 1 or a photosynthetic function) and non-truncated N-termini (verified by the presence of the spliced leader) were analysed. The general presence of one or two hydrophobic domains within the N-termini of plastid-targeted proteins was confirmed with positions estimated between residues 9-60 for the signal peptide (SP) and 97-130 for the stop-transfer signal (STS) using Kyte-Doolittle hydrophobicity score calculated for each position (Fig. 6). The sequences were sorted into previously defined classes based on the presence of these hydrophobic domains (Durnford and Gray 2006): 47% were of class I (containing both SP and STS), 37% of class II (containing SP only), with the surprising 16% exhibiting neither of these hydrophobic domains and therefore referred to as “unclassified”. The proportion of “unclassified” preproteins is relatively high and suggests that there is a significant cohort of plastid proteins with non-conventional targeting that elude straightforward *in silico* identification. The mechanism of their targeting to the plastid remains elusive, they either use an alternative signal, piggyback on class I and II preproteins, and/or are targeted to a specific plastid sub-compartment. However, when possible bias in protein functions associated with different preprotein classes was investigated using a pivot table and chi-squared test, no statistically significant pattern was observed.

**Fig. 6:**
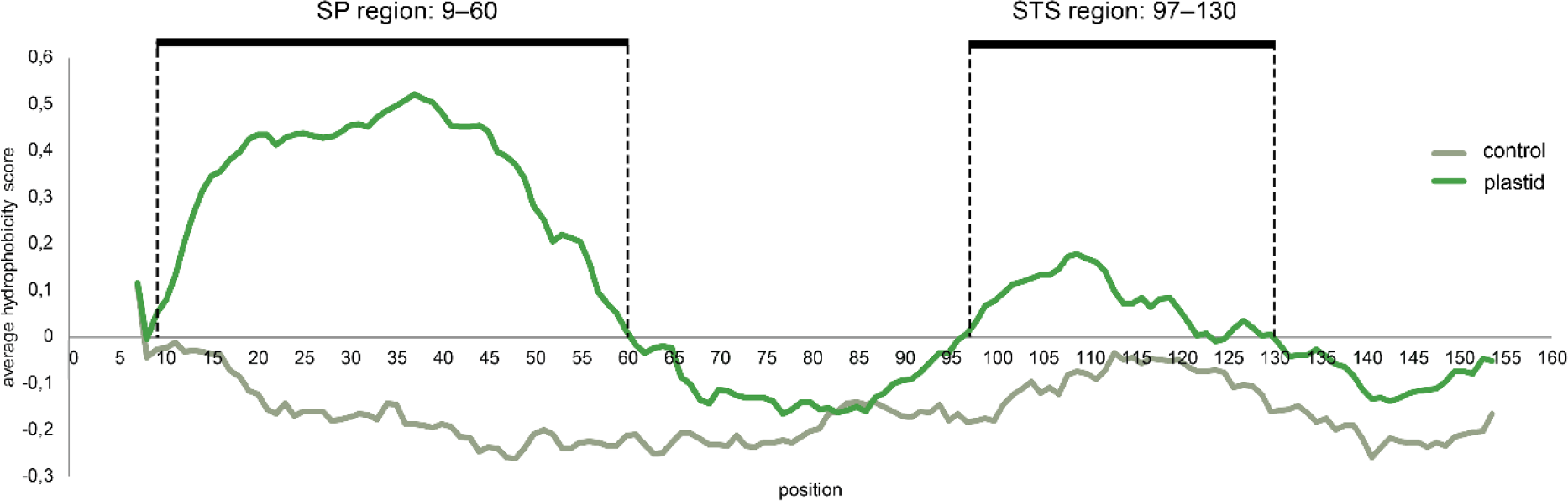
The two hydrophobic domains in the N-termini of 375 well-supported plastid-targeted proteins with complete N-terminus as seen on the graph of average hydrophobicity score per position compared to the negative control (375 randomly selected proteins of *E. gracilis*). The range of the SP was estimated as positions 9-60 (note, that the first and last six positions of the 160 aa long sequence are missing as a result of window-based scoring; position 7 is the first position with score assigned), the range of the STS was estimated as positions 97-130. The hydrophobic peak representing the STS is clearly less accentuated than the one representing SP reflecting the fact that it is present in just a subset of plastid-targeted proteins termed class II pre-proteins.

The canonical euglenophyte plastid-targeting sequence includes a region directly downstream of the SP that exhibits features of the plastid transit peptide (TP) and can mediate plastid protein import in the heterologous plant system, indicating that it is recognised by the plant TOC as an authentic plant TP (Sláviková et al. 2005). The presence of the TP-like (TPL) region in euglenophyte presequences may thus look surprising, given the apparent absence of the TOC complex in these organisms. To get a better understanding of the function of the TPL region in plastid protein targeting in euglenophytes, we investigated its sequence characteristics more closely. The putative TPL region and the mature protein region were selected from the 375 plastid proteins, either based on the actual positions of the hydrophobic domains in each protein, or estimated from the overall hydrophobicity profile of all the studied N-termini. Their amino acid compositions differed significantly from the mature protein region (Fig. 7) as opposed to the control sets of 375 randomly selected proteins, where the only significant difference recorded was the relative enrichment in serine (for full results for both sets and all classes of preproteins, see Tables S12 and bar plots S13).

**Fig. 7:**
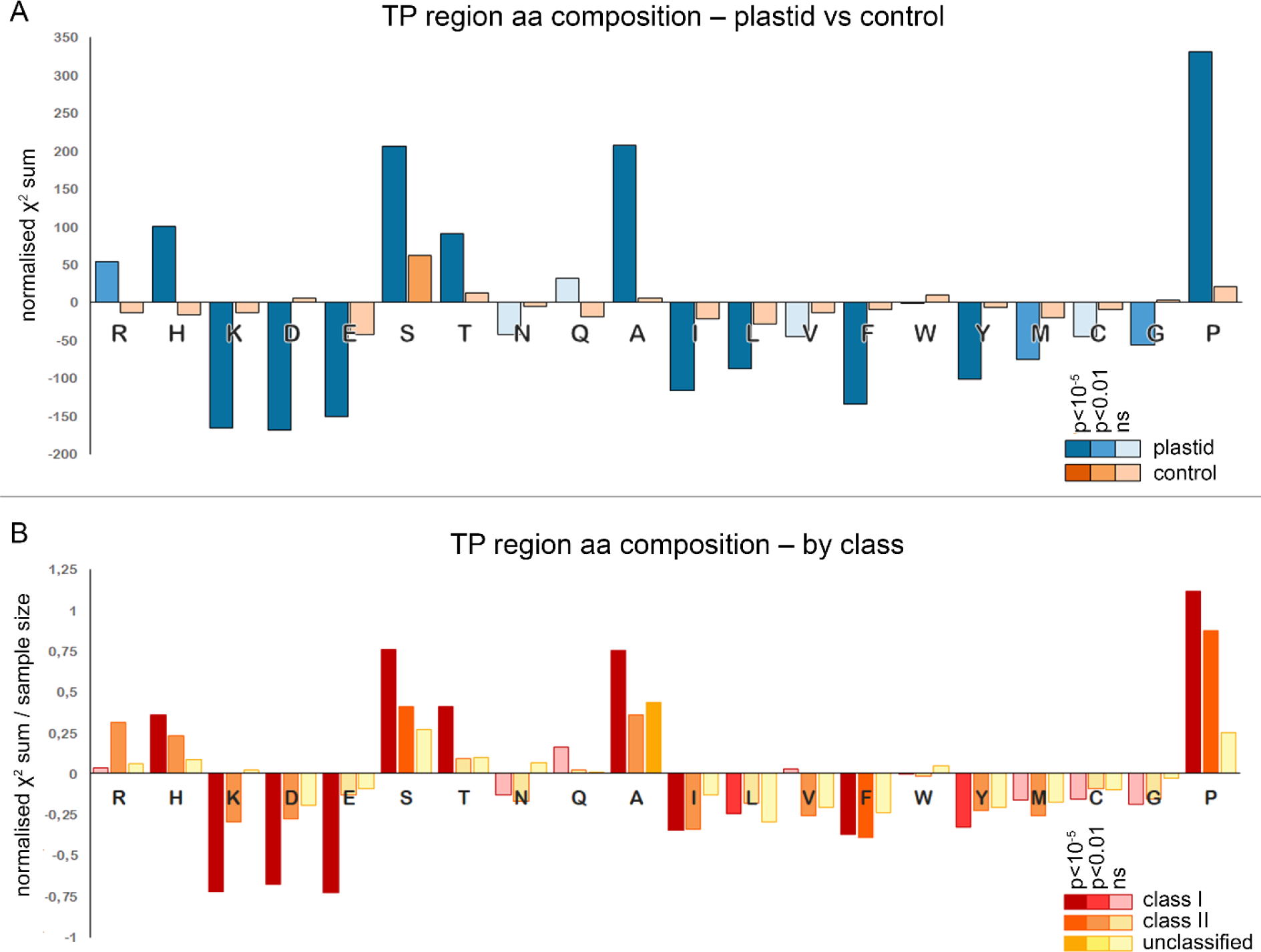
Statistical comparison of the amino acid frequencies in the putative transit peptide region and the putative mature chain of the same protein for the set of 375 proteins regardless of their classification for the plastid protein sample in comparison to the negative control (shades of blue and orange, respectively, A). The same comparison was also performed separately for the plastid proteins of class I, class II, and “unclassified” (shades of red, orange, and yellow, respectively, B). The vertical axis represents normalized χ^2^ sum reflecting whether the amino acid frequency is higher or lower than expected (positive or negative values) and the relative degree of its enrichment or depletion. The medium coloured bars represent statistically significant results with *p* < 0.01, the dark coloured bars represent those with *p* < 10^−5^.

## Discussion

### Quality assessment and comparisons to other sets of plastid proteomes

To assess the quality and completeness of our proteomic dataset, we compared it with other mass spectrometry-based plastid proteomes and predicted datasets of plastid-targeted proteins.

In comparison to the mass spectrometry-determined plastid proteomes of *Arabidopsis thaliana* (Huang et al. 2013), *Chlamydomonas reinhardtii* (Terashima et al. 2011), and *Bigelowiella natans* (Hopkins et al. 2012), the *E. gracilis* plastid proteome is relatively large and contains a higher portion of proteins lacking functional assignment and/or readily identifiable homologs (42.5% vs 15% in *A. thaliana*, 25% in *C. reinhardtii* and 25% in *B. natans*). While this may be partially due to sampling bias, the proportion of proteins sorted into equivalent functional categories clearly differs (Table 1). Notable differences are present in the very low number of assignable proteins involved in amino acid metabolism and also in the relatively small proportion of photosynthetic proteins, which might reflect metabolic versatility of *Euglena*, with phototrophy being non-essential and readily “switched off” in dark and/or anaerobic conditions. In addition to this metabolic plasticity, the higher structural complexity of the secondary plastid with additional membrane and inter-membrane compartments is also likely to contribute to the relatively large size of the proteome.

**Tab. 1:**
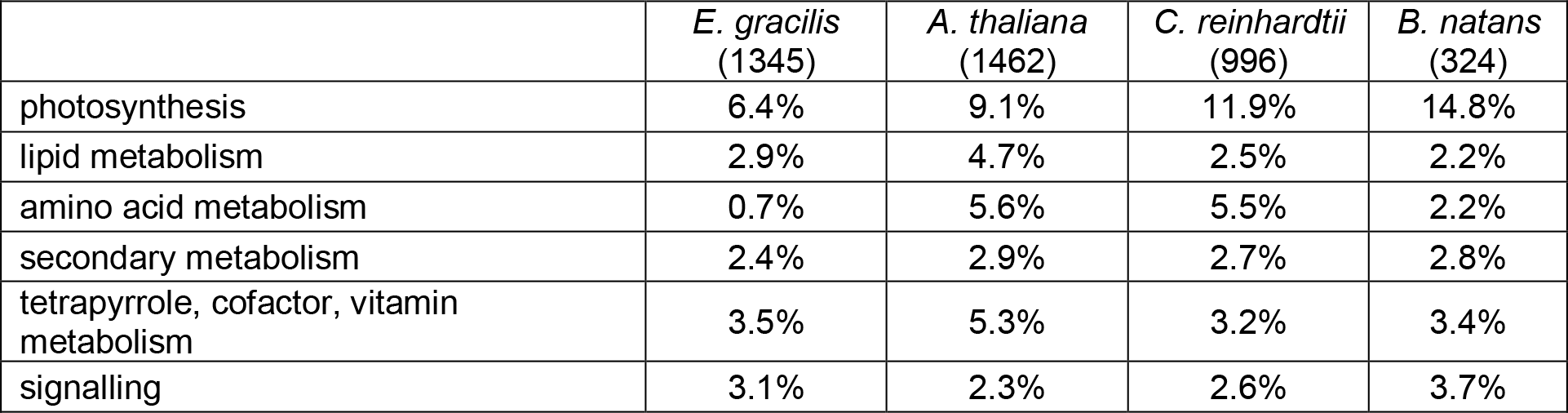
Comparison of the proportions of proteins of selected functional categories in the four plastid proteomes; the number in brackets under the respective organism names is the total number of identified plastid proteins. The plastid proteome of *Euglena* differs from the other three in the very low amount of amino acid metabolism proteins and in the relatively small proportion of photosynthetic proteins.

|  | *E. gracilis*<br>(1345) | *A. thaliana*<br>(1462) | *C. reinhardtii*<br>(996) | *B. natans*<br>(324) |
| --- | --- | --- | --- | --- |
| photosynthesis | 6.4% | 9.1% | 11.9% | 14.8% |
| lipid metabolism | 2.9% | 4.7% | 2.5% | 2.2% |
| amino acid metabolism | 0.7% | 5.6% | 5.5% | 2.2% |
| secondary metabolism | 2.4% | 2.9% | 2.7% | 2.8% |
| tetrapyrrole, cofactor, vitamin metabolism | 3.5% | 5.3% | 3.2% | 3.4% |
| signalling | 3.1% | 2.3% | 2.6% | 3.7% |

The presence of contaminant proteins in the organellar fractions is inevitable, as are failed protein detections. The purity of the plastid fraction assessed by relative distribution of marker proteins revealed moderate contamination by other cell compartments (see Material and methods and Figs. S1 and S2.1-3). The failed protein detections may reach up to 30% as indicated by comparisons to the set of plastid genome-encoded proteins and sets of high-confidence plastid proteins predicted by Durnford and Gray (2006) and Záhonová et al. (2018). These stem from loss of proteins/peptides in the mass spectrometry pipeline or limitations in detecting low abundance and small proteins. Besides technical issues, biological factors may be responsible for the failure to identify some of the *bona fide* plastid proteins. Dual protein localization is a common phenomenon reported for aminoacyl-tRNA synthetases (Duchêne et al. 2005; Gile et al. 2015) as well as other proteins (Carrie and Small 2013), and it may interfere with assignment of protein to a particular organellar proteome. Indeed, we identified only 18 aminoacyl-tRNA synthetases (including tyrosyl- and methionyl-tRNA synthetase fused in a single protein but missing leucyl- and prolyl-tRNA synthetases), despite the fact that tRNAs for all 20 amino acids are encoded in the plastid genome (Hallick et al. 1993). The dual localization candidates are listed in the respective section of supplementary-dataset-1.xlsx.

These analyses indicate that our plastid proteome is, to some extent, incomplete, and moderately contaminated with other cellular fractions. Still it represents the most comprehensive estimate of *E. gracilis* plastid protein content available and is a solid basis for analyses presented in this study.

### *Euglena* plastid proteome is of mixed evolutionary origin

Although it is clear that the euglenid plastid originated from the pyramimonadalean green alga (Turmel et al. 2009; Jackson et al. 2018), high proportion of plastid proteins show phylogenetic relationship to other algal groups. This fact was noted previously in studies based on small cohorts of *E. gracilis* genes (Maruyama et al. 2011; Markunas and Triemer 2016; Lakey and Triemer 2017; Ponce-Toledo et al. 2018) and it was suggested that these genes were acquired from a “chromophyte” prey or symbiont by the common ancestor of eukaryovorous and phototrophic euglenids. Our results from a fuller set of experimentally determined plastid proteins support this hypothesis and allow semiquantitative evaluation of the contributions from phototrophic eukaryotes. The number of genes sorted into Rhodophyta (6), Cryptophyta (4) and Glaucophyta (2) category is very low, and arguably at the level of false positive determinations from the phylogenetic pipeline. On the other hand, the number of genes related to chlorarachniophytes (25), ochrophytes (22) and haptophytes (19) are of sufficient size and there presence may reflect some events in the evolutionary past of the group.

These observed affiliations could be explained by: (1) gene transfers into the eukaryovorous ancestor of euglenids, *sensu* the “you are what you eat” hypothesis (Doolittle 1998), (2) gene transfers in the initial stage of endosymbiont integration when the euglenid host was presumably obligatory mixotrophic, much like the extant *Rapaza viridis* (Yamaguchi et al. 2012), such transfers could have compensated for the reductive evolution of the endosymbiont genome, as it was proposed for the chromatophore of *Paulinella* (Marin et al. 2005; Nowack et al. 2016), (3) gene transfers from a cryptic endosymbiont putatively present in the euglenophyte ancestor and replaced by the extant organelle, similar to the cases of serial endosymbioses and overall plastid fluidity in dinoflagellates (Saldarriaga et al. 2001; Yoon et al. 2005; Takano et al. 2008; Matsumoto et al. 2011; Xia et al. 2013; Burki et al. 2014; Dorrell and Howe 2015); and finally (4) by gene transfers from euglenids to the other algal group.

In the case of *E. gracilis* plastid proteins affiliated to chlorarachniophyte homologs, the direction of transfer from euglenophytes to chlorarachniophytes seem more likely, given the fact that the latter group was estimated as > 200 MY younger than the former (Jackson et al. 2018). In some cases, the determined affiliation to chlorarachniophytes may also result from undersampling of green algae in the pipeline, since both euglenids and chlorarachniophytes acquired their plastids from chlorophytes. In the case of ochrophyte and haptophyte related genes we have no reason to prefer from-euglenid direction of the transfer and these algae might indeed be important contributors to the euglenophyte plastid proteome.

The predicted functions of proteins affiliated to non-green algae vary and their distribution between particular metabolic pathways and multisubunit complexes is rather patchy (Fig. 2). This contrasts with situations where a cluster of functionally related proteins, such as enzymes catalyzing consecutive steps of a metabolic pathway, is transferred laterally. The latter is typical for prokaryote-to-eukaryote gene transfers (Keeling and Palmer 2008) and one such case will be discussed further below.

Presence of non-green algal related genes in the plastid proteome is congruent with previous studies focused on smaller sets of *Euglena* proteins. Markunas and Triemer (2016) focused on the phylogenetic origin of the Calvin cycle enzymes. Our phylogenetic pipeline used only sequences from *E. gracilis* and not from other euglenids and was stricter in relationship assignment than theirs. As a result, some of the enzymes reported by Markunas and Triemer (2016) as related to algae of the red plastid lineage (ribulose-1,5-bisphosphatase small chain, triosephosphate isomerase, phosphoribulokinase, two of the three plastid forms of fructose 1,6-bisphosphatase) were rejected here due to poorly supported topology or assignment to a broader or different category. We also compared our results to the phylogenetic-origin assignments of *E. gracilis* proteins reported by Ponce-Toledo at al. (2018). Out of 42 sequences analysed in both studies, 23 were consistently evaluated as affiliated to “green” or “red” lineages, although often with BS < 75%, four could not be assigned to a specific algal group, 12 were not determined as algal affiliated and only for three proteins were predicted algal affiliations in conflict (i.e. red vs. green), but with low BS (supplementary-dataset-2.xlsx). Both comparisons demonstrate that sensitivity of phylogenetic origin determination might vary among studies as a result of different research aims and tested hypotheses and subsequent taxon sampling, tree interpretation, and other methodical approaches, and the studies are not always comparable in the strict sense. At the same time, we believe that within a single study, in which phylogenetic estimates are performed under strictly defined and identical conditions for all genes, such as the study presented in this paper, the sizes and functionalities of phylogenetic gene groups are comparable and carry interesting information about the mixed origin of the plastid proteome.

The presence of proteins of prokaryotic affiliation is also noteworthy, especially since many of these point at the Chlamydiae as a donor group. A surprisingly high number of chlamydial-like proteins is present in primary plastids of plants and algae (Becker et al. 2008; Moustafa et al. 2008), which led to the “ménage à trois” hypothesis proposing that a chlamydial endosymbiont was directly implicated in the integration of the co-occurring cyanobacterial endosymbiont that would later become a plastid (Facchinelli et al. 2013; Cenci et al. 2016). Whether this could also be the case of secondary plastid establishments remains to be investigated. The chlamydiae-related plastid proteins in euglenophytes (some of which are also functionally significant and will be discussed further) are not shared by other eukaryotic lineages, which suggests these are not ancestral (inherited with the plastid) but instead represent novel, independent acquisitions. Could this mean that a chlamydial endosymbiont (or intracellular parasite) inhabiting the common ancestor of euglenophytes has played a similarly important role in the establishment of the secondary plastid of this algal group? The “ménage à trois” hypothesis and even the purported chlamydial origin of the critical genes have been contested, however (Deschamps 2014; Domman et al. 2015), so it is possible that the presence of these genes in euglenophytes (and maybe even plastids in general) is simply a result of chlamydiae, a group of obligate intracellular parasites or commensals (Subtil et al. 2014), being particularly prone to donating their genes to eukaryotes.

### Metabolic functions of the *Euglena* plastid: commonalities and deviations

We reconstructed the major metabolic pathways of the *Euglena* plastid from the proteomic and transcriptomic data (Fig. 2). While some proteins of the reconstructed pathway were not found in the proteome, it is likely that many such gaps are due to failure to detect respective peptides, as frequently these are represented in the transcriptome. Overall, the plastid metabolic functions predicted here agree with previous reconstructions based solely on *in silico* evidence (Ebenezer et al. 2019) and include most commonly present plastid pathways. However, both the overall pathway complement and several individual pathways do exhibit features indicative of considerable novelty and these will be discussed in details below.

As expected, the photosynthetic apparatus is well represented. In accordance with previous findings (Wildner and Hauska 1974), no gene or protein for plastocyanin was identified (white circle in e^-^ transport section of Fig. 2, Fig. S5.2), so electron transfer between the cytochrome *b*_6_*f* complex and the photosystem I relies solely on two different isoforms of cytochrome *c*_6_. Interestingly, *E. gracilis* also encodes a homolog of cytochrome *c*_6A_ (seqid 17930), previously only known from land plants and some green algae (Howe et al. 2006). Cytochrome *c*_6A_ is thought to function as a redox-sensing regulator within the thylakoid lumen and the structure of its N-terminus in *E. gracilis* is in accord with this localization. This protein was not detected in the plastid proteome, which is not unexpected given its low abundance and difficulty to detect in plants (Howe et al., 2006). Notable is also the presence of two homologs of the alpha subunit of the F-type ATPase in the proteome, a canonical copy encoded by the plastid genome and a divergent copy encoded in the nucleus (seqid 3018). Finally, the *E. gracilis* plastid contains two different terminal oxidases (seqids 2887 and 18315), i.e. components of the chlororespiration pathway mediating reoxidation of reduced plastoquinone by O_2_ (Nawrocki et al. 2015). One represents the conventional enzyme (acquired from a dinoflagellate source), whereas the other falls among mitochondrial alternative oxidases (Fig. S6). The latter is apparently not a contaminant from the mitochondrion, as it is significantly enriched in the plastid fraction and has a predicted plastid-targeting signal, while at least three different isoforms of the mitochondrial alternative oxidase bearing predicted mitochondrial transit peptides are present in *E. gracilis*. Why this terminal oxidase was recruited into *Euglena* plastid remains to be investigated.

The Calvin cycle is fully represented, but the route to glucose metabolism is apparently missing due to the lack of glucose-6-phosphate isomerase. This is probably related to the loss of starch synthesis during the emergence of the euglenophyte secondary plastid and the switch to extraplastidial paramylon (β-1,3-glucan) as the main storage compound (Barsanti et al. 2001). The key enzyme of paramylon synthesis, *E. gracilis* glucan synthase‐like 2 (EgGSL2; Tanaka et al., 2017), is indeed absent from the plastid proteome, but its paralog, EgGSL1 (seqid 4050), was identified as a high-confidence plastid protein. Unfortunately, its substrate specificity and function are unknown. The nearly complete representation in the *E. gracilis* plastid proteome of the C5 pathway of tetrapyrrole (i.e. haem and chlorophyll) synthesis, using glutamate as the precursor of the key intermediate aminolevulinate, is in accord with previous evidence (Kořený and Oborník 2011; Lakey and Triemer 2017), while the enzymes of the parallel C4 (Shemin) pathway, utilizing glycine and succinate as aminolevulinate precursors, are absent, in agreement with predicted mitochondrial/cytosolic localization (Kořený and Oborník 2011).

The *E. gracilis* plastid is also a site for biosynthesis of fatty acids and glycerolipids. The latter include the major plastid phospholipid phosphatidylglycerol and three glycolipids, monogalactosyl- and digalactosyl- diacylglycerol (MGDG and DGDG) and sulfoquinovosyl-diacylglycerol (SQDG), in agreement with biochemical evidence documenting these lipids in the *E. longa* plastid (Matson et al. 1970; Blee and Schantz 1978; Shibata et al. 2018). We were able to reconstruct pathways for synthesizing these compounds, albeit with several enzymes missing from the proteome, but predicted by the transcriptome. Interestingly, our data suggests that *E. gracilis* lacks one of the key enzymes of SQDG synthesis, UDP-sulfoquinovose synthase (SQD1), which catalyzes formation of UDP-sulfoquinovose from UDP-glucose. This is not due to the potential incompleteness of our data, as no SQD1 candidates were identified in transcriptome assemblies available for other euglenophytes. However, it was demonstrated that sulfite, one of the SQD1 substrates, is readily incorporated into SQDG when incubated with isolated *E. gracilis* plastids (Saidha and Schiff 1989), suggesting the presence of an alternative (SQD1-unrelated) form of UDP-sulfoquinovose synthase. In any case, UDP-sulfoquinovose is produced in *E. gracilis*, with the conventional sulfoquinovosyltransferase (SQD2) catalyzing the ultimate step of SQDG formation (Fig. 2).

Plastids are generally the site of production of various amino acids, some of which (such as phenylalanine, tyrosine, and tryptophan) are produced exclusively in the organelle in algal and plant cells (Rippert et al. 2009; Maeda and Dudareva 2012; Reyes-Prieto and Moustafa 2012). In contrast, our analysis shows that the role of plastids in amino acid metabolism is highly limited in *Euglena*. Only an incomplete serine synthesis pathway could be reconstructed, with no identifiable plastid-targeted phosphoserine aminotransferase. In addition, only phosphoserine phosphatase could be identified in the proteome, whereas the plastid localization of the enzyme catalyzing the initial step of the pathway (phosphoglycerate dehydrogenase, divided into two contigs, sequids 23072 and 17279) remains an *in silico* prediction. The *E. gracilis* plastid proteome does contain an isoform of cysteine synthase A, but a plastidial version of the enzyme that converts serine to O-acetylserine, the substrate of cysteine synthase A, was not detected, suggesting that O-acetylserine needs to be imported into the organelle. The plastid further harbours serine/glycine hydroxymethyltransferase (SHMT), whose primary role likely is to provide a formyl group for the synthesis of formylmethionyl-tRNA required for translation initiation in the plastid rather than to make glycine from serine (Fig. S5.1). There was no evidence for other amino acid biosynthetic pathways within the *E. gracilis* plastid.

### Terpenoid biosynthesis in the *Euglena* plastid

Two different biochemical routes are responsible for the synthesis of isopentenyl pyrophosphate (IPP), a precursor of steroids and terpenoids, in eukaryotes: the MVA pathway, localized to the mitochondria/cytosol, and the MEP (DOXP) pathway, always compartmentalized into the plastid (Zhao et al. 2013). *E. gracilis* was originally believed to harbour only the MVA pathway (Disch et al. 1998; Lange et al. 2000; Watanabe et al. 2017), which was later disproved by biochemical evidence for a role of the MEP pathway in the synthesis of carotenoids (Kim et al. 2004) and identification of homologs of pathway enzymes in the transcriptome data (O’Neill et al. 2015). Our proteomic data add the final piece of information on this pathway in *E. gracilis* by demonstrating plastid localization. In the proteome we also identified the enzyme for the synthesis of the carotenoid precursor geranylgeranyl pyrophosphate (GGPP) from IPP, and most enzymes required for the synthesis of carotenoids previously reported in *Euglena* biochemically (Krinsky and Goldsmith 1960): antheraxanthin, neoxanthin, β-, γ- and ζ-carotene, retinol, lutein and cryptoxanthin. Our proteomic and transcriptomic data, however, fail to provide evidence for β-carotene ketolase synthesizing two of the carotenoids determined previously, echinenone and canthaxanthin (Krinsky and Goldsmith 1960). Still, some of the enzymes of these pathways were found in the transcriptome (marked as grey circles in Fig. 2).

Furthermore, the proteome suggests that IPP, made by the MEP pathway, is used for the synthesis of plastoquinone (via solanesyl-PP) in the *E. gracilis* plastid, as is usual in plastids in general (Lohr et al. 2012). In contrast, biochemical investigations revealed that, uniquely among plastid-bearing eukaryotes, the MEP pathway does not provide IPP for synthesis of phytyl-PP, a precursor of tocopherols (vitamin E), phylloquinone (vitamin K), and chlorophyll, in *E. gracilis* (Kim et al. 2004). Our analyses support this finding and provide an insight into the alternative pathway. Phytyl-PP is conventionally made in the plastid by the activity of ChlP, a GGPP reductase conserved across both eukaryotic and prokaryotic phototrophic organisms (Heyes & Neil Hunter, 2009). The *E. gracilis* plastid proteome lacks this enzyme and no ChlP homolog is detectable in the transcriptome, explaining why GGPP formed in the plastid by the MEP pathway cannot serve as a phytyl-PP precursor in this alga. An alternative route to phytyl-PP operates in plant chloroplasts, which recycles phytol released during chlorophyll degradation and significantly contributes to tocopherol biosynthesis (Vom Dorp et al. 2015). This pathway consists of two enzymes: VTE5 (phytol kinase, seqid 13197) and VTE6 (phytyl phosphate kinase, seqid 14381), both of which are present in the plastid proteome. The proteome also contains a homolog of a recently characterized chlorophyll dephytylase (CLD1, seqid 12254; Fig. S5.5; Lin et al., 2016), which further suggests that, like in plants, phytol is salvaged from degraded chlorophyll in *E. gracilis*. This obviously cannot be the only means of producing phytyl-PP in *E. gracilis*, as the compound itself is required for chlorophyll synthesis. Interestingly, the non-photosynthetic chlorophyll-lacking *E. longa* retains both VTE5 and VTE6 (GGOE01004182.1 and GGOE01053790.1; Záhonová et al. 2018), and suggests that phytol is a genuine intermediate of *de novo* phytyl-PP in *Euglena*. As direct phytol synthesis has not been characterized in any organism, to our knowledge, how and in which cellular compartment phytol is made in *Euglena* is thus unclear.

*E. gracilis* is well known to produce tocopherol (Watanabe et al. 2017) and homologs of some enzymes responsible for its synthesis were previously described in the transcriptome (O’Neill et al. 2015). There has been some controversy about the subcellular localization of tocopherol (vitamin E) production in *E. gracilis* (Watanabe et al. 2017), which we now resolve by reconstructing the full pathway, from the precursors homogentisate and phytyl-PP to α-tocopherol, and identification of all four enzymes in the plastid proteome (Fig. 2). Another terpenoid-quinone produced by *E. gracilis* is phylloquinone, or more precisely its derivative 5’-monohydroxyphylloquinone (Ziegler et al. 1989). In plants and green algae most steps of phylloquinone synthesis occur in the plastid, except three reactions from o-succinylbenzoate to dihydroxynaphthoate, which are catalyzed by peroxisomal enzymes (Emonds-Alt et al. 2017; Cenci et al. 2018). *E. gracilis* encodes a homolog of the multifunctional protein PHYLLO catalyzing all four steps of the first part of the pathway (seqid 121), but the protein is absent from the plastid proteome and lacks a targeting presequence, consistent with earlier biochemical evidence placing synthesis of o-succinylbenzoate in the cytosol (Seeger and Bentley 1991). The same study tentatively associated the o-succinylbenzoate to dihydroxynaphthoate part of the pathway to the plastid envelope rather than peroxisomes, but the respective enzymes must be unrelated to the peroxisomal enzymes MenE, MenB, and MenI, as their homologs were not detected in either the plastid proteome or transcriptome. Genome analyses indicate the existence of an unknown pathway of dihydroxynaphthoate synthesis in some algae (e.g. glaucophytes) and cyanobacteria (Cenci et al., 2018), so it is possible that *E. gracilis* converts o-succinylbenzoate to dihydroxynaphthoate by employing enzymes of this uncharacterized route. The final part of the pathway, starting with phytylation of dihydroxynaphthoate, is predicted to be plastid-localized in *E. gracilis* (Fig. S5.5), although only one of the respective enzymes (MenA, seqid 7550) was identified in our proteomic analysis and the identity of the enzyme catalyzing the presumably final step (phylloquinone hydroxylation) is unknown.

### Surprising gifts from chlamydiae: an extra SUF system and sulfite reductase

The SUF pathway is essential and universally present in plastids, supplying multiple core enzymes with iron-sulfur (FeS) clusters (Lu 2018). Beyond plastid-bearing eukaryotes, the SUF system or its components are present in some anaerobic protist lineages as a result of LGT from Archaebacteria (Tsaousis et al. 2012; Stairs et al. 2014; Leger et al. 2016) or Proteobacteria (Karnkowska et al. 2016). Curiously, our data suggest that euglenophytes also underwent lateral acquisition of bacterial SUF. These genes are most closely related to

Chlamydiales and instead of compensating for a loss of other FeS cluster assembly machinery, they localize to the plastid along with the original SUF inherited via the green algal ancestor (Fig. 4). This apparent redundancy does not represent a transient state and/or result of neutral evolution because the two parallel SUF systems are also present in *E. longa* and the distantly related *Eutreptiella* spp. (sources and accession numbers in Table S7, phylogenetic trees in Fig. S8). The relative protein abundance, estimated by LFQ intensity, suggests that chlamydial Suf proteins are generally more abundant than the algal cohort, which suggests that the bacterially derived pathway is likely active and contributes towards plastid iron-sulphur metabolism (Fig. 4).

Co-occurrence of two SUF pathways in euglenophyte plastids may indicate functional specialization. One possibility is that they mediate FeS cluster assembly for different client proteins in the same compartment which, to our knowledge, would be unprecedented in eukaryotes. An alternative explanation, i.e. distinct plastid sub-compartmentalization we suggest is more likely, for example with one in the stroma in common with other plastids (Lu 2018), and the second restricted to one of the intermembrane spaces, serving the co-localized FeS proteins. A further novel aspect of FeS cluster assembly machinery in *E. gracilis* is the presence of an ABC transporter (seqid 3116) in the plastid proteome, which is homologous to mitochondrial Atm1 protein required for the export of unspecified FeS cluster intermediate from mitochondrion to the cytosol. Presence of this transporter suggests that such intermediates may be transported across a membrane, either into or outside the plastid or between its sub-compartments. For the detailed results of a direct search of FeS cluster assembly associated proteins, see supplementary-dataset-1.xlsx

While direct experimental data are necessary to understand this FeS biosynthetic novelty, some indication may be present amongst the other genes of chlamydial origin. We found two such proteins, both with orthologs in *E. longa* and *Eutreptiella*. One of them is an isoform of ferredoxin (divided into two contigs, seqids 37420 and 61063 in our data, seqid 74687 in Yoshida et al. (2016), supplementary-dataset-2.xlsx; Fig. S9), itself a FeS protein, which may represent a client of the chlamydial SUF, and the second is the alpha subunit of NADPH-dependent sulfite reductase (CysJ). Significantly, the beta subunit of this enzyme (CysI) contains a Fe_4_S_4_ cluster (Smith and Stroupe 2012) and seems to be more related to spirochaetes (Fig. S10). The plastid localization of sulfite reductase is by itself notable, as previous biochemical studies associated sulfate assimilation, including the reaction catalyzed by sulfite reductase, with the mitochondrion in *E. gracilis* (Saidha et al. 1988). The plastid localization of sulfite reduction is consistent with the presence of cysteine synthase in this organelle, as this enzyme assimilates hydrogen sulfide produced in this reaction.

### New insights into the mechanism of plastid protein import in euglenophytes

The composition of the plastid protein import complexes TOC and TIC was investigated in detail. Surprisingly only two subunits were identified in the transcriptome (disregarding sequences highly divergent to Tic55 and Tic62): Tic32 and three isoforms of Tic21 (Záhonová et al. 2018). Proteomics confirmed the plastid localization of all Tic21 isoforms, but not Tic32 (Fig. 3). Tic21 cooperates with Tic20 to form a minor protein-conducting pore independent of the central channel in plant plastids (Teng et al. 2006; Kikuchi et al. 2009), so its assuming the main translocase function in a highly reduced complex is conceivable. Tic32, on the other hand, is a non-essential regulatory subunit with a versatile enzymatic activity (Balsera et al. 2010) that could readily serve some TIC-independent purpose. The recently characterized components of the plastid import machinery conserved in plants and green algae, namely Tic236 (Chen et al. 2018) and a heteromeric AAA-ATPase complex consisting of Ycf2 and FtsH-like proteins (Kikuchi et al. 2018), are likewise not discernible in the proteome or transcriptome (Záhonová et al., 2018), providing additional support for the notion that the plastid protein import mechanism in euglenophytes differs profoundly from that of other eukaryotes.

Two proteins (seqids 13308 and 11030), representing a novel subgroup of rhomboid-related pseudoproteases conserved in other euglenophytes and similar to derlins known from the ERAD pathway were also present (Fig. S11, supplementary fasta alignment file derlins.txt). This machinery was partially duplicated and recruited for protein transport in the plastids of “chromalveolates” to form a system termed SELMA (Maier et al. 2015), with two derlin family proteins (Der1-1 and Der1-2) presumably constituting the protein-conducting channel. It is uncertain whether the two aforementioned proteins could serve the same function in the euglenophyte plastid, and euglenophytes appear to lack homologs of other SELMA components. Nevertheless, we believe they are noteworthy candidates for components of the unknown euglenophyte plastid protein import system, specifically as potential translocases of the middle plastid membrane (Fig. 3), filling a gap in the protein import machinery caused by absence of TOC. If confirmed by functional experiments, this would be an extremely intriguing example of evolutionary convergence as well as an outrageous undermining of the assumed cyanobacterial origin of the middle membrane of the euglenophyte plastid. The potential loss of one of the two “primary” membranes seems very unlikely but it is probably not unprecedented, as demonstrated in the recently described stramenopile with only two plastidial membranes, one of which is clearly of host origin (Wetherbee et al. 2018).

Delivery of plastid-targeted proteins in euglenophytes involves fusion of Golgi-derived vesicles with the outermost plastid membrane. Multiple paralogs of certain membrane trafficking components are present in *E. gracilis*, and it was speculated that some may be involved in Golgi to plastid transport (Ebenezer et al. 2019). Interestingly, the plastid proteome contains homologs of coat complex subunits, several Rab GTPases and SNARE proteins, some of which cannot be dismissed as contaminants. For example, SNARE protein GOSR1 has two paralogs in *E. gracilis*, one of which is detected in the plastid proteome with log_10_ CP/MT ratio > 3 (seqid 18194), and the other (seqid 19152) was not captured at all. A second notable candidate is Rab5, a conserved Rab GTPase associated with the endosomal system (Langemeyer et al. 2018), again with convincing log_10_ CP/MT ratio > 3. These two proteins may play roles in targeting protein-transporting vesicles to the outermost plastid membrane (Fig. 3).

### Novel features of the plastid transit peptide may indicate novel receptors

The TPL region of plastid-targeted proteins exhibits several features which are in accordance with previous findings regarding general characteristics of plastid transit peptides (Bruce 2000; Patron and Waller 2007; Felsner et al. 2010; Li and Teng 2013) and also transit peptides of euglenophytes in particular (Durnford and Gray 2006). Specifically, they are enriched in hydroxy residues (S, T) and alanine (A) and depleted in acidic/negatively charged residues (D, E). The TPL is also significantly depleted in three highly hydrophobic residues, namely leucine, isoleucine, and phenylalanine (L, I, F), lysine (K), tyrosine (Y), methionine (M), and glycine (G), but enriched in arginine (R), histidine (H), and proline (P), the latter being the most significant, with *p* < 10^−5^. Overall net charge is slightly positive. The TPL was previously described as generally enriched in hydroxy residues, but our data suggests that this is the case only for the two polar ones, serine and threonine, while the non-polar hydroxy residue, tyrosine, is present in a significantly lower amount than in the mature chain. The enrichment in proline is unexpected and intriguing, and the secondary structure of the TPL might be greatly affected by the rigid structure of this amino acid and its propensity to forming turns and loops (MacArthur and Thornton 1991). When analysed separately for each class (class I, class II, and “unclassified”) the above-mentioned significant differences in amino acid composition can be seen in class I and class II, but not in the “unclassified” proteins (Fig. 7), suggesting that a detectable TPL is not present in the investigated region of “unclassified” preproteins.

The *E. gracilis* plastid TPL region is not conserved at the sequence level, but it has characteristic amino acid composition. In addition to the previously described positive net charge and enrichment in hydroxy residues and alanine, it is also significantly depleted in most non-polar residues and might form a distinctive secondary structure due to its high proline contents. It is possible that some of these additional characteristics or even a secondary or tertiary structure of the peptide is recognised by an as yet unknown receptor, consistent with the major reorganization of the plastid translocases discussed above as well as the state of knowledge regarding the secondary plastids of “chromalveolates” where TPL is recognized not only by TOC, but also by SELMA (Maier et al. 2015). The latter shows that recruitment of a novel receptor for pre-existing, only slightly, if at all modified targeting signal is possible. In case the two derlin-like proteins discussed in the previous section are in fact part of the plastid protein import pathway, this brings up the exciting question whether the TPL recognition by this potential SELMA-like system in *Euglena* could be provided by an evolutionarily convergent mechanism or even by the same (albeit yet uncharacterized) receptor.

## Materials and Methods

### Cell fractionation and protein sample preparation

The plastid fraction was isolated as described previously (Davis and Merrett 1973; Moreno-Sánchez et al. 2000; Dobáková et al. 2015). Cells of *E. gracilis* strain SAG 1224-5/15 were collected by centrifugation at 800 × g for 10 min and resuspended in SHE buffer (250 mM sucrose, 10 mM 4-(2-hydroxyethyl)-1-piperazineethanesulfonic acid (HEPES), 1 mM ethylenediaminetetraacetic acid (EDTA), pH 7.3) supplemented with 0.4% fatty acid-free bovine serum. All the following steps were performed on ice. The cells were disrupted by sonication at 80% power using a thick 19.5-mm probe (Ultrasonic homogenizer model 3,000; Biologics, Inc.). Sonication was performed in six 10-s pulses cycles with 2-min breaks between them. The sonicate was centrifuged for 15 min at 800 × g and 4°C and the resulting supernatant was centrifuged for 15 min at 8,500 × g and 4 °C. The pellet was resuspended in 3 ml of STM buffer (250 mM sucrose, 20 mM Tris–HCl, 2 mM MgCl_2_, pH 8.0) with 40 U of DNase I (Thermo Scientific) and incubated for 30–60 min on ice. The DNase-treated lysate (5 ml) was loaded on top of a sucrose discontinuous density gradient in ultracentrifuge tubes (Cat. No. 344058, Beckman). The gradient was prepared by layering decreasingly dense sucrose solutions upon one another from 2.0, 1.75, 1.5, 1.25, 1, to 0.5 M, 5 ml each and was centrifuged in the SW-28 rotor at 87,041 × g (22,000 rpm) at 4°C for 4.5 h (L8-M Ultracentrifuge, Beckman). Ultracentrifugation separated the plastids from mitochondria and peroxisomes (Davis and Merrett 1973). The plastid fraction was situated at the interface of the 1.5 and 1.25 M sucrose and was collected by a syringe. Plastids were washed twice in SHE buffer to remove excessive sucrose. The final pellet was stored at −80°C for subsequent procedures. Figures documenting the resulting gradient and the assessment of the fractions quality by SDS-PAGE and immunoblot are included in the supplementary data (Fig. S1).

### Mass spectrometry-based protein identification and quantification

Samples were sonicated in NuPAGE LDS sample buffer (Thermo Scientific) containing 2 mM dithiothreitol and separated on a NuPAGE Bis-Tris 4–12% gradient polyacrylamide gel (Thermo Scientific) under reducing conditions. The sample lane was divided into eight slices that were excised from the Coomassie-stained gel, destained, and then subjected to tryptic digest and reductive alkylation. The treated fractions were subjected to liquid chromatography tandem mass spectrometry (LC-MS/MS) on an UltiMate 3000 RSLCnano System (Thermo Scientific) coupled to a Q exactive mass spectrometer (Thermo Scientific) performed by the Proteomic Facility at the University of Dundee. Mass spectra were analysed using MaxQuant (version 1.5, Cox & Mann, 2008), using the predicted translated transcriptome reported elsewhere (GEFR00000000.1; Ebenezer et al. 2019) as search database. Minimum peptide length was set to six amino acids, isoleucine and leucine were considered indistinguishable and false discovery rates (FDR) of 0.01 were calculated at the levels of peptides, proteins, and modification sites based on the number of hits against the reversed sequence database. Ratios were calculated from label-free quantification (LFQ) intensities using only peptides that could be uniquely mapped to a given protein. If the identified peptide sequence set of one protein contained the peptide set of another protein, these two proteins were assigned to the same protein group. P values were calculated applying t-test based statistics using Perseus (Cox and Mann 2008; Tyanova et al. 2016).

We identified 3,736 distinct protein groups using the MaxQuant analysis and for further analyses, data were reduced to 2,544 protein groups by rejecting those groups not identified at the peptide level in at least two of the three replicates for one organellar fraction. This filtered set includes a cohort of 774 protein groups that were observed in only one organellar fraction (168 and 606 in the mitochondrial and plastid fraction, respectively) and in order to form ratios, a constant small value (0.01) was added to the average LFQ intensities of each fraction (to avoid division by zero). The resulting CP/MT ratio reflects the enrichment of protein groups in the plastid fraction compared to the mitochondrial fraction and is the main indicator of the confidence in plastid localization of a given protein in this study. For clarity, CP/MT ratios were log_10_ transformed and values over 3 (CP/MT ratio > 1000) are indicated as “3+” in all tables and figures, indicating extremely high or infinite enrichment.

As proteins encoded in the *E. gracilis* plastid genome (Hallick et al., 1993) were missing from the translated transcriptome, the MaxQuant LFQ analysis was repeated using both search databases. The resulting quantifications of additional 32 plastid encoded proteins encoded were included in the plastid candidate dataset (supplementary-dataset-1.xlsx; seqid prefix “NP”). Additionally, we searched the translated transcriptome published by Yoshida et al. (2016; GDJR00000000.1). The obtained non-redundant identifications are provided in a separate sheet in supplementary-dataset-2.xlsx and were not included in the final plastid candidate dataset, but we did consider some of them in the reconstructions presented further below.

The purity of the plastid and mitochondrial fractions was assessed based on distribution and relative abundance of marker proteins (Figs. S1 and S2). LFQ analysis of the two organellar fractions and whole cell lysate, detected 8216 protein groups and revealed moderate contamination of both mitochondrial and plastid fractions by other cell compartments (Fig. S2.2 and Table S2.1). Comparison of the plastid and mitochondrial fractions suggests a modest degree of cross-contamination between the two organelles (Fig. S2.3).

### Proteome parsing, annotation, and sorting

Proteins enriched in the plastid fraction (log_10_ CP/MT ratio above zero) were considered plastid candidates. These were sorted into four categories based on log_10_ CP/MT ratio (0-1, 1-2, 2-3, and 3+) reflecting the reliability of their plastid localizations and were examined further. First, the set of candidates was annotated automatically using BLAST (Altschul et al. 1997) against the NCBI non-redundant protein database (https://www.ncbi.nlm.nih.gov/protein, August 2016 version), and assigned a KO number by KAAS http://www.genome.jp/tools/kaas/; Moriya et al., 2007). These annotations were checked manually consulting UniProt database (http://www.uniprot.org/) and OrthoFinder results and corrected as necessary. Proteins with no, very few, or very low-scoring homologs identifiable by BLAST were additionally searched by HHpred against PDB, COG, ECOD, and Pfam databases (https://toolkit.tuebingen.mpg.de/#/tools/hhpred; Soding et al., 2005) and, if possible, assigned at least vague annotation. Proteins with KO and/or EC numbers assigned, proteins of definite functions in certain metabolic pathways or molecular complexes, as well as proteins with more vaguely defined yet relatively clear functions were sorted manually into 18 custom-defined categories roughly based on the KEGG pathway classification (these were: "core metabolic pathways", "oxidative phosphorylation and electron transport", "photosynthesis", "carbohydrate metabolism", "lipid metabolism", "amino acid metabolism", "metabolism of cofactors and vitamins", "metabolism of terpenoids and polyketides", "DNA replication, recombination and repair", "transcription and transcription regulation", "RNA processing and degradation", "ribosome, aminoacyl-tRNA biosynthesis and translation", "protein transport, folding, processing, and degradation", "Fe-S cluster assembly and sulfur metabolism", "metabolite and ion transport", "regulation and signal transduction", "reaction to oxidative and toxic stress", and "other"; see Table S4 for more detailed description of each category). After the functional annotation, 32 proteins were discarded from the preliminary dataset of 1,377 candidates based on their clearly non-plastidial (i.e. mitochondrial, nuclear, or vacuolar) function combined with low enrichment values (log_10_ CP/MT ratio in the 0-1 range; supplementary-dataset-1.xlsx). The resulting plastid proteome consists of 1,345 proteins (for the full list with annotations and other details, see supplementary-dataset-1.xlsx).

Bipartite plastid targeting sequences were predicted using a combination of three available programs: SignalP (version 4.1, http://www.cbs.dtu.dk/services/SignalP/; Petersen et al., 2011) and PrediSI (http://www.predisi.de/; Hiller et al., 2004) for prediction of signal peptides, and ChloroP (http://www.cbs.dtu.dk/services/ChloroP/; Emanuelsson et al., 1999) for prediction of chloroplast transit peptides after *in silico* removal of the positively predicted signal peptides at their putative cleavage sites. The proteins were then annotated as having full bipartite signal, signal peptide only, transit peptide only, or no signal.

Major metabolic pathways were reconstructed from the proteome using the KEGG Mapper web tool (https://www.genome.jp/kegg/tool/map_pathway.html) combined with manual curation. Minor pathways with only a small number of members present and/or most of the members showing only weak evidence for plastid localization as judged by the enrichment (log_10_ CP/MT ratio only slightly above zero), as well as pathways of clearly non-plastidial localization were omitted.

### Determination of the evolutionary origin of plastid proteome

Each putative plastid protein was used as a query for a BLAST search against a custom database composed of 208 transcriptome and genome assemblies from eukaryotes, eubacteria, and archaebacteria, including 55 photosynthetic eukaryotes (dinoflagellates were omitted from the set due to complex plastid histories), 12 cyanobacteria, and 13 excavates (11 discobids and 2 non-discobids). For each of the *E. gracilis* sequences, a set of homologs with the e-value lower than 10^−3^ was retrieved with BLASTp, aligned with MAFFT (version 7.221; Katoh et al., 2002; Katoh and Standley, 2013), and trimmed automatically with TrimAl (version 1.2; Capella-Gutierrez et al., 2009). Maximum likelihood (ML) phylogenetic trees were constructed with RAxML (version 7.0.3; Stamatakis, 2006) from trimmed alignments longer than 74 sites and containing more than three unique sequences, and BS were calculated from 100 iterations. Tree topologies were analysed automatically and sorted based on the composition of the set of taxa clustered with the *E. gracilis* sequence in a bipartition supported by BS ≥ 75%. Only bipartitions formed by cutting internal (not terminal) branches were considered. The *E. gracilis* protein was assigned to one of the ten categories (Viridiplantae, Rhodophyta, Ochrophyta, Glaucophyta, Cryptophyta, Haptophyta, Chlorarachniophyta, Discoba, other Eukaryota or Prokaryota), if the bipartition comprised of *E. gracilis* and members of the concerned group only. If the bipartition comprised members of more than one concerned taxon of algae, the protein was assigned to one of the following assorted categories: primary algae (Viridiplantae, Rhodophyta, or Glaucophyta), primary or secondary green algae (Viridiplantae or Chlorarachniophyta), primary or secondary red algae (Rhodophyta, Ochrophyta, Haptophyta, or Cryptophyta), secondary red algae (Ochrophyta, Haptophyta, or Cryptophyta) or miscellaneous algae (any other combination of algal taxa). All trees were subsequently manually checked and are available at https://drive.google.com/drive/folders/13zdxl2CjdXhzB-OgrXqIJdq7YaQ1V9h2. The database and the detailed description of the bioinformatic pipeline will be published as a part of a related project and are currently available upon request.

### N-terminal domain analysis

The hydrophobicity and amino acid composition of the N-termini of plastid-targeted proteins were investigated. The analysis was restricted to proteins that were confidently non-truncated at the N-terminus, translated in cytoplasm, and imported into the plastid, i.e. that fulfilled the following criteria: a) highly credible plastid localization, i.e. log_10_ CP/MT ratio higher than one, or a clear photosynthetic function; and b) encoded by complete transcripts containing the spliced leader (ATTTTTTTTCG; Tessier et al., 1991). This cohort consisted of 375 sequences in total. Another set of the same size and characteristics, yet disregarding the criterion a), was selected randomly from the complete transcriptome of *E. gracilis* and used as a negative control.

The sequences were trimmed to the first 160 amino acids based on the expected length of the N-terminal signals (Sulli et al. 1999; Inagaki et al. 2000; Sláviková et al. 2005; Durnford and Gray 2006) and each position was assigned a hydrophobicity score using the Kyte-Doolittle scoring method with a window size of 13 residues (Kyte and Doolittle 1982); overall hydrophobicity and its distribution in different sequence positions was calculated. Using a custom script and parameters determined based on the hydrophobicity profile, the proteins from both the sample and the control set were sorted into preprotein classes: class I with two hydrophobic domains, i.e. the signal peptide (SP) and the stop-transfer signal (STS), the latter believed to serve as a transmembrane anchor in the transport vesicle (Sulli et al. 1999); class II with the SP only; and “unclassified” without any hydrophobic motif. The putative transit peptide (TP) region was approximated and extracted based on the positions of SP and STS (for class I), SP only (for class II), or, in the case of proteins without these hydrophobic regions, from its typical range as estimated from class I and II proteins. Next, the amino acid composition of the TP-like (TPL) region was investigated: the frequency of each amino acid in the TPL region was compared to the frequency of the same amino acid in the “mature” polypeptide (starting either at the position 140 or immediately after the STS region, if present, and ending at the C-terminus) of the same protein with the χ^2^ test performed using R (version 2.4.4.; R-Core-Team, 2013). Thus, a series of pairwise comparisons between the “TPL” and “mature” regions of the same sequence were done instead of a global screen across the whole set to avoid bias stemming from possible deviations from the average amino acid composition, as occurs in some specialized proteins (i.e. ion-binding proteins or proteins with multiple transmembrane domains). The test was performed both separately for each plastid protein class (I, II, and “unclassified”) as predicted by the custom script, and for the whole datasets of plastid proteins and the control sets of random proteins.

## Conclusions

The proteome of the *E. gracilis* plastid provides a wider outlook on the function, upkeep and origin of this organelle acquired by secondary endosymbiosis. Although majority of proteins with well-resolved phylogenies are affiliated to green algae/plants (donors of the plastid), non-negligible number of genes shows affiliation to chlorarachniophytes, ochrophytes, haptophytes and prokaryotes pointing to other potential players in the process of the plastid establishment. The detailed reconstruction of biochemical pathways elucidates number of remarkable metabolic and molecular quirks of this organism. *E. gracilis* possesses both mevalonate and non-mevalonate pathways for GGPP synthesis and the pathways for biosynthesis of carotenoids and tocopherols draw this precursor from distinct pools, however, a direct link between the tocopherol synthesis and non-plastidal mevalonate pathway is yet to be identified. Additionally, chlorophyll recycling appears to be an important source of phytol for the synthesis of tocopherols and other compounds. Our data also confirms plastid localization of the C5 pathway for aminolevulinate synthesis, while the Shemin pathway, also present in this organism, is localized elsewhere. The plastid has a surprisingly minor role in the *E. gracilis* amino acid metabolism, similar to its secondarily non-phototrophic cousin *E. longa* and in stark contrast to most plastid-bearing organisms. Interestingly, two sets of the SUF system are present in the plastid, the additional one of chlamydial evolutionary origin. We suspect the additional SUF might occupy its own plastid sub-compartment, providing FeS clusters to co-localized proteins, such as the plastidial sulfite reductase.

We find protein evidence for a single Tic protein in multiple isoforms, further suggesting that the TIC translocase “complex” might in fact consist of a single channel-forming subunit in *Euglena*. We confirm plastid localization of several Golgi- and ER- related proteins that are apparent paralog duplications, the most notable being GOSR1 and Rab5 which we propose serve as a part of a protein-transporting vesicle-docking system on the outermost plastid membrane. Two proteins distantly related to derlins are present in the plastid proteome and might represent the channel-forming core of a system analogous to SELMA of “chromalveolate” plastids and mediate protein transport across the middle plastid membrane, replacing the apparently absent TOC.

Additionally, a model set of nucleus-encoded high-confidence plastid proteins was used for the re-assessment of the plastid-targeting signals. This analysis was able to spot several novel features of these N-terminal peptides, which were previously overlooked as a result of the methodological constraints stemming from *in silico* localization predictions. One such feature is a significant proline enrichment of the TPL region which might impact its structure. A non-negotiable number of plastid-targeted proteins lack the typical N-terminal signal domain and are likely imported via an alternative mechanism and/or into a different plastid sub-compartment.

## Supporting information

derlins.fasta

## Acknowledgements

Authors would like to thank Anna Nenarokova (Biology Centre, České Budějovice) for suggestions and consultations regarding the data analysis and the Proteomics Facility at the University of Dundee for excellent technical service. This work was supported by Czech Grant Agency (15-21974S to V.H., 17-21409S to M.E.); ERC CZ (award LL1601 to J.L.); ERD Funds (project OPVVV 16_019/0000759 to J.L., M.C.F. and V.H.); Yousef Jameel Academic Program (through the Yousef Jameel PhD Scholarship); the Cambridge Commonwealth; European and International Trust; the Cambridge University Student Registry; the Cambridge Philosophical Society (all to T.E.E.) and the Medical Research Council (Grant #: P009018/1 to M.C.F.). Computational resources were supplied by the Ministry of Education, Youth and Sports of the Czech Republic under the Projects CESNET (Project No. LM2015042) and CERIT-Scientific Cloud (Project No. LM2015085) provided within the program Projects of Large Research, Development and Innovations Infrastructures.

## References

Altschul S, Madden TL, Schäffer AA, Zhang J, Zhang Z, Miller W, Lipman DJ. 1997. Gapped BLAST and PSI-BLAST: a new generation of protein database search programs. Nucleic Acids Res. 25(17):3389–3402.

Archibald JM. 2015. Genomic perspectives on the birth and spread of plastids. Proc Natl Acad Sci U S A. 112(33):1421374112-.

Balsera M, Soll J, Buchanan BB. 2010. Redox extends its regulatory reach to chloroplast protein import. Trends Plant Sci. 15(9):515–21. doi:10.1016/j.tplants.2010.06.002.

Barsanti L, Vismara R, Passarelli V, Gualtieri P. 2001. Paramylon (β-1,3-glucan) content in wild type and WZSL mutant of Euglena gracilis. Effects of growth conditions. J Appl Phycol. 13(1):59–65.

Becker B, Hoef-Emden K, Melkonian M. 2008. Chlamydial genes shed light on the evolution of photoautotrophic eukaryotes. BMC Evol Biol. 8(1):203. doi:10.1186/1471-2148-8-203.

Blee E, Schantz R. 1978. Biosynthesis of galactolipids in Euglena gracilis: I, Incorporation of UDP galactose into galactosyldiglycerides. Plant Sci Lett. 13:247–255.

Bölter B, Soll J. 2016. Once upon a Time - Chloroplast Protein Import Research from Infancy to Future Challenges. Mol Plant. 9(6):798–812.

Bruce BD. 2000. Chloroplast transit peptides: structure, function and evolution. Trends Cell Biol. 10(10):440–447. doi:10.1016/S0962-8924(00)01833-X.

Burki F, Imanian B, Hehenberger E, Hirakawa Y, Maruyama S, Keeling PJ. 2014. Endosymbiotic gene transfer in tertiary plastid-containing dinoflagellates. Eukaryot Cell. 13(2):246–55.

Capella-Gutierrez S, Silla-Martinez JM, Gabaldon T. 2009. trimAl: a tool for automated alignment trimming in large-scale phylogenetic analyses. Bioinformatics. 25(15):1972–1973.

Carrie C, Small I. 2013. A reevaluation of dual-targeting of proteins to mitochondria and chloroplasts. Biochim Biophys Acta - Mol Cell Res. 1833(2):253–259. doi:10.1016/J.BBAMCR.2012.05.029.

Cenci U, Ducatez M, Kadouche D, Colleoni C, Ball SG. 2016. Was the Chlamydial Adaptative Strategy to Tryptophan Starvation an Early Determinant of Plastid Endosymbiosis? Front Cell Infect Microbiol. 6:67.

Cenci U, Qiu H, Pillonel T, Cardol P, Remacle C, Colleoni C, Kadouche D, Chabi M, Greub G, Bhattacharya D, et al. 2018. Host-pathogen biotic interactions shaped vitamin K metabolism in Archaeplastida. Sci Rep. 8(1):15243.

Chen Y-L, Chen L-J, Chu C-C, Huang P-K, Wen J-R, Li H. 2018. TIC236 links the outer and inner membrane translocons of the chloroplast. Nature. 564(7734):125–129.

Cox J, Mann M. 2008. MaxQuant enables high peptide identification rates, individualized p.p.b.-range mass accuracies and proteome-wide protein quantification. Nat Biotechnol. 26(12):1367–1372.

Davis B, Merrett MJ. 1973. Malate Dehydrogenase Isoenzymes in Division Synchronized Cultures of Euglena. Plant Physiol. 51(6).

Deschamps P. 2014. Primary endosymbiosis: have cyanobacteria and Chlamydiae ever been roommates? Acta Soc Bot Pol. 83(4):291–302.

Disch A, Schwender J, Müller C, Lichtenthaler HK, Rohmer M. 1998. Distribution of the mevalonate and glyceraldehyde phosphate/pyruvate pathways for isoprenoid biosynthesis in unicellular algae and the cyanobacterium Synechocystis PCC 6714. Biochem J. 333(Pt 2)(2):381–8.

Dobáková E, Flegontov P, Skalický T, Lukeš J. 2015. Unexpectedly Streamlined Mitochondrial Genome of the Euglenozoan Euglena gracilis. Genome Biol Evol. 7(12):3358–3367.

Doetsch NA, Favreau MR, Kuscuoglu N, Thompson MD, Hallick RB. 2001. Chloroplast transformation in Euglena gracilis: splicing of a group III twintron transcribed from a transgenic psbK operon. Curr Genet. 39(1):49–60.

Domman D, Horn M, Embley TM, Williams TA. 2015. Plastid establishment did not require a chlamydial partner. Nat Commun. 6(1):6421.

Doolittle WF. 1998. You are what you eat: a gene transfer ratchet could account for bacterial genes in eukaryotic nuclear genomes. Trends Genet. 14(8):307–11.

van Dooren GG, Striepen B. 2013. The Algal Past and Parasite Present of the Apicoplast. Annu Rev Microbiol. 67(1):271–289.

Vom Dorp K, Hölzl G, Plohmann C, Eisenhut M, Abraham M, Weber APM, Hanson AD, Dörmann P. 2015. Remobilization of Phytol from Chlorophyll Degradation Is Essential for Tocopherol Synthesis and Growth of Arabidopsis. Plant Cell. 27(10):2846–59.

Dorrell RG, Howe CJ. 2015. Integration of plastids with their hosts: Lessons learned from dinoflagellates. Proc Natl Acad Sci U S A. 112(33):10247–54.

Duchêne A-M, Giritch A, Hoffmann B, Cognat V, Lancelin D, Peeters NM, Zaepfel M, Maréchal-Drouard L, Small ID. 2005. Dual targeting is the rule for organellar aminoacyl-tRNA synthetases in Arabidopsis thaliana. Proc Natl Acad Sci U S A. 102(45):16484–9.

Durnford DG, Gray MW. 2006. Analysis of Euglena gracilis plastid-targeted proteins reveals different classes of transit sequences. Eukaryot Cell. 5(12):2079–91.

Ebenezer TE, Zoltner M, Burrel A, Nenarokova A, Novák Vanclová AMG, Prasad B, Soukal P, Santana-Molina C, O'Neill E, Nankissoor NN et al. 2019.Transcriptome, proteome and draft genome of Euglena gracilis. BMC Biology. 17(1):11.

Ehrenberg CG. 1830. Organisation, Systematik und geographisches Verhältniss der Infusionsthierchen. Berlin: Druckerei der Königlichen Akademie der Wissenschaften.

Emanuelsson O, Nielsen H, von Heijne G. 1999. ChloroP, a neural network-based method for predicting chloroplast transit peptides and their cleavage sites. Protein Sci. 8(5):978–984.

Emonds-Alt B, Coosemans N, Gerards T, Remacle C, Cardol P. 2017. Isolation and characterization of mutants corresponding to the MENA, MENB, MENC and MENE enzymatic steps of 5’-monohydroxyphylloquinone biosynthesis in Chlamydomonas reinhardtii. Plant J. 89(1):141–154.

Facchinelli F, Colleoni C, Ball SG, Weber APM. 2013. Chlamydia, cyanobiont, or host: who was on top in the ménage à trois? Trends Plant Sci. 18(12):673–9.

Felsner G, Sommer MS, Gruenheit N, Hempel F, Moog D, Zauner S, Martin W, Maier UG. 2011. ERAD components in organisms with complex red plastids suggest recruitment of a preexisting protein transport pathway for the periplastid membrane. Genome Biol Evol. 3(0):140–50.

Felsner G, Sommer MS, Maier UG. 2010. The physical and functional borders of transit peptide-like sequences in secondary endosymbionts. BMC Plant Biol. 10(1):223.

Geimer S, Belicová A, Legen J, Sláviková S, Herrmann RG, Krajcovic J. 2009. Transcriptome analysis of the Euglena gracilis plastid chromosome. Curr Genet. 55(4):425–38.

Gile GH, Moog D, Slamovits CH, Maier U-G, Archibald JM. 2015. Dual Organellar Targeting of Aminoacyl-tRNA Synthetases in Diatoms and Cryptophytes. Genome Biol Evol. 7(6):1728–1742.

Gould SB, Maier U-G, Martin WF. 2015. Protein Import and the Origin of Red Complex Plastids. Curr Biol. 25(12):R515–R521.

Gumińska N, Płecha M, Zakryś B, Milanowski R. 2018. Order of removal of conventional and nonconventional introns from nuclear transcripts of Euglena gracilis. Field MC, editor. PLOS Genet. 14(10):e1007761.

Hallick RB, Hong L, Drager RG, Favreau MR, Monfort A, Orsat B, Spielmann A, Stutz E. 1993. Complete sequence of Euglena gracilis chloroplast DNA. Nucleic Acids Res. 21(15):3537–3544.

Hannaert V, Brinkmann H, Nowitzki U, Lee JA, Albert M-A, Sensen CW, Gaasterland T. M M, Michels P, Martin W. 2000. Enolase from Trypanosoma brucei, from the Amitochondriate Protist Mastigamoeba balamuthi, and from the Chloroplast and Cytosol of Euglena gracilis: Pieces in the Evolutionary Puzzle of the Eukaryotic Glycolytic Pathway. Mol Biol Evol. 17(7):989–1000.

Harris J. 1695. Some Microscopical Observations of Vast Numbers of Animalcula Seen in Water by John Harris, M. A. Rector of Winchelsea in Sussex, and F. R. S. Philos Trans R Soc London. 19(215-235):254–259.

Hempel F, Bullmann L, Lau J, Zauner S, Maier UG. 2009. ERAD-derived preprotein transport across the second outermost plastid membrane of diatoms. Mol Biol Evol. 26(8):1781–90.

Heyes DJ, Neil Hunter C. 2009. Biosynthesis of Chlorophyll and Bacteriochlorophyll. In: Tetrapyrroles. New York, NY: Springer New York. p. 235–249.

Hiller K, Grote A, Scheer M, Münch R, Jahn D. 2004. PrediSi: Prediction of signal peptides and their cleavage positions. Nucleic Acids Res. 32(WEB SERVER ISS.):375–379.

Hopkins JF, Spencer DF, Laboissiere S, Neilson JAD, Eveleigh RJM, Durnford DG, Gray MW, Archibald JM. 2012. Proteomics Reveals Plastid- and Periplastid-Targeted Proteins in the Chlorarachniophyte Alga Bigelowiella natans. Genome Biol Evol. 4(12):1391–1406.

Howe CJ, Schlarb-Ridley BG, Wastl J, Purton S, Bendall DS. 2006. The novel cytochrome c6 of chloroplasts: a case of evolutionary bricolage? J Exp Bot. 57(1):13–22.

Huang M, Friso G, Nishimura K, Qu X, Olinares PDB, Majeran W, Sun Q, van Wijk KJ. 2013. Construction of Plastid Reference Proteomes for Maize and *Arabidopsis* and Evaluation of Their Orthologous Relationships; The Concept of Orthoproteomics. J Proteome Res. 12(1):491–504.

Inagaki J, Fujita Y, Hase T, Yamamoto Y. 2000. Protein translocation within chloroplast is similar in Euglena and higher plants. Biochem Biophys Res Commun. 277(2):436–442.

Jackson C, Knoll AH, Chan CX, Verbruggen H. 2018. Plastid phylogenomics with broad taxon sampling further elucidates the distinct evolutionary origins and timing of secondary green plastids. Sci Rep. 8(1):1523.

Jenkins KP, Hong L, Hallick RB. 1995. Alternative splicing of the Euglena gracilis chloroplast roaA transcript. RNA. 1(6):624–633.

Karnkowska A, Vacek V, Zubáčová Z, Treitli SC, Petrželková R, Eme L, Novák L, Žárský V, Barlow LD, Herman EK, et al. 2016. A Eukaryote without a Mitochondrial Organelle. Curr Biol. 26(10):1274–84.

Katoh K, Misawa K, Kuma K, Miyata T. 2002. MAFFT: a novel method for rapid multiple sequence alignment based on fast Fourier transform. Nucleic Acids Res. 30(14):3059–3066.

Katoh K, Standley DM. 2013. MAFFT Multiple Sequence Alignment Software Version 7: Improvements in Performance and Usability. Mol Biol Evol. 30(4):772–780. doi:10.1093/molbev/mst010.

Keeling PJ, Palmer JD. 2008. Horizontal gene transfer in eukaryotic evolution. Nat Rev Genet. 9(8):605–618.

Kikuchi S, Asakura Y, Imai M, Nakahira Y, Kotani Y, Hashiguchi Y, Nakai Y, Takafuji K, Bédard J, Hirabayashi-Ishioka Y, et al. 2018 Oct 11. A Ycf2-FtsHi heteromeric AAA-ATPase complex is required for chloroplast protein import. Plant Cell.:tpc.00357.2018.

Kikuchi S, Oishi M, Hirabayashi Y, Lee DW, Hwang I, Nakai M. 2009. A 1-megadalton translocation complex containing Tic20 and Tic21 mediates chloroplast protein import at the inner envelope membrane. Plant Cell. 21(6):1781–97.

Kim D, Filtz MR, Proteau PJ. 2004. The methylerythritol phosphate pathway contributes to carotenoid but not phytol biosynthesis in Euglena gracilis. J Nat Prod. 67(6):1067–1069.

Kořený L, Oborník M. 2011. Sequence evidence for the presence of two tetrapyrrole pathways in Euglena gracilis. Genome Biol Evol. 3(1):359–364.

Krinsky NI, Goldsmith TH. 1960. The carotenoids of the flagellated alga, Euglena gracilis. Arch Biochem Biophys. 91(12):271–279.

Kuo RC, Zhang H, Zhuang Y, Hannick L, Lin S. 2013. Transcriptomic Study Reveals Widespread Spliced Leader Trans-Splicing, Short 5’-UTRs and Potential Complex Carbon Fixation Mechanisms in the Euglenoid Alga Eutreptiella sp. PLoS One. 8(4):e60826.

Kyte J, Doolittle RF. 1982. A simple method for displaying the hydropathic character of a protein. J Mol Biol. 157(1):105–132.

Lakey B, Triemer R. 2017. The tetrapyrrole synthesis pathway as a model of horizontal gene transfer in euglenoids. Müller K, editor. J Phycol. 53(1):198–217.

Lange BM, Rujan T, Martin W, Croteau R. 2000. Isoprenoid biosynthesis: the evolution of two ancient and distinct pathways across genomes. Proc Natl Acad Sci U S A. 97(24):13172–7.

Langemeyer L, Fröhlich F, Ungermann C. 2018. Rab GTPase Function in Endosome and Lysosome Biogenesis. Trends Cell Biol. 28(11):957–970.

Larkum AWD, Lockhart PJ, Howe CJ. 2007. Shopping for plastids. Trends Plant Sci. 12(5):189–195.

Lau JB, Stork S, Moog D, Schulz J, Maier UG. 2016. Protein-protein interactions indicate composition of a 480 kDa SELMA complex in the second outermost membrane of diatom complex plastids. Mol Microbiol. 100(1):76–89.

Leander BS. 2004. Did trypanosomatid parasites have photosynthetic ancestors? Trends Microbiol. 12(6):251–258.

Leander BS, Lax G, Karnkowska A, Simpson AGB. 2017. Euglenida. In: Handbook of the Protists. Cham: Springer International Publishing. p. 1–42.

Leander BS, Triemer RE, Farmer M a. 2001. Character evolution in heterotrophic euglenids. Eur J Protistol. 37:337–356.

Leger MM, Eme L, Hug LA, Roger AJ. 2016. Novel Hydrogenosomes in the Microaerophilic Jakobid *Stygiella incarcerata*. Mol Biol Evol. 33(9):2318–2336.

Li H min, Teng YS. 2013. Transit peptide design and plastid import regulation. Trends Plant Sci. 18(7):360–366.

Lin Y-P, Wu M-C, Charng Y-Y. 2016. Identification of a Chlorophyll Dephytylase Involved in Chlorophyll Turnover in Arabidopsis. Plant Cell. 28(12):2974–2990.

Lohr M, Schwender J, Polle JEW. 2012. Isoprenoid biosynthesis in eukaryotic phototrophs: A spotlight on algae. Plant Sci. 185-186:9–22.

Lu Y. 2018. Assembly and Transfer of Iron-Sulfur Clusters in the Plastid. Front Plant Sci. 9:336.

MacArthur MW, Thornton JM. 1991. Influence of proline residues on protein conformation. J Mol Biol. 218(2):397–412.

Maeda H, Dudareva N. 2012. The Shikimate Pathway and Aromatic Amino Acid Biosynthesis in Plants. Annu Rev Plant Biol. 63(1):73–105.

Maier UG, Zauner S, Hempel F. 2015 Jun 1. Protein import into complex plastids: Cellular organization of higher complexity. Eur J Cell Biol.

Marin B, Nowack ECM, Melkonian M. 2005. A plastid in the making: evidence for a second primary endosymbiosis. Protist. 156(4):425–32.

Markunas CM, Triemer RE. 2016. Evolutionary History of the Enzymes Involved in the Calvin-Benson Cycle in Euglenids. J Eukaryot Microbiol. 63(3):326–339.

Maruyama S, Suzaki T, Weber APM, Archibald JM, Nozaki H. 2011. Eukaryote-to-eukaryote gene transfer gives rise to genome mosaicism in euglenids. BMC Evol Biol. 11(1):105.

Mateášiková-Kováčová B, Vesteg M, Drahovská H, Záhonová K, Vacula R, Krajčovič J. 2012. Nucleus-encoded mRNAs for chloroplast proteins GapA, PetA, and PsbO are trans-spliced in the flagellate Euglena gracilis irrespective of light and plastid function. J Eukaryot Microbiol. 59(6):651–653.

Matson RS, Meifei, Chang SB. 1970. Comparative studies of biosynthesis of galactolipids in Euglena-gracilis strain-Z. Plant Physiol. 45(4):531-.

Matsumoto T, Shinozaki F, Chikuni T, Yabuki A, Takishita K, Kawachi M, Nakayama T, Inouye I, Hashimoto T, Inagaki Y. 2011. Green-colored Plastids in the Dinoflagellate Genus Lepidodinium are of Core Chlorophyte Origin. Protist. 162(2):268–276.

Minge M a, Shalchian-Tabrizi K, Tørresen OK, Takishita K, Probert I, Inagaki Y, Klaveness D, Jakobsen KS. 2010. A phylogenetic mosaic plastid proteome and unusual plastid-targeting signals in the green-colored dinoflagellate Lepidodinium chlorophorum. BMC Evol Biol. 10:191.

Moreno-Sánchez R, Covián R, Jasso-Chávez R, Rodríguez-Enríquez S, Pacheco-Moisés F, Torres-Márquez ME. 2000. Oxidative phosphorylation supported by an alternative respiratory pathway in mitochondria from Euglena. Biochim Biophys Acta. 1457(3):200–10.

Moriya Y, Itoh M, Okuda S, Yoshizawa AC, Kanehisa M. 2007. KAAS: an automatic genome annotation and pathway reconstruction server. Nucleic Acids Res. 35(Web Server issue):W182–5.

Moustafa A, Reyes-Prieto A, Bhattacharya D. 2008. Chlamydiae Has Contributed at Least 55 Genes to Plantae with Predominantly Plastid Functions. DeSalle R, editor. PLoS One. 3(5):e2205.

Muchhal US, Schwartzbach SD. 1994. Characterization of the unique intron-exon junctions of Euglena gene(s) encoding the polyprotein precursor to the light-harvesting chlorophyll a/b binding protein of photosystem II. Nucleic Acids Res. 22(25):5737–44.

Nawrocki WJ, Tourasse NJ, Taly A, Rappaport F, Wollman F-A. 2015. The Plastid Terminal Oxidase: Its Elusive Function Points to Multiple Contributions to Plastid Physiology. Annu Rev Plant Biol. 66(1):49–74.

Nowack ECM, Price DC, Bhattacharya D, Singer A, Melkonian M, Grossman AR. 2016. Gene transfers from diverse bacteria compensate for reductive genome evolution in the chromatophore of Paulinella chromatophora. Proc Natl Acad Sci U S A. 113(43):12214–12219.

O’Neill EC, Trick M, Hill L, Rejzek M, Dusi RG, Hamilton CJ, Zimba P V., Henrissat B, Field RA. 2015. The transcriptome of Euglena gracilis reveals unexpected metabolic capabilities for carbohydrate and natural product biochemistry. Mol Biosyst. 11(10):2808–2820.

Patron NJ, Waller RF. 2007. Transit peptide diversity and divergence: A global analysis of plastid targeting signals. Bioessays. 29(10):1048–58.

Petersen TN, Brunak S, von Heijne G, Nielsen H. 2011. SignalP 4.0: discriminating signal peptides from transmembrane regions. Nat Methods. 8(10):785–6.

Ponce-Toledo RI, Moreira D, López-García P, Deschamps P, Ruiz-Trillo I. 2018 Jun 19. Secondary Plastids of Euglenids and Chlorarachniophytes Function with a Mix of Genes of Red and Green Algal Ancestry. Ruiz-Trillo I, editor. Mol Biol Evol.

R-Core-Team. 2013. R: A language and environment for statistical computing.

Reyes-Prieto A, Moustafa A. 2012. Plastid-localized amino acid biosynthetic pathways of Plantae are predominantly composed of non-cyanobacterial enzymes. Sci Rep. 2(1):955.

Rippert P, Puyaubert J, Grisollet D, Derrier L, Matringe M. 2009. Tyrosine and Phenylalanine Are Synthesized within the Plastids in Arabidopsis. PLANT Physiol. 149(3):1251–1260.

Saidha T, Na SQ, Li JY, Schiff JA. 1988. A sulphate metabolizing centre in Euglena mitochondria. Biochem J. 253(2):533–9.

Saidha T, Schiff JA. 1989. The role of mitochondria in sulfolipid biosynthesis by Euglena chloroplasts. Biochim Biophys Acta - Lipids Lipid Metab. 1001(3):268–273.

Saldarriaga JF, Taylor FJR, Keeling PJ, Cavalier-Smith T. 2001. Dinoflagellate Nuclear SSU rRNA Phylogeny Suggests Multiple Plastid Losses and Replacements. J Mol Evol. 53(3):204–213.

Schwartzbach SD, Shigeoka S. 2017. Euglena : Biochemistry, Cell and Molecular Biology. Cham: Springer International Publishing.

Seeger JW, Bentley R. 1991. Phylloquinone (Vitamin K1) biosynthesis inEuglena gracilis strain Z. Phytochemistry. 30(11):3585–3589.

Sheiner L, Striepen B. 2013. Protein sorting in complex plastids. Biochim Biophys Acta-Mol Cell Res. 1833(2):352–359.

Shibata S, Arimura S, Ishikawa T, Awai K. 2018. Alterations of Membrane Lipid Content Correlated With Chloroplast and Mitochondria Development in Euglena gracilis. Front Plant Sci. 9:370.

Sláviková S, Vacula R, Fang Z, Ehara T, Osafune T, Schwartzbach SD. 2005. Homologous and heterologous reconstitution of Golgi to chloroplast transport and protein import into the complex chloroplasts of Euglena. J Cell Sci. 118(Pt 8):1651–1661.

Smith KW, Stroupe ME. 2012. Mutational Analysis of Sulfite Reductase Hemoprotein Reveals the Mechanism for Coordinated Electron and Proton Transfer. Biochemistry. 51(49):9857–9868.

Soding J, Biegert A, Lupas AN. 2005. The HHpred interactive server for protein homology detection and structure prediction. Nucleic Acids Res. 33(Web Server):W244–W248.

Sommer MS, Gould SB, Lehmann P, Gruber A, Przyborski JM, Maier U-G. 2007. Der1-mediated Preprotein Import into the Periplastid Compartment of Chromalveolates? Mol Biol Evol. 24(4):918–928.

Spork S, Hiss J a, Mandel K, Sommer M, Kooij TW a, Chu T, Schneider G, Maier UG, Przyborski JM. 2009. An unusual ERAD-like complex is targeted to the apicoplast of Plasmodium falciparum. Eukaryot Cell. 8(8):1134–45.

Stairs CW, Eme L, Brown MW, Mutsaers C, Susko E, Dellaire G, Soanes DM, van der Giezen M, Roger AJ. 2014. A SUF Fe-S Cluster Biogenesis System in the Mitochondrion-Related Organelles of the Anaerobic Protist Pygsuia. Curr Biol. 24(11):1176–1186.

Stamatakis A. 2006. RAxML-VI-HPC: maximum likelihood-based phylogenetic analyses with thousands of taxa and mixed models. Bioinformatics. 22(21):2688–2690.

Stiller JW, Schreiber J, Yue J, Guo H, Ding Q, Huang J. 2014. The evolution of photosynthesis in chromist algae through serial endosymbioses. Nat Commun. 5(1):5764.

Subtil A, Collingro A, Horn M. 2014. Tracing the primordial Chlamydiae: extinct parasites of plants? Trends Plant Sci. 19(1):36–43.

Sulli C, Fang ZW, Muchhal U, Schwartzbach SD. 1999. Topology of Euglena chloroplast protein precursors within endoplasmic reticulum to Golgi to chloroplast transport vesicles. J Biol Chem. 274(1):457–463.

Takano Y, Hansen G, Fujita D, Horiguchi T. 2008. Serial Replacement of Diatom Endosymbionts in Two Freshwater Dinoflagellates, Peridiniopsis spp. (Peridiniales, Dinophyceae). Phycologia. 47(1):41–53.

Tanaka Y, Ogawa T, Maruta T, Yoshida Y, Arakawa K, Ishikawa T. 2017. Glucan synthase-like 2 is indispensable for paramylon synthesis in Euglena gracilis. FEBS Lett. 591(10):1360–1370.

Teng Y-S, Su Y, Chen L-J, Lee YJ, Hwang I, Li H. 2006. Tic21 is an essential translocon component for protein translocation across the chloroplast inner envelope membrane. Plant Cell. 18(9):2247–57.

Terashima M, Specht M, Hippler M. 2011. The chloroplast proteome: a survey from the Chlamydomonas reinhardtii perspective with a focus on distinctive features. Curr Genet. 57(3):151–68.

Tessier LH, Keller M, Chan RL, Fournier R, Weil JH, Imbault P. 1991. Short leader sequences may be transferred from small RNAs to pre-mature mRNAs by trans-splicing in Euglena. EMBO J. 10(9):2621–5.

Tsaousis AD, Ollagnier de Choudens S, Gentekaki E, Long S, Gaston D, Stechmann A, Vinella D, Py B, Fontecave M, Barras F, et al. 2012. Evolution of Fe/S cluster biogenesis in the anaerobic parasite Blastocystis. Proc Natl Acad Sci. 109(26):10426–10431.

Turmel M, Gagnon M-C, O’Kelly CJ, Otis C, Lemieux C. 2009. The chloroplast genomes of the green algae Pyramimonas, Monomastix, and Pycnococcus shed new light on the evolutionary history of prasinophytes and the origin of the secondary chloroplasts of euglenids. Mol Biol Evol. 26(3):631–48.

Tyanova S, Temu T, Sinitcyn P, Carlson A, Hein MY, Geiger T, Mann M, Cox J. 2016. The Perseus computational platform for comprehensive analysis of (prote)omics data. Nat Methods. 13(9):731–740.

Watanabe F, Yoshimura K, Shigeoka S. 2017. Biochemistry and Physiology of Vitamins in Euglena. In: Schwartzbach SD, Shigeoka S, editors. Euglena: Biochemistry, Cell and Molecular Biology. Cham: Springer International Publishing. p. 65–90.

Wetherbee R, Jackson CJ, Repetti SI, Clementson LA, Costa JF, van de Meene A, Crawford S, Verbruggen H. 2018 Dec 9. The golden paradox -a new heterokont lineage with chloroplasts surrounded by two membranes. J Phycol.:jpy.12822.

Wildner GF, Hauska G. 1974. Localization of the reaction site of cytochrome 552 in chloroplasts from Euglena gracilis: Cytochrome content and photooxidation in different chloroplast preparations. Arch Biochem Biophys. 164(1):127–135.

Xia S, Zhang Q, Zhu H, Cheng Y, Liu G, Hu Z. 2013. Systematics of a Kleptoplastidal Dinoflagellate, Gymnodinium eucyaneum Hu (Dinophyceae), and Its Cryptomonad Endosymbiont. Söderhäll I, editor. PLoS One. 8(1):e53820.

Yamaguchi A, Yubuki N, Leander BS. 2012. Morphostasis in a novel eukaryote illuminates the evolutionary transition from phagotrophy to phototrophy: description of Rapaza viridis n. gen. et sp. (Euglenozoa, Euglenida). BMC Evol Biol. 12(1):29.

Yoon HS, Hackett JD, Van Dolah FM, Nosenko T, Lidie KL, Bhattacharya D. 2005. Tertiary endosymbiosis driven genome evolution in dinoflagellate algae. Mol Biol Evol. 22(5):1299–308.

Yoshida Y, Tomiyama T, Maruta T, Tomita M, Ishikawa T, Arakawa K. 2016. De novo assembly and comparative transcriptome analysis of Euglena gracilis in response to anaerobic conditions. BMC Genomics. 17(1):182.

Záhonová K, Füssy Z, Birčák E, Novák Vanclová AMG, Klimeš V, Vesteg M, Krajčovič J, Oborník M, Eliáš M. 2018. Peculiar features of the plastids of the colourless alga Euglena longa and photosynthetic euglenophytes unveiled by transcriptome analyses. Sci Rep. 8(1):17012.

Zhao L, Chang W, Xiao Y, Liu H, Liu P. 2013. Methylerythritol Phosphate Pathway of Isoprenoid Biosynthesis. Annu Rev Biochem 82(1):497–530.

Ziegler K, Maldener I, Lockau W. 1989. 5’-Monohydroxyphylloquinone as a Component of Photosystem I. Zeitschrift für Naturforsch C. 44(5–6):468–472.

