## supplementary data for "Proteome of the secondary plastid of *Euglena gracilis* reveals metabolic quirks and colourful history"

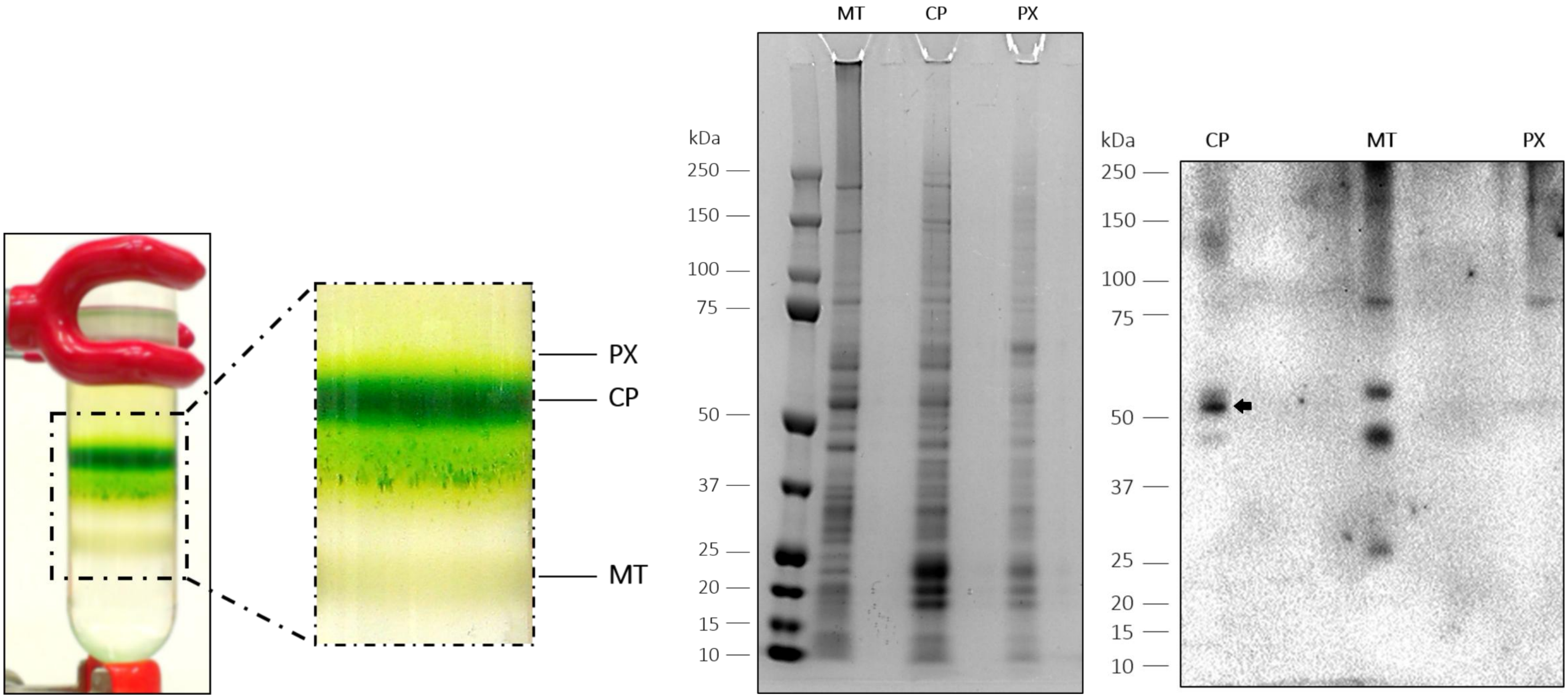


S1.1: The result of cell fractionation on a sucrose gradient: the plastidial fraction is marked as CP, while the mitochondrial and peroxisomal fractions are marked MT and PX, respectively.

S1.2: Coomasie-stained gel (Nupage Bis-Tris Mini Gels 4-12%, IM-8042, Life Technologies) of the cellular fractions with Precision Plus Protein™ Dual Color Standards (#1610374, Bio-Rad).

S1.3: Immunoblot of the cellular fractions with rabbit anti-RbcL (1:5000, AS03 037, Agrisera, protein size: 52 kDa) as primary antibody and goat anti-Rabbit (1:2000, A6154-5x, Sigma) as secondary antibody, and Protein Ladder (161-0374, Biorad), the positive band of expected size is marked by arrow.

| | Cytosol | Acidocalcisome | Endosome | Peroxisome | Lysosome | Golgi | Nucleus | Surface |
|---|---|---|---|---|---|---|---|---|
| Protein ID EG_transcript_ | **21524** | **2633** | **181** | **15991** | **10514** | **5712** | **53416** | **32527** |
| Annotation | Aldolase | VP1 Vacuolar proton translocating pyrophosphatase | CHC Clathrin heavy chain | PEX2 Peroxisomal biogenesis factor 2 | Lysosomal aspartic protease | Coatomer subunit γ2 | Histone H4 | Articulin 80 kDa |
| **Ratio CP/W** | **0.01** | **1.95** | **0.21** | **W only** | **3.30** | **0.01** | **0.04** | **0.01** |
| -log10p | 5.9 | 1.2 | 3.1 | 1.4 | 2.0 | 3.4 | 3.3 | 4.7 |
| Ratio Mt/W | 0.02 | 1.46 | 0.53 | W only | 7.4 | 0.01 | 0.02 | 0.04 |
| -log10p | 3.3 | 0.5 | 1.9 | 2.2 | 2.6 | 3.3 | 3.6 | 4.4 |
| Ratio CP/Mt | 0.26 | 1.33 | 0.40 | NaN | 0.44 | 0.74 | 2.0 | 0.15 |
| -log10p | 1.5 | 0.6 | 2.8 | NaN | 1.5 | 0.3 | 0.8 | 2.9 |
| Unique peptides | 19 | 31 | 90 | 1 | 7 | 40 | 20 | 13 |

| | Chloroplast | | | Mitochondrion | | |
|---|---|---|---|---|---|---|
| Protein ID EG_transcript_ | **40006** | **158** | **25897** | **2112** | **23844** | **8912** |
| Annotation | light-harvesting complex I protein precursor LhcB5 | Photosystem I P700 chlorophyll a apoprotein A1 | light-harvesting complex I protein precursor Lhca2 | Pyruvate dehydrogenase [NADP(+)], mitochondrial | ubiquinol-cytochrome c reductase iron-sulfur subunit | F-type H+-transporting ATPase subunit beta |
| **Ratio CP/W** | **4.5** | **1.95** | **CP only** | **1.31** | **3.10** | **4.6** |
| -log10p | 3.2 | 1.7 | 1.9 | 0.6 | 4.4 | 3.8 |
| Ratio Mt/W | 0.2 | 0.10 | NaN | 4.7 | 4.27 | 4.8 |
| -log10p | 1.8 | 2.3 | NaN | 2.6 | 3.5 | 3.4 |
| Ratio CP/Mt | 24.2 | 20.5 | CP only | 0.28 | 0.73 | 0.95 |
| -log10p | 2.7 | 2.8 | 2.2 | 2.1 | 1.3 | 0.1 |
| Unique peptides | 4 | 30 | 2 | 76 | 35 | 32 |

S2.1: Assessment of the purity of isolated fractions using selected marker proteins and their relative abundance compared against the whole cell lysate and between the organellar fractions.

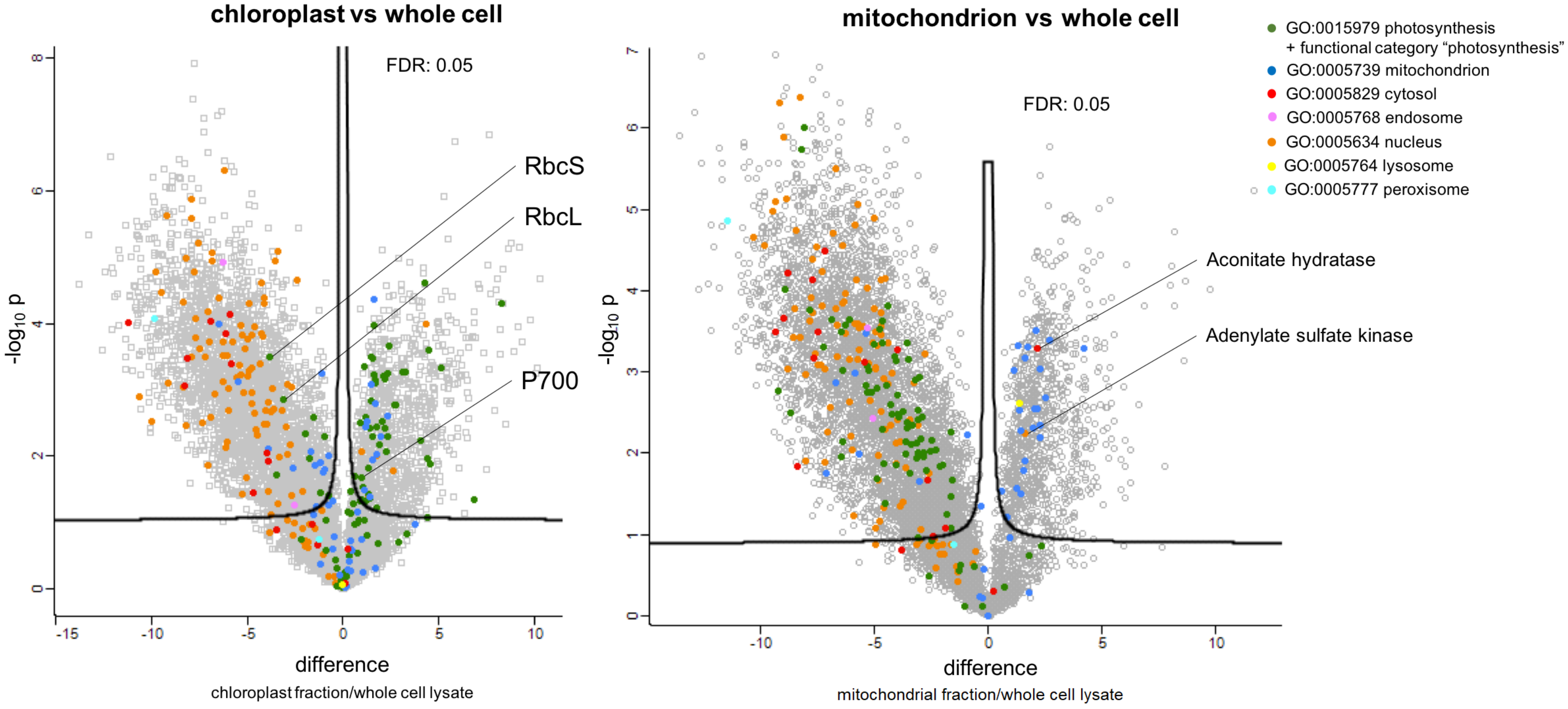


S2.2: Volcano plots from p-values versus the corresponding t-test difference of 8216 protein groups quantified in the two organellar fractions and whole cell lysate. Green and blue dots represent proteins assigned to “photosynthetic” and “mitochondrial” GO categories, respectively. The remaining colours represent other selected GO categories (indicated at the top right) associated with other cellular compartments. Stringent cutoff curves for statistically significant enrichment (black curves) were calculated from the estimated false discovery rate (FDR).

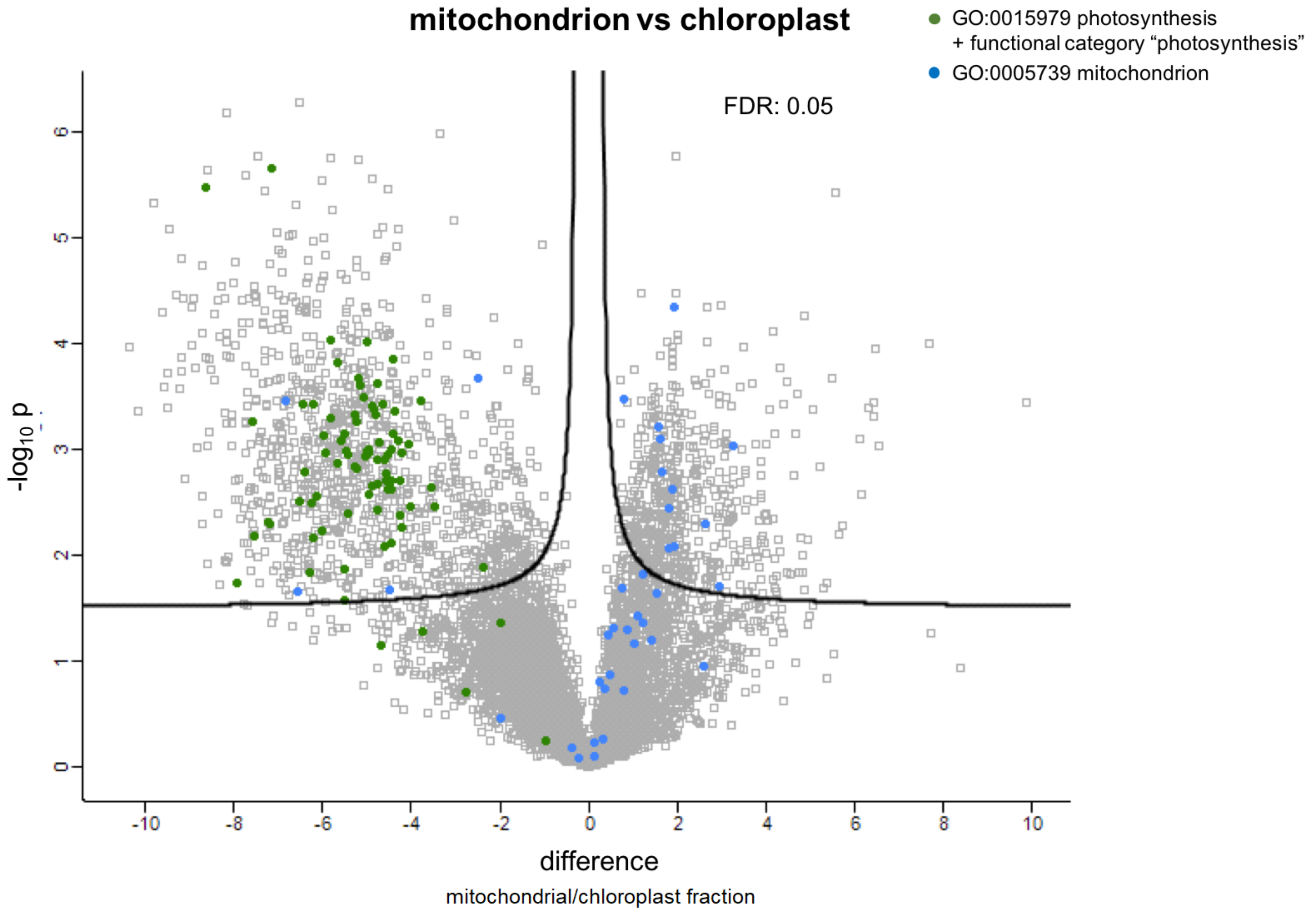


S2.3: Volcano plot from p-values versus the corresponding t-test difference of 3,736 protein groups quantified in the mitochondrial and chloroplast fraction. Green and blue dots represent proteins assigned to "photosynthetic" and "mitochondrial" GO categories, respectively. Stringent cutoff curves for statistically significant enrichment (black curves) were calculated from the estimated false discovery rate (FDR).

| custom category name | description |
|---|---|
| **protein transport, folding, processing, and degradation** | protein translocases of plastid envelope and thylakoid membranes, signal peptidases, heat shock proteins, enzymes of post-translational protein modifications such as methylation, acetylation, glycosylation, proline cis/trans isomerisation, disulfide bond formation and breakage, and components of protein degradation systems |
| **metabolite and ion transport** | proteins involved in transport of all non-protein compounds, including predicted membrane transporters of undetermined substrates |
| **photosynthesis** | components of photosystems, light-harvesting antennae, cytochrome b6/f complex and proteins involved in their biogenesis |
| **ribosome, aminoacyl-tRNA biosynthesis and translation** | ribosomal proteins and proteins involved in ribosome biogenesis, aminoacyl-tRNA synthetases, translation regulators |
| **regulation and signal transduction** | proteins involved in other than transcriptional and translational regulation, signaling molecules and their receptors, not-further-specified protein kinases, phosphatases, adenylate cyclases and similar enzymes typically involved in signal transduction |
| **metabolism of cofactors and vitamins** | mostly proteins involved in chlorophyll biosynthesis, several enzymes of metabolism of ubiquinone and retinol |
| **lipid metabolism** | enzymes of fatty acid biosynthesis, elongation, modification and degradation, synthesis of glycerolipids and glycerophospholipids |
| **core metabolic pathways** | proteins of glycolysis, pentose phosphate pathway, pyruvate metabolism, carbon fixation, one-carbon and acetyl-CoA metabolism |
| **oxidative phosphorylation and electron transport** | components of ATP synthase and electron transport chain |
| **transcription and transcription regulation** | transcription and translation factors and other proteins involved in gene expression regulation |
| **RNA processing and degradation** | RNAses and other enzymes responsible for RNA splicing, maturation and degradation |
| **metabolism of terpenoids and polyketides** | proteins involved in biosynthesis of terpenoids and carotenoids |
| **DNA replication, recombination and repair** | DNA polymerases, ligases, helicases, proteins involved in DNA maintenance and repair |
| **reaction to oxidative and toxic stress** | enzymes involved in detoxification of xenobiotics and protection from reactive oxygen species and photo oxidative damage, proteins responsible for redox balance |
| **amino acid metabolism** | proteins involved in amino acid synthesis and interconversions, enzymes of shikimate pathway |
| **FeS cluster assembly and sulfur metabolism** | components of SUF pathway, proteins involved in metabolism of sulfur compounds |
| **carbohydrate metabolism** | enzymes of starch and other saccharides metabolism |
| **other** | members of other pathways which were minor in comparison to other functions |

S3: List of the custom protein categories with their descriptions and examples.

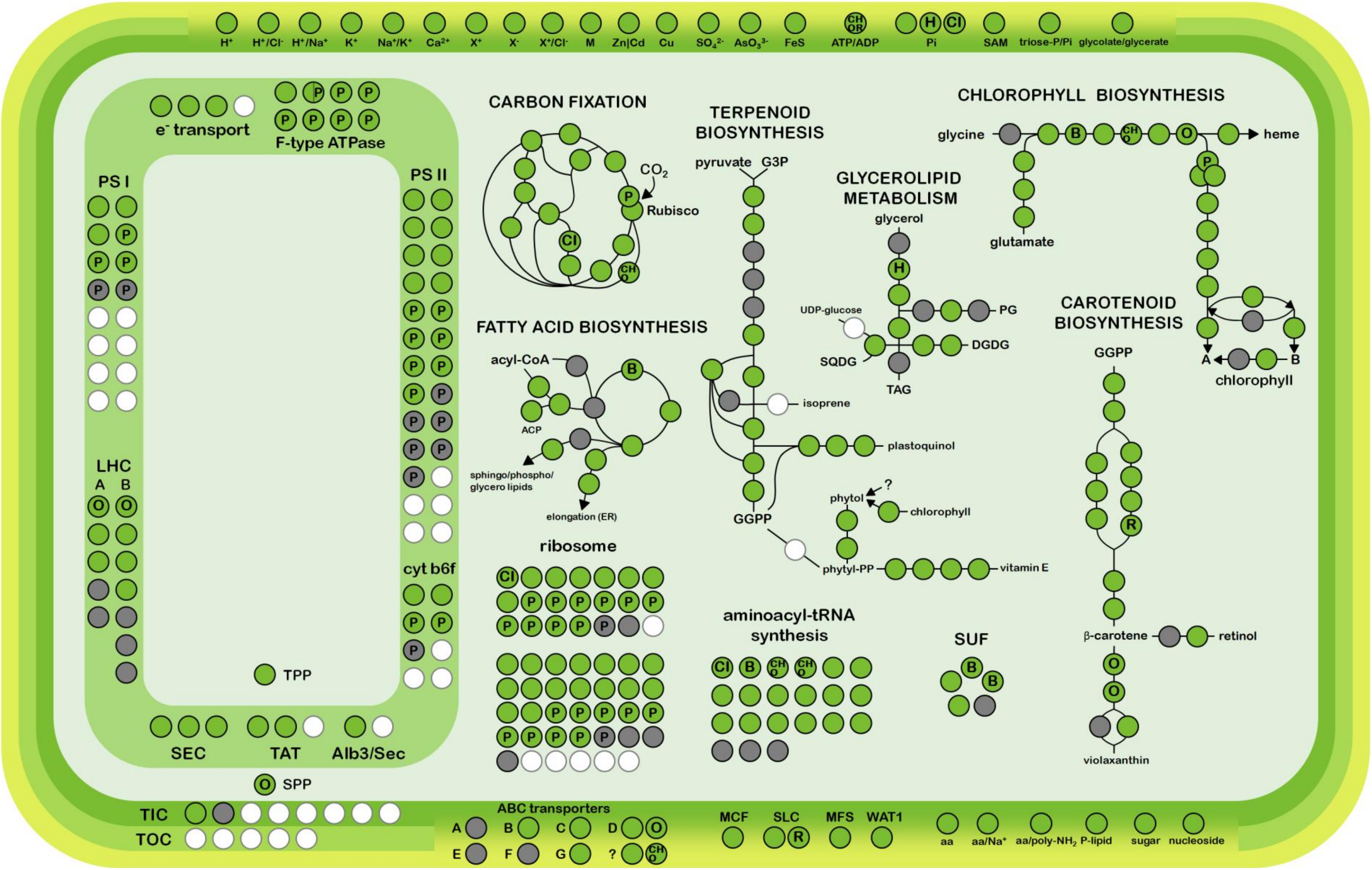


S4: Overview of the *E. gracilis* chloroplast metabolism as reconstructed from mass spectrometry-based proteome: Enzymes present in the plastid proteome in at least one isoform are marked as green circles, grey circles represent enzymes which were identified on the RNA or DNA level (in this study or previously) but are absent from the proteome; white circles represent genes completely absent in Euglena; circles marked by the letter "P" represent genes coded in the plastid genome while rest of the circles represent genes coded in the nucleus; circles marked by other letters represent genes with at least one of their isoforms gained via lateral transfer from one of the following donor groups: "Ch" for chlorarachniophytes, "Cr" for cryptophytes, "H" for haptophytes, "O" for ochrophytes, "R" for rhodophytes, "CHO" for unresolved secondary algae (cryptophytes, haptophytes or ochrophytes), CHOR for unresolved primary or secondary red algae (cryptophytes, haptophytes, ochrophytes or rhodophytes), and "B" for bacteria. Genes related to green algae and discobes are not marked, as well as genes with either "green or chlorarachniophyte" or completely unresolved algal origin. Multiple overlapping circles represent multiple subunits of certain enzymes.

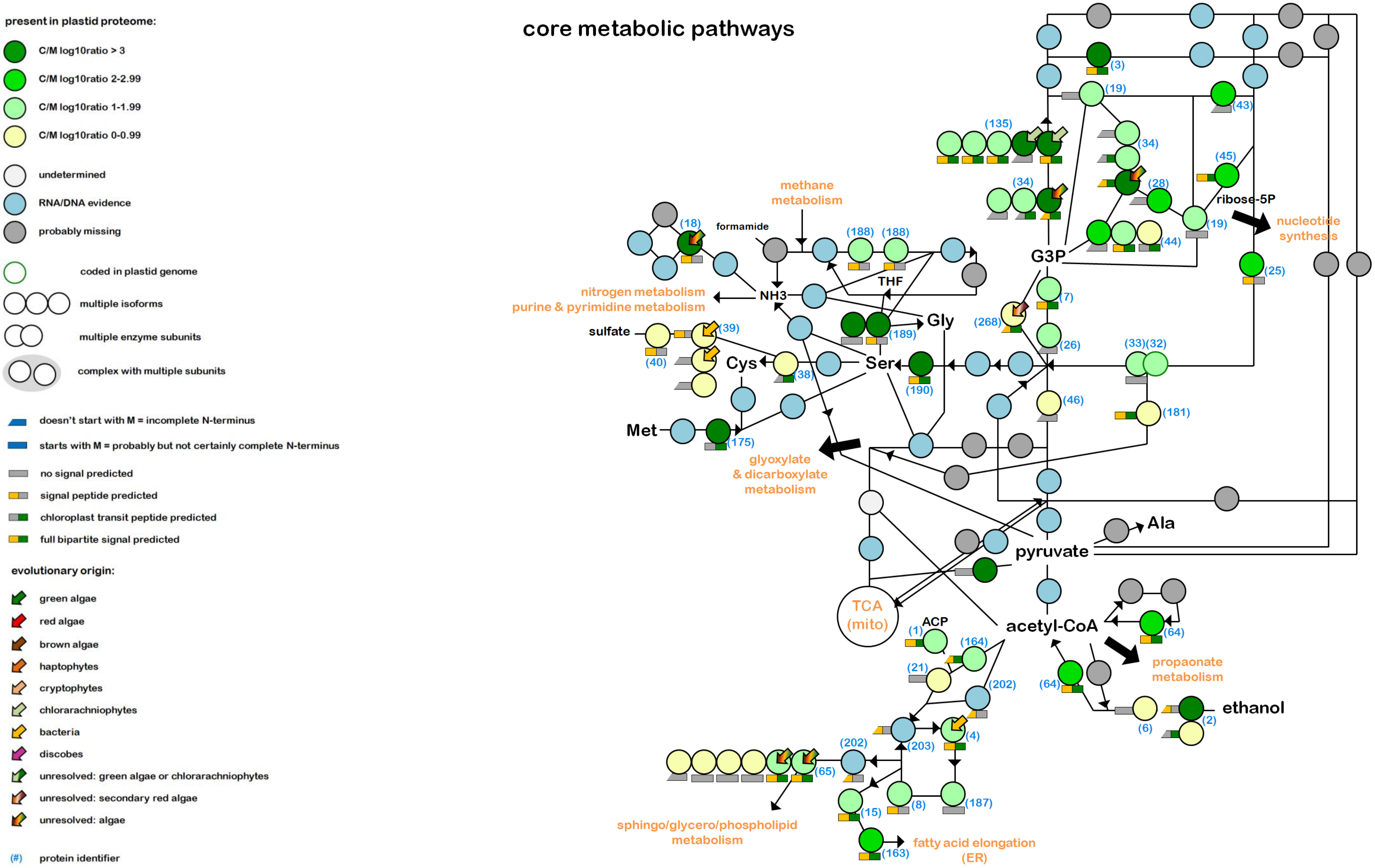


S5.0: Legend for following (S5.1-11) detailed metabolic maps; proteins used in these maps can be searched in supplemental table **supplementary-dataset-1.xlsx** by the identifier in brackets; the identifier is also specified for some proteins with mere DNA/RNA-level evidence which could, however, localize to plastid and fill the gaps in the otherwise plastidial pathways or complexes.
S5.1: Metabolic map of *E. gracilis* chloroplast core metabolic pathways

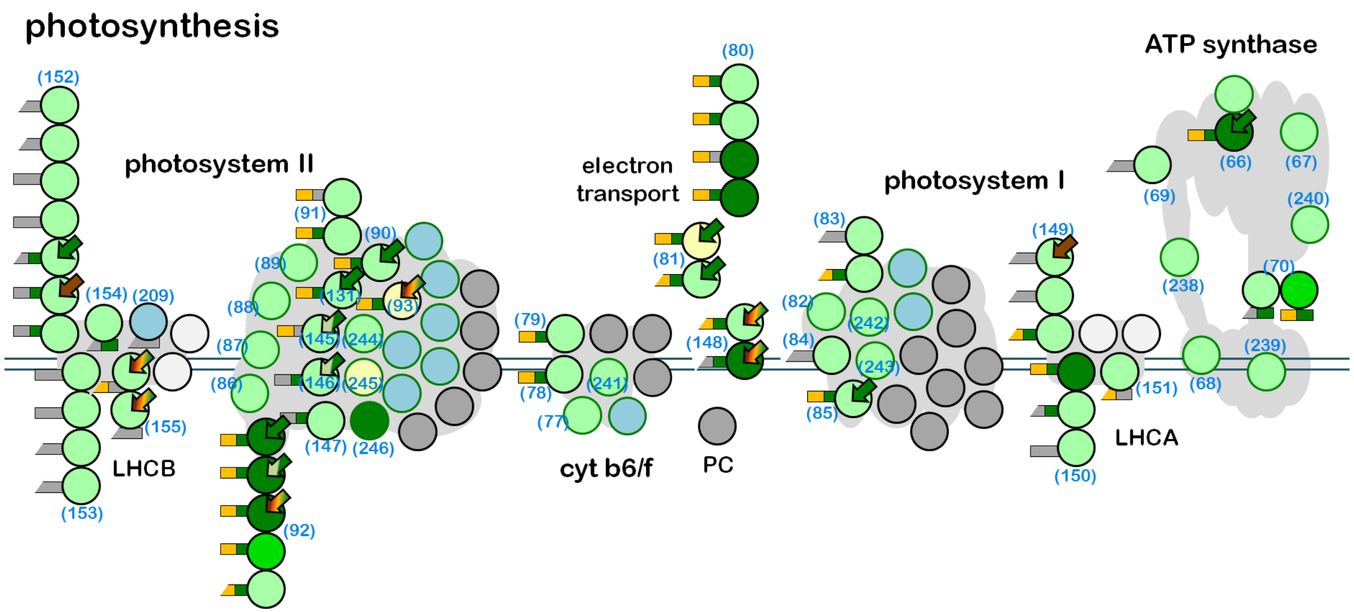


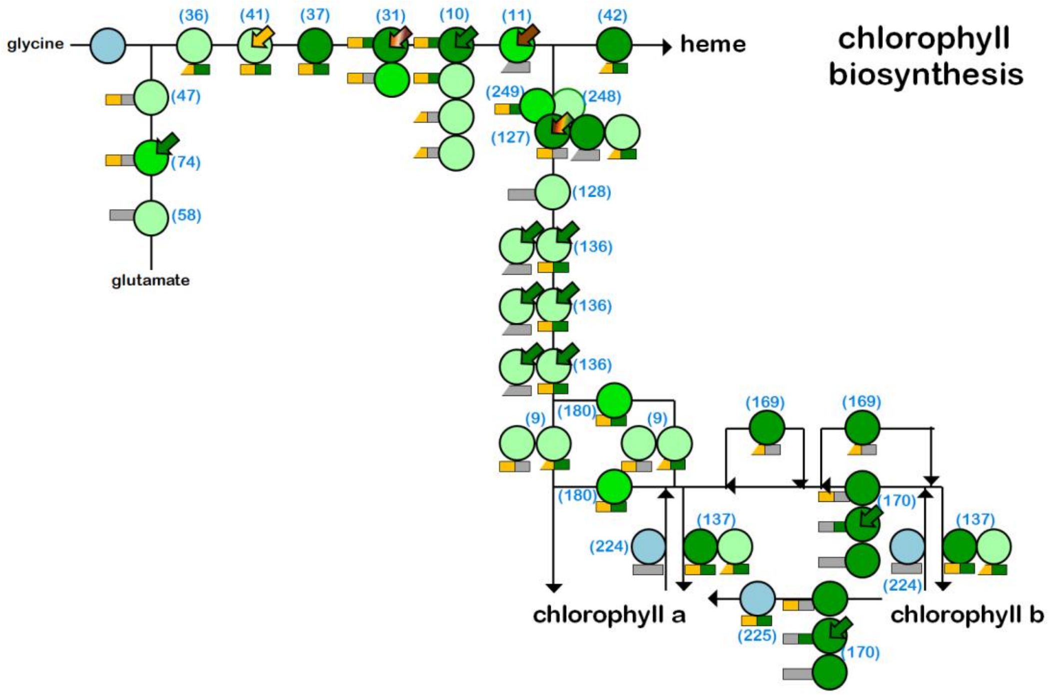


S5.2: Map of *E. gracilis* chloroplast photosynthetic apparatus
S5.3: Metabolic map of *E. gracilis* chloroplast chlorophyll synthesis

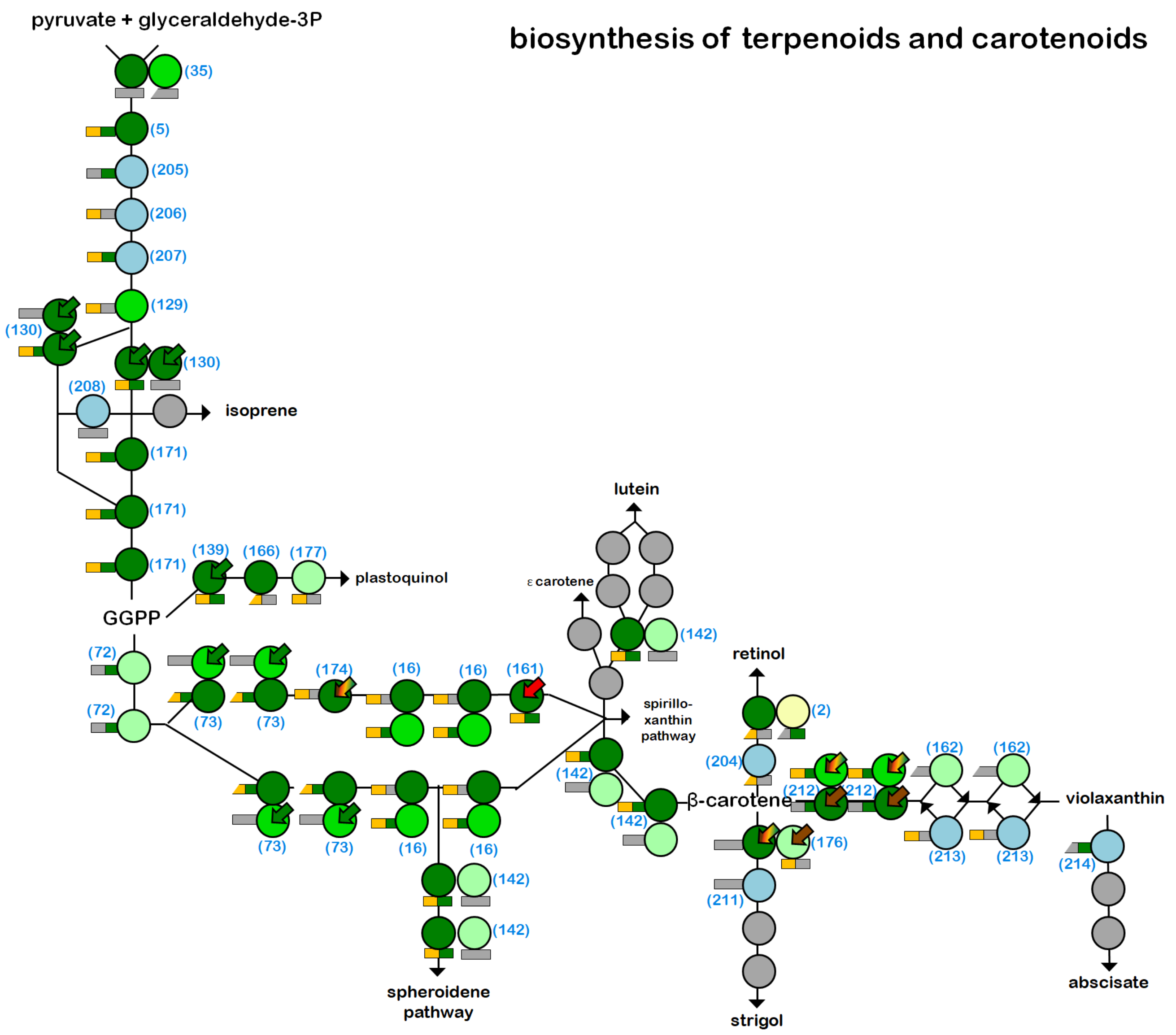


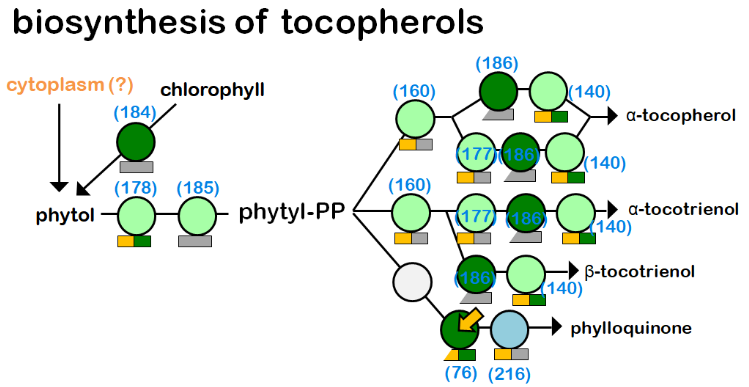


S5.4: Metabolic map of *E. gracilis* chloroplast terpenoid and carotenoid biosynthesis
S5.5: Metabolic map of *E. gracilis* chloroplast tocopherol biosynthesis

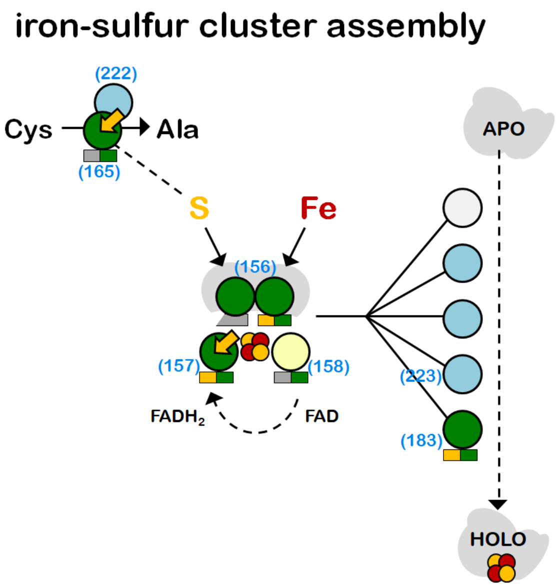


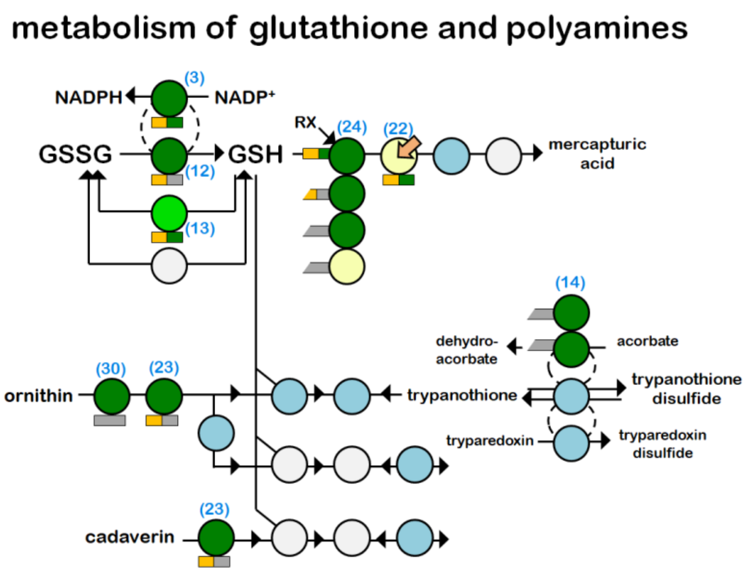


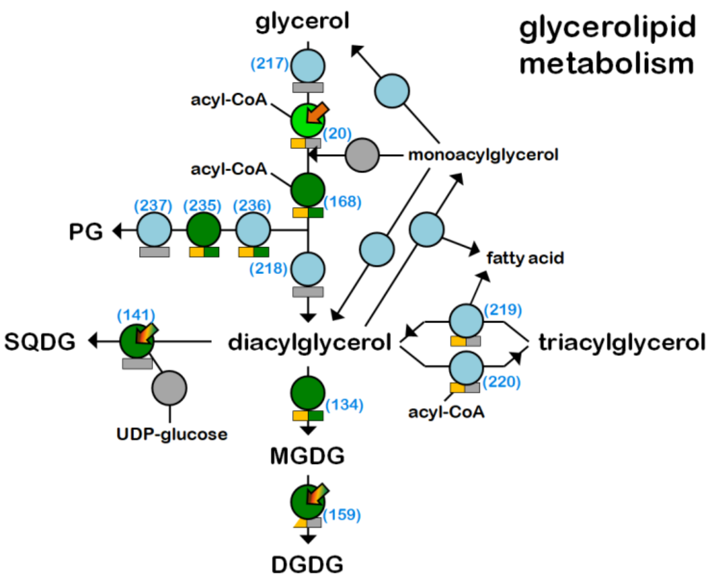


S5.6: Metabolic map of *E. gracilis* chloroplast SUF system
S5.7: Metabolic map of *E. gracilis* chloroplast part of metabolism of glutathione and polyamines
S5.8: Metabolic map of *E. gracilis* chloroplast metabolism of glycerolipids

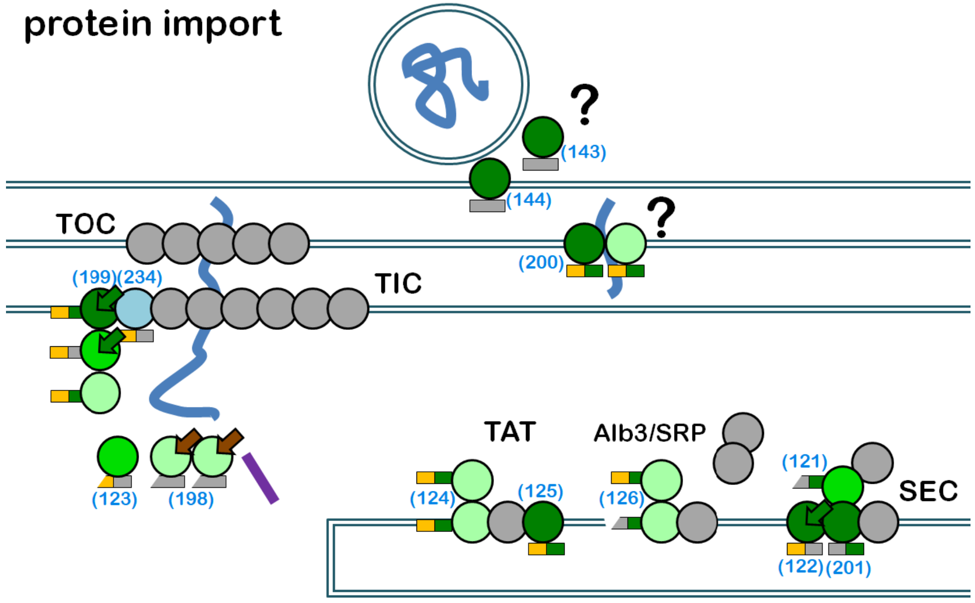


S5.9: Map of *E. gracilis* chloroplast protein importing machinery

### genetic information processing

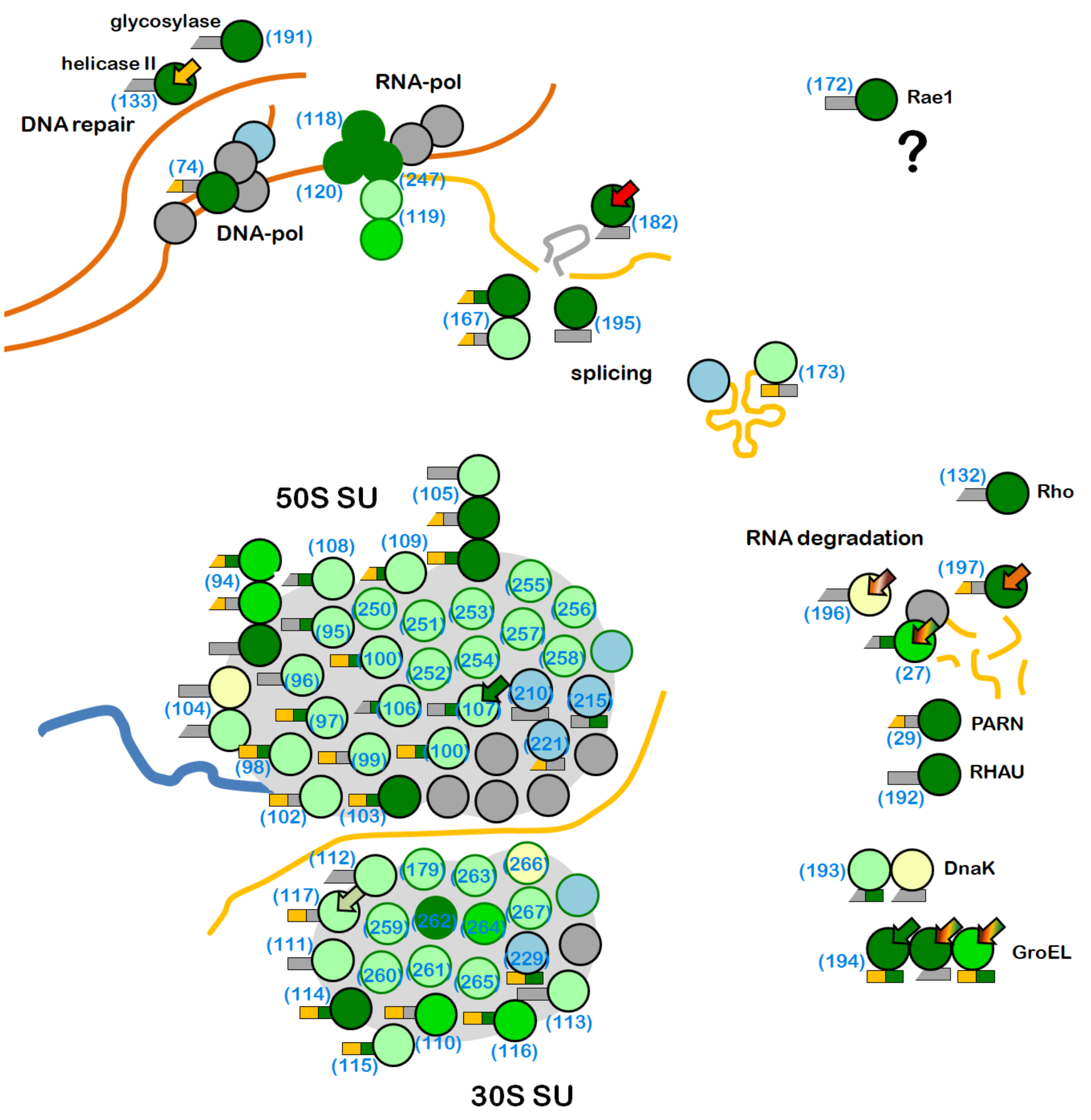


### aminoacyl-tRNA synthesis

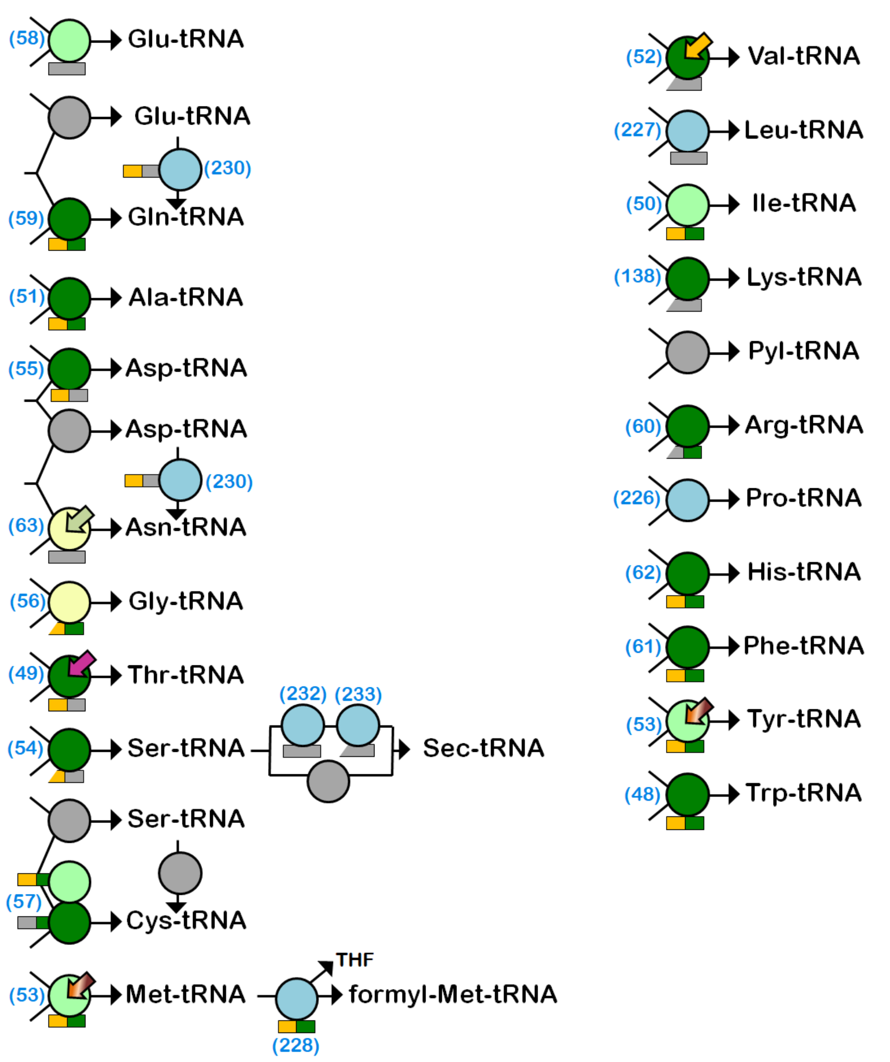


S5.10: Map of *E. gracilis* chloroplast transcription, translation, RNA processing and degradation
S5.11: Map of *E. gracilis* chloroplast aminoacyl-tRNA synthesis

S6: Phylogenetic trees showing positions of the two plastid terminal oxidases (PTOX) identified in transcriptomic data of the three euglenophytes. While the PTOX1 position suggests conventional enzyme inherited from algae, the PTOX2 falls among mitochondrial alternative oxidases.

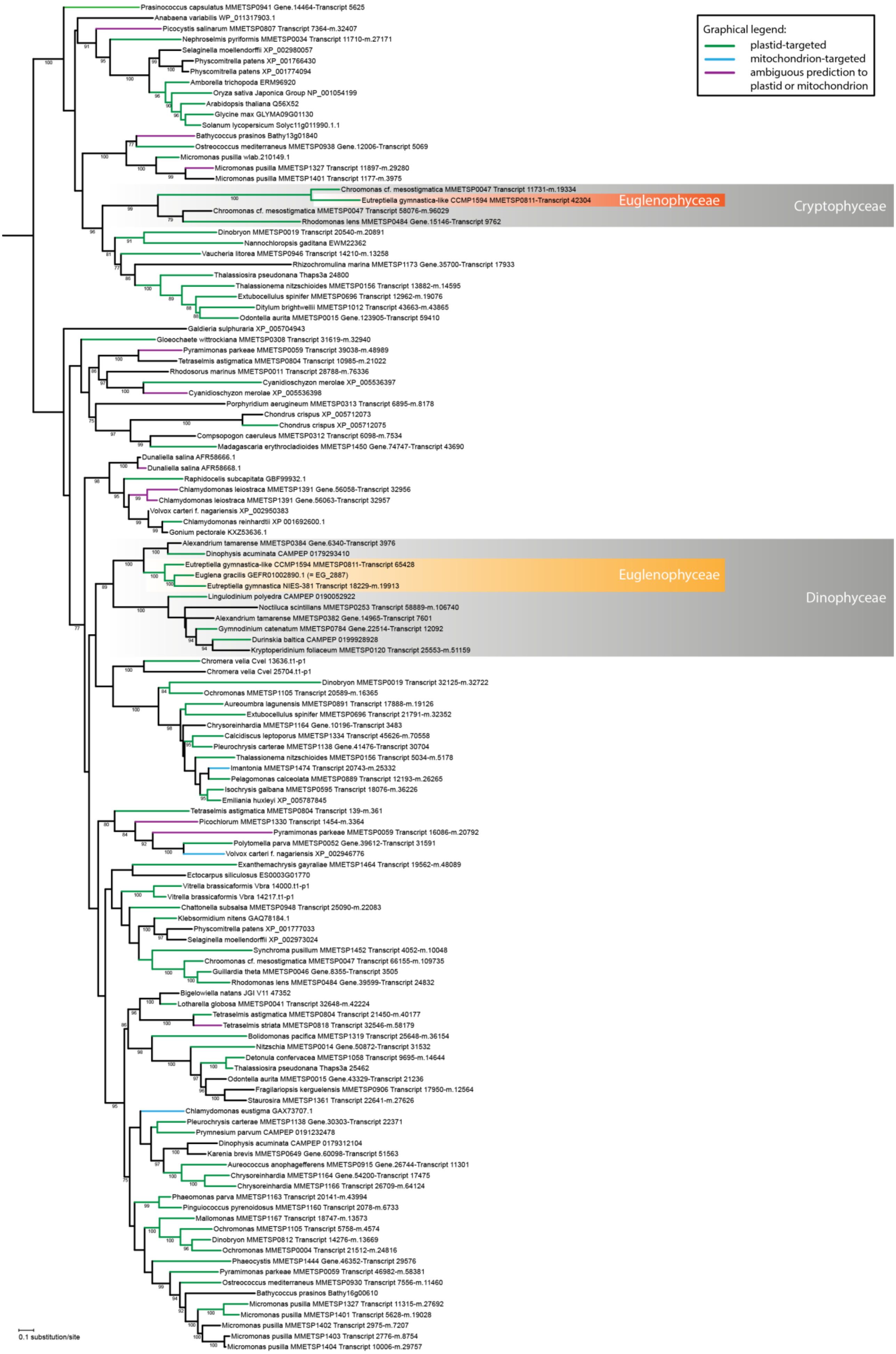

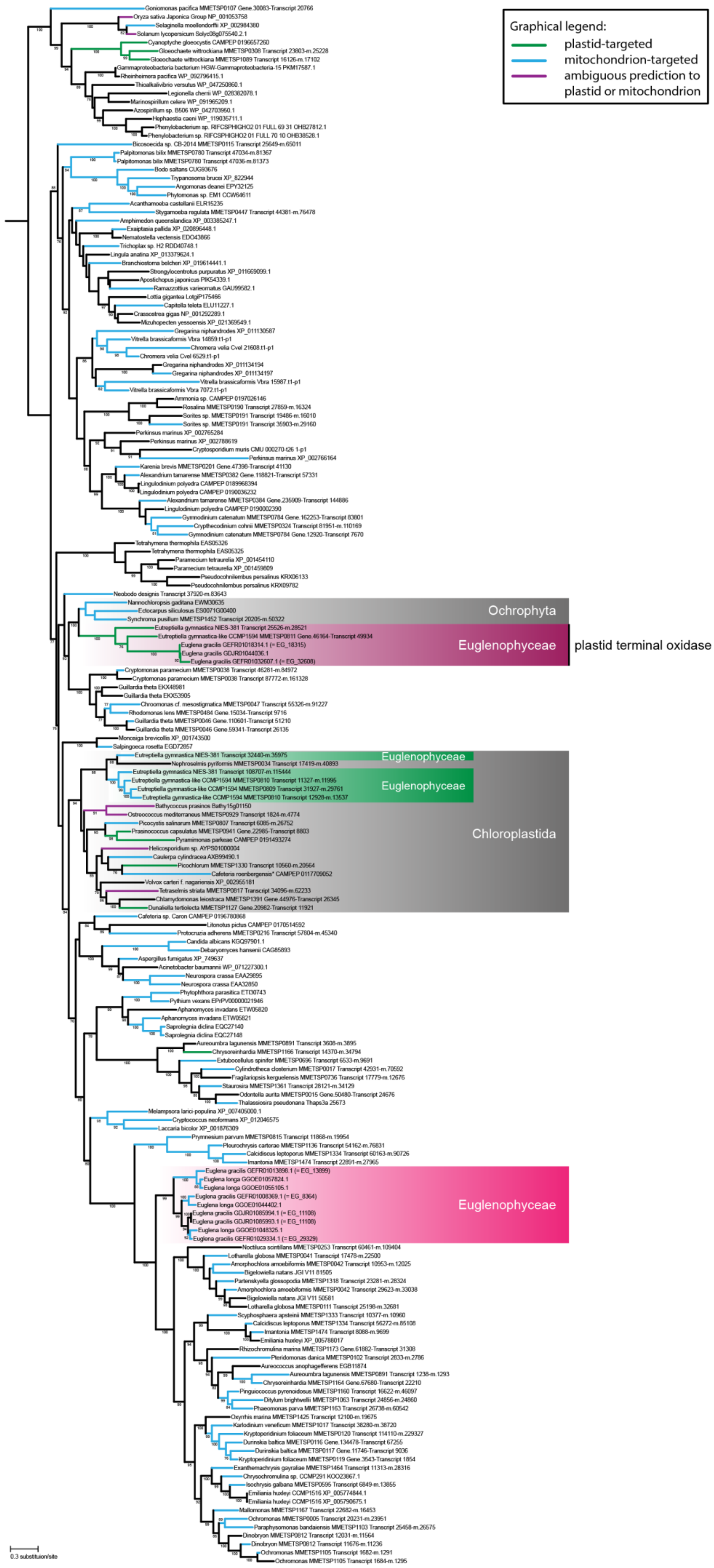

Graphical legend:
plastid-targeted
mitochondrion-targeted
ambiguous prediction to plastid or mitochondrion
Ochrophyta
Euglenophyceae
plastid terminal oxidase
Euglenophyceae
Euglenophyceae
Chloroplastida
Euglenophyceae
0.3 substitution/site

S7: Overview of SUF subunits identified in transcriptomes of *Euglena gracilis*, *Euglena longa* and *Eutreptiella gymnastica* with their sources and accession numbers and proteins corresponding to the *E. gracilis* transcripts with their respective log10 CP/MT ratios representing the credibility of plastidal localization. In case of SufB1, SufE1 and SufS1, the protein was captured in one replicate of mass spectrometry analysis of the plastidal fraction only, suggesting the protein is of lower abundance but plastid-localized. In case of SufD1 and SufS2, the protein was not captured by mass spectrometry and its localization was inferred based on the N-terminal signals and/or localizations of their putative interaction partners.

| pathway | annotation | *E. gracilis* protein ID | log10 CP/MT | *E. gracilis* (GenBank) | *Eut. gymnastica NIES-381* (MMETSP) | *Eut. gymnastica-like CCMP1594* (MMETSP) | *E. longa* (Záhonová *et* Füssy *et al.* 2018) |
|---|---|---|---|---|---|---|---|
| SUF1 | SufB1 | 6397 | >0, one replicate only | GDJR01089002 | CAMNT_0000679585 + reads | CAMNT_0046511287 | Contig18694 + Contig38168 + PCR |
| | SufC1 | 13141 | 3+ | GDJR01041264 | CAMNT_0000683911 | CAMNT_0046510015 | Contig4493 + SL-PCR |
| | SufD1 | 6034, 33281 | not captured by MS | GDJR01038170 | MMETSP0039-Transcript_4862 | CAMNT_0046444101 | Contig15035 |
| | SufE1 | 23255 | >0, one replicate only | GDJR01007756 | MMETSP0039-Transcript_116548 | CAMNT_0046518931 | Contig14296 |
| | SufS1 | 12732 | >0, one replicate only | GDJR01047871 | CAMNT_0000708793 corrected + reads | CAMNT_0046452723 | Contig3452 |
| SUF2 | SufB2 | 8044 | 3+ | GEFR01008046 | - | - | Contig15117 |
| | SufC2 | 24338 | 3+ | GDJR01028245 | - | - | Contig63589 |
| | SufD2 | 13381 | 0,750 | GDJR01049034 | MMETSP0039-Transcript_74532 | MMETSP0809-Transcript_58667 + MMETSP0810-Transcript_89787 + MMETSP0811-Transcript_48154 + MMETSP0811-Transcript_48153 + MMETSP0811-Transcript_48158 + reads | Contig36957 |
| | SufE2 | 32911 | not captured by MS | GDJR01012039 | MMETSP0039-Transcript_109175 | - | Contig52519 + reads |
| | SufS2 | 9032 | 3+ | GDJR01072295 | MMETSP0039-Transcript_75235 | CAMNT_0046485773 + CAMNT_0046488253 + CAMNT_0046492131 + reads | Contig31 + Contig43468 |

S8: Phylogenetic trees showing positions of SUF subunits identified in transcriptomic data of the three euglenophytes; the algae-related genes are highlighted in green while the genes of prokaryotic origin are highlighted in purple.

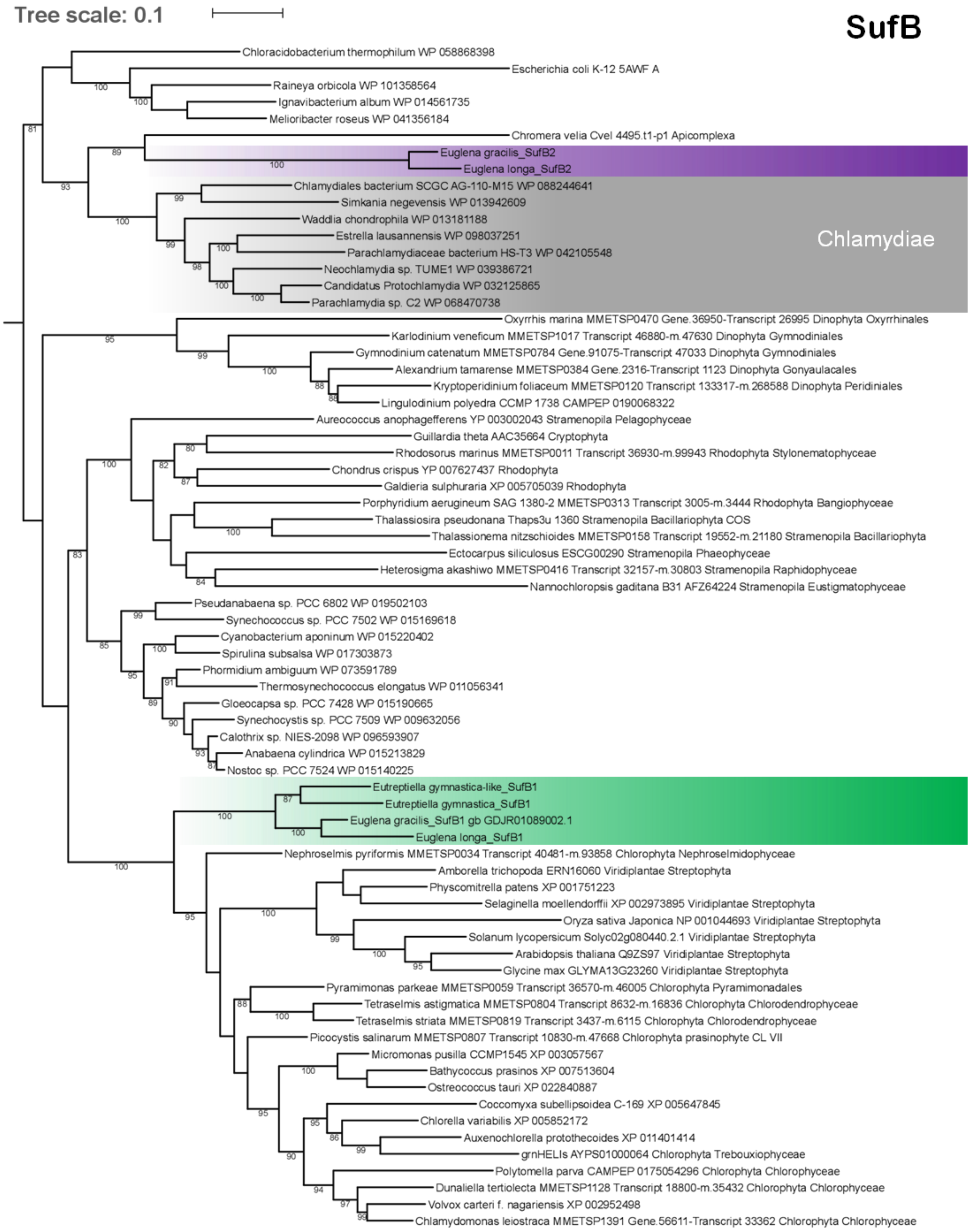


.

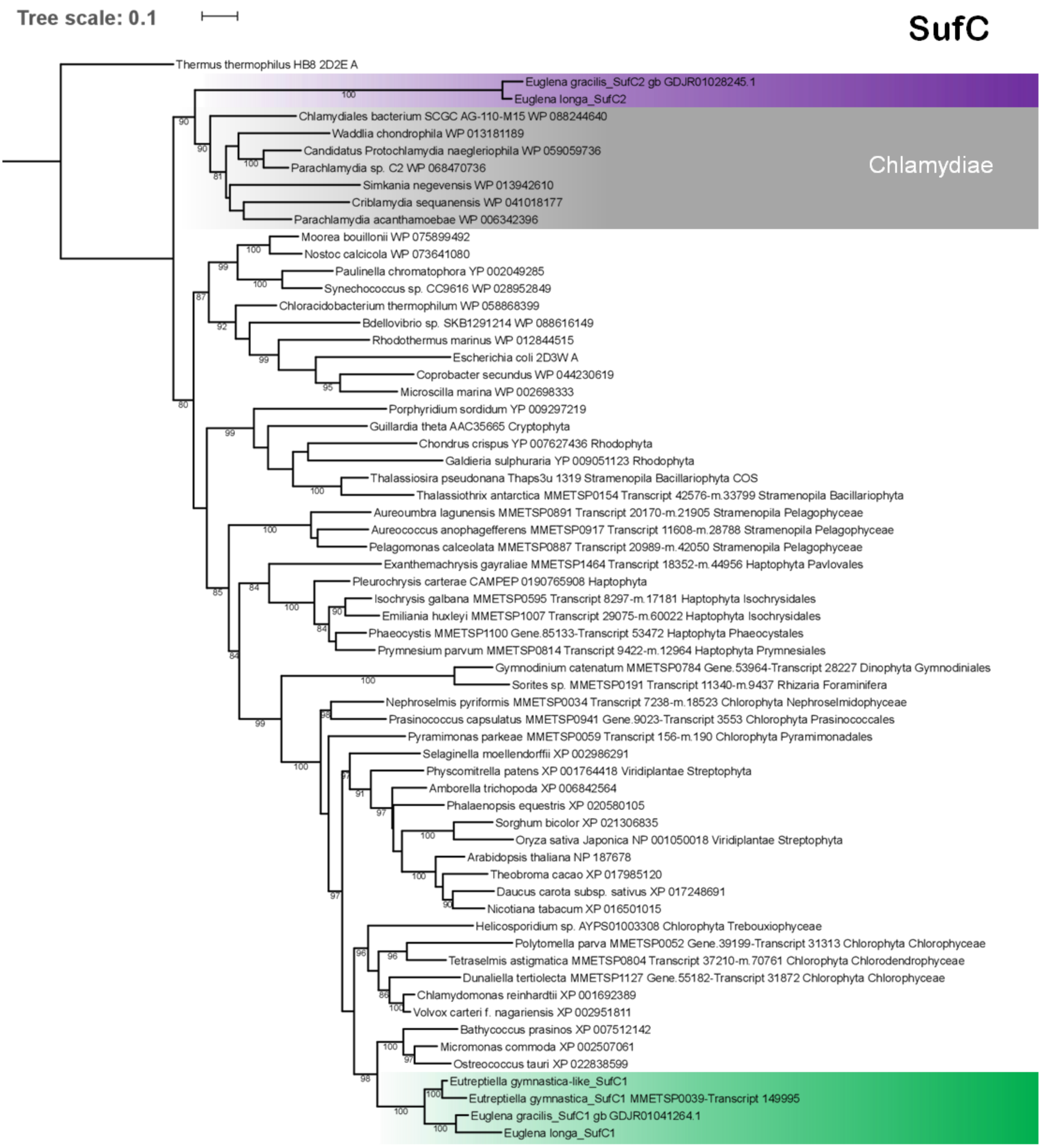

Tree scale: 0.1
SufC
Thermus thermophilus HB8 2D2E A
Euglena gracilis_SufC2 gb GDJR01028245.1
Euglena longa_SufC2
Chlamydiales bacterium SCGC AG-110-M15 WP 088244640
Waddlia chondrophila WP 013181189
Candidatus Protochlamydia naegleriophila WP 059059736
Parachlamydia sp. C2 WP 068470736
Simkania negevensis WP 013942610
Criblamydia sequanensis WP 041018177
Parachlamydia acanthamoebae WP 006342396
Chlamydiae
Moorea bouillonii WP 075899492
Nostoc calcicola WP 073641080
Paulinella chromatophora YP 002049285
Synechococcus sp. CC9616 WP 028952849
Chloracidobacterium thermophilum WP 058868399
Bdellovibrio sp. SKB1291214 WP 088616149
Rhodothermus marinus WP 012844515
Escherichia coli 2D3W A
Coprobacter secundus WP 044230619
Microscilla marina WP 002698333
Porphyridium sordidum YP 009297219
Guillardia theta AAC35665 Cryptophyta
Chondrus crispus YP 007627436 Rhodophyta
Galdieria sulphuraria YP 009051123 Rhodophyta
Thalassiosira pseudonana Thaps3u 1319 Stramenopila Bacillariophyta COS
Thalassiothrix antarctica MMETSP0154 Transcript 42576-m.33799 Stramenopila Bacillariophyta
Aureoumbra lagunensis MMETSP0891 Transcript 20170-m.21905 Stramenopila Pelagophyceae
Aureococcus anophagefferens MMETSP0917 Transcript 11608-m.28788 Stramenopila Pelagophyceae
Pelagomonas calceolata MMETSP0887 Transcript 20989-m.42050 Stramenopila Pelagophyceae
Exanthemachrysis gayraliae MMETSP1464 Transcript 18352-m.44956 Haptophyta Pavlovales
Pleurochrysis carterae CAMPEP 0190765908 Haptophyta
Isochrysis galbana MMETSP0595 Transcript 8297-m.17181 Haptophyta Isochrysidales
Emiliania huxleyi MMETSP1007 Transcript 29075-m.60022 Haptophyta Isochrysidales
Phaeocystis MMETSP1100 Gene.85133-Transcript 53472 Haptophyta Phaeocystales
Prymnesium parvum MMETSP0814 Transcript 9422-m.12964 Haptophyta Prymnesiales
Gymnodinium catenatum MMETSP0784 Gene.53964-Transcript 28227 Dinophyta Gymnodiniales
Sorites sp. MMETSP0191 Transcript 11340-m.9437 Rhizaria Foraminifera
Nephroselmis pyriformis MMETSP0034 Transcript 7238-m.18523 Chlorophyta Nephroselmidophyceae
Prasinococcus capsulatus MMETSP0941 Gene.9023-Transcript 3553 Chlorophyta Prasinococcales
Pyramimonas parkeae MMETSP0059 Transcript 156-m.190 Chlorophyta Pyramimonadales
Selaginella moellendorffii XP 002986291
Physcomitrella patens XP 001764418 Viridiplantae Streptophyta
Amborella trichopoda XP 006842564
Phalaenopsis equestris XP 020580105
Sorghum bicolor XP 021306835
Oryza sativa Japonica NP 001050018 Viridiplantae Streptophyta
Arabidopsis thaliana NP 187678
Theobroma cacao XP 017985120
Daucus carota subsp. sativus XP 017248691
Nicotiana tabacum XP 016501015
Helicosporidium sp. AYPS01003308 Chlorophyta Trebouxiophyceae
Polytomella parva MMETSP0052 Gene.39199-Transcript 31313 Chlorophyta Chlorophyceae
Tetraselmis astigmatica MMETSP0804 Transcript 37210-m.70761 Chlorophyta Chlorodendrophyceae
Dunaliella tertiolecta MMETSP1127 Gene.55182-Transcript 31872 Chlorophyta Chlorophyceae
Chlamydomonas reinhardtii XP 001692389
Volvox carteri f. nagariensis XP 002951811
Bathycoccus prasinos XP 007512142
Micromonas commoda XP 002507061
Ostreococcus tauri XP 022838599
Eutreptiella gymnastica-like_SufC1
Eutreptiella gymnastica_SufC1 MMETSP0039-Transcript 149995
Euglena gracilis_SufC1 gb GDJR01041264.1
Euglena longa_SufC1

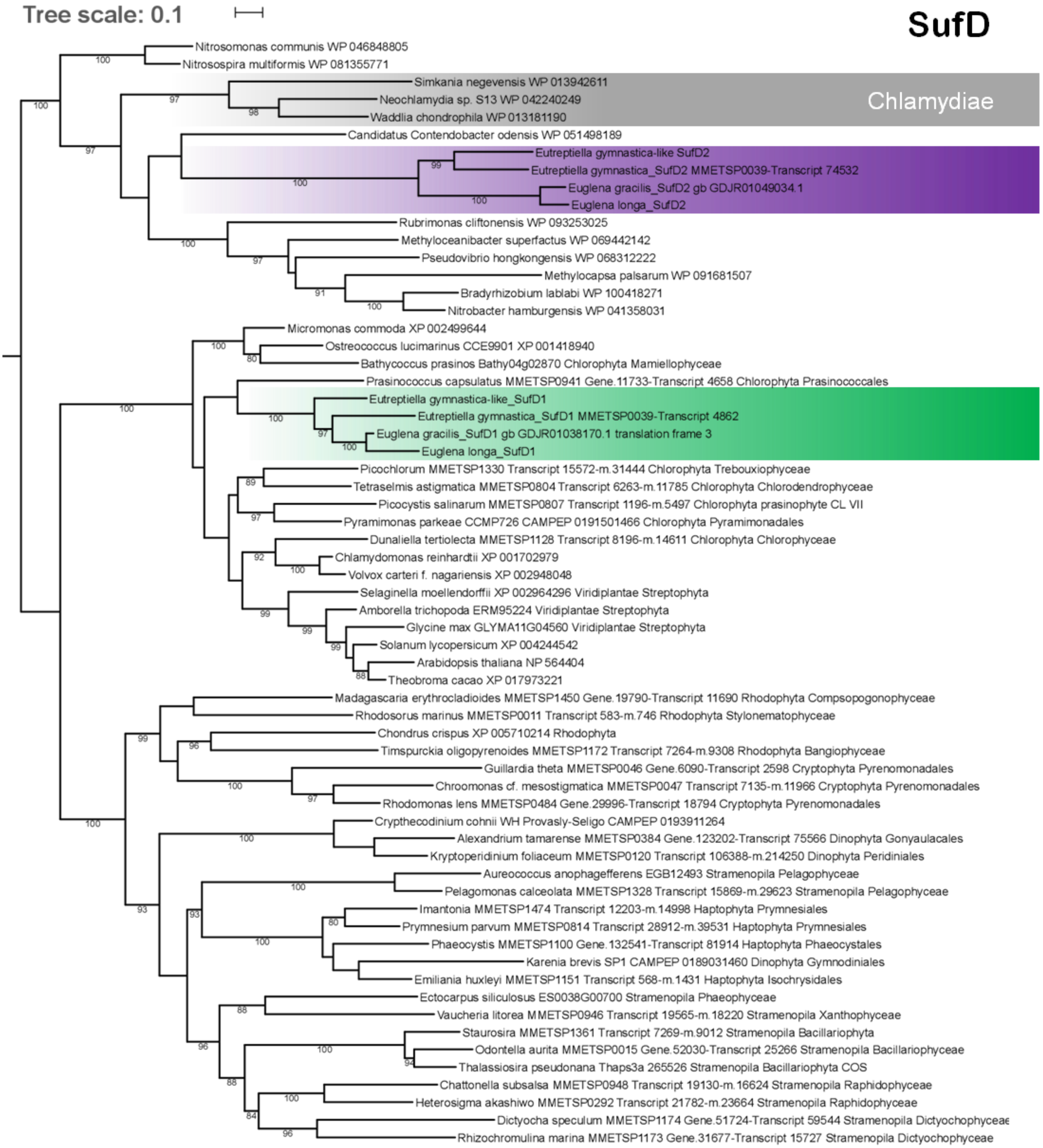

Tree scale: 0.1
SufD
Chlamydiae
Nitrosomonas communis WP 046848805
Nitrosospira multiformis WP 081355771
Simkania negevensis WP 013942611
Neochlamydia sp. S13 WP 042240249
Waddlia chondrophila WP 013181190
Candidatus Contendobacter odensis WP 051498189
Eutreptiella gymnastica-like SufD2
Eutreptiella gymnastica_SufD2 MMETSP0039-Transcript 74532
Euglena gracilis_SufD2 gb GDJR01049034.1
Euglena longa_SufD2
Rubrimonas cliftonensis WP 093253025
Methyloceanibacter superfactus WP 069442142
Pseudovibrio hongkongensis WP 068312222
Methylocapsa palsarum WP 091681507
Bradyrhizobium lablabi WP 100418271
Nitrobacter hamburgensis WP 041358031
Micromonas commoda XP 002499644
Ostreococcus lucimarinus CCE9901 XP 001418940
Bathycoccus prasinos Bathy04g02870 Chlorophyta Mamiellophyceae
Prasinococcus capsulatus MMETSP0941 Gene.11733-Transcript 4658 Chlorophyta Prasinococcales
Eutreptiella gymnastica-like_SufD1
Eutreptiella gymnastica_SufD1 MMETSP0039-Transcript 4862
Euglena gracilis_SufD1 gb GDJR01038170.1 translation frame 3
Euglena longa_SufD1
Picochlorum MMETSP1330 Transcript 15572-m.31444 Chlorophyta Trebouxiophyceae
Tetraselmis astigmatica MMETSP0804 Transcript 6263-m.11785 Chlorophyta Chlorodendrophyceae
Picocystis salinarum MMETSP0807 Transcript 1196-m.5497 Chlorophyta prasinophyte CL VII
Pyramimonas parkeae CCMP726 CAMPEP 0191501466 Chlorophyta Pyramimonadales
Dunaliella tertiolecta MMETSP1128 Transcript 8196-m.14611 Chlorophyta Chlorophyceae
Chlamydomonas reinhardtii XP 001702979
Volvox carteri f. nagariensis XP 002948048
Selaginella moellendorffii XP 002964296 Viridiplantae Streptophyta
Amborella trichopoda ERM95224 Viridiplantae Streptophyta
Glycine max GLYMA11G04560 Viridiplantae Streptophyta
Solanum lycopersicum XP 004244542
Arabidopsis thaliana NP 564404
Theobroma cacao XP 017973221
Madagascaria erythrocladioides MMETSP1450 Gene.19790-Transcript 11690 Rhodophyta Compsopogonophyceae
Rhodosorus marinus MMETSP0011 Transcript 583-m.746 Rhodophyta Stylonematophyceae
Chondrus crispus XP 005710214 Rhodophyta
Timspurckia oligopyrenoides MMETSP1172 Transcript 7264-m.9308 Rhodophyta Bangiophyceae
Guillardia theta MMETSP0046 Gene.6090-Transcript 2598 Cryptophyta Pyrenomonadales
Chroomonas cf. mesostigmatica MMETSP0047 Transcript 7135-m.11966 Cryptophyta Pyrenomonadales
Rhodomonas lens MMETSP0484 Gene.29996-Transcript 18794 Cryptophyta Pyrenomonadales
Crypthecodinium cohnii WH Provasly-Seligo CAMPEP 0193911264
Alexandrium tamarense MMETSP0384 Gene.123202-Transcript 75566 Dinophyta Gonyaulacales
Kryptoperidinium foliaceum MMETSP0120 Transcript 106388-m.214250 Dinophyta Peridiniales
Aureococcus anophagefferens EGB12493 Stramenopila Pelagophyceae
Pelagomonas calceolata MMETSP1328 Transcript 15869-m.29623 Stramenopila Pelagophyceae
Imantonia MMETSP1474 Transcript 12203-m.14998 Haptophyta Prymnesiales
Prymnesium parvum MMETSP0814 Transcript 28912-m.39531 Haptophyta Prymnesiales
Phaeocystis MMETSP1100 Gene.132541-Transcript 81914 Haptophyta Phaeocystales
Karenia brevis SP1 CAMPEP 0189031460 Dinophyta Gymnodiniales
Emiliania huxleyi MMETSP1151 Transcript 568-m.1431 Haptophyta Isochrysidales
Ectocarpus siliculosus ES0038G00700 Stramenopila Phaeophyceae
Vaucheria litorea MMETSP0946 Transcript 19565-m.18220 Stramenopila Xanthophyceae
Staurosira MMETSP1361 Transcript 7269-m.9012 Stramenopila Bacillariophyta
Odontella aurita MMETSP0015 Gene.52030-Transcript 25266 Stramenopila Bacillariophyceae
Thalassiosira pseudonana Thaps3a 265526 Stramenopila Bacillariophyta COS
Chattonella subsalsa MMETSP0948 Transcript 19130-m.16624 Stramenopila Raphidophyceae
Heterosigma akashiwo MMETSP0292 Transcript 21782-m.23664 Stramenopila Raphidophyceae
Dictyocha speculum MMETSP1174 Gene.51724-Transcript 59544 Stramenopila Dictyochophyceae
Rhizochromulina marina MMETSP1173 Gene.31677-Transcript 15727 Stramenopila Dictyochophyceae

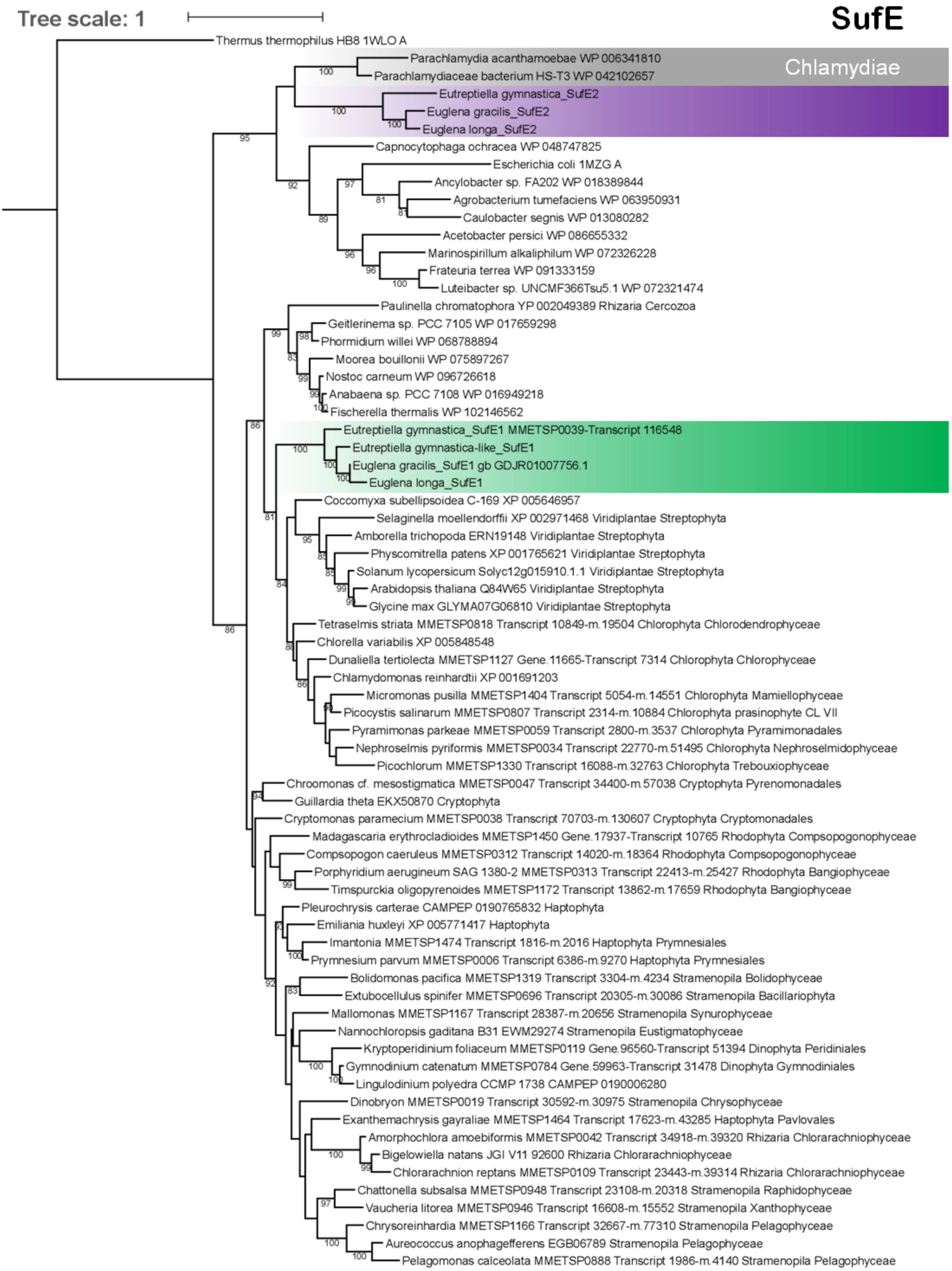
Tree scale: 1
SufE
Chlamydiae
Thermus thermophilus HB8 1WLO A
Parachlamydia acanthamoebae WP 006341810
Parachlamydiaceae bacterium HS-T3 WP 042102657
Eutreptiella gymnastica_SufE2
Euglena gracilis_SufE2
Euglena longa_SufE2
Capnocytophaga ochracea WP 048747825
Escherichia coli 1MZG A
Ancylobacter sp. FA202 WP 018389844
Agrobacterium tumefaciens WP 063950931
Caulobacter segnis WP 013080282
Acetobacter persici WP 086655332
Marinospirillum alkaliphilum WP 072326228
Frateuria terrea WP 091333159
Luteibacter sp. UNCMF366Tsu5.1 WP 072321474
Paulinella chromatophora YP 002049389 Rhizaria Cercozoa
Geitlerinema sp. PCC 7105 WP 017659298
Phormidium willei WP 068788894
Moorea bouillonii WP 075897267
Nostoc carneum WP 096726618
Anabaena sp. PCC 7108 WP 016949218
Fischerella thermalis WP 102146562
Eutreptiella gymnastica_SufE1 MMETSP0039-Transcript 116548
Eutreptiella gymnastica-like_SufE1
Euglena gracilis_SufE1 gb GDJR01007756.1
Euglena longa_SufE1
Coccomyxa subellipsoidea C-169 XP 005646957
Selaginella moellendorffii XP 002971468 Viridiplantae Streptophyta
Amborella trichopoda ERN19148 Viridiplantae Streptophyta
Physcomitrella patens XP 001765621 Viridiplantae Streptophyta
Solanum lycopersicum Solyc12g015910.1.1 Viridiplantae Streptophyta
Arabidopsis thaliana Q84W65 Viridiplantae Streptophyta
Glycine max GLYMA07G06810 Viridiplantae Streptophyta
Tetraselmis striata MMETSP0818 Transcript 10849-m.19504 Chlorophyta Chlorodendrophyceae
Chlorella variabilis XP 005848548
Dunaliella tertiolecta MMETSP1127 Gene.11665-Transcript 7314 Chlorophyta Chlorophyceae
Chlamydomonas reinhardtii XP 001691203
Micromonas pusilla MMETSP1404 Transcript 5054-m.14551 Chlorophyta Mamiellophyceae
Picocystis salinarum MMETSP0807 Transcript 2314-m.10884 Chlorophyta prasinophyte CL VII
Pyramimonas parkeae MMETSP0059 Transcript 2800-m.3537 Chlorophyta Pyramimonadales
Nephroselmis pyriformis MMETSP0034 Transcript 22770-m.51495 Chlorophyta Nephroselmidophyceae
Picochlorum MMETSP1330 Transcript 16088-m.32763 Chlorophyta Trebouxiophyceae
Chroomonas cf. mesostigmatica MMETSP0047 Transcript 34400-m.57038 Cryptophyta Pyrenomonadales
Guillardia theta EKX50870 Cryptophyta
Cryptomonas paramecium MMETSP0038 Transcript 70703-m.130607 Cryptophyta Cryptomonadales
Madagascaria erythrocladioides MMETSP1450 Gene.17937-Transcript 10765 Rhodophyta Compsopogonophyceae
Compsopogon caeruleus MMETSP0312 Transcript 14020-m.18364 Rhodophyta Compsopogonophyceae
Porphyridium aerugineum SAG 1380-2 MMETSP0313 Transcript 22413-m.25427 Rhodophyta Bangiophyceae
Timspurckia oligopyrenoides MMETSP1172 Transcript 13862-m.17659 Rhodophyta Bangiophyceae
Pleurochrysis carterae CAMPEP 0190765832 Haptophyta
Emiliania huxleyi XP 005771417 Haptophyta
Imantonia MMETSP1474 Transcript 1816-m.2016 Haptophyta Prymnesiales
Prymnesium parvum MMETSP0006 Transcript 6386-m.9270 Haptophyta Prymnesiales
Bolidomonas pacifica MMETSP1319 Transcript 3304-m.4234 Stramenopila Bolidophyceae
Extubocellulus spinifer MMETSP0696 Transcript 20305-m.30086 Stramenopila Bacillariophyta
Mallomonas MMETSP1167 Transcript 28387-m.20656 Stramenopila Synurophyceae
Nannochloropsis gaditana B31 EWM29274 Stramenopila Eustigmatophyceae
Kryptoperidinium foliaceum MMETSP0119 Gene.96560-Transcript 51394 Dinophyta Peridiniales
Gymnodinium catenatum MMETSP0784 Gene.59963-Transcript 31478 Dinophyta Gymnodiniales
Lingulodinium polyedra CCMP 1738 CAMPEP 0190006280
Dinobryon MMETSP0019 Transcript 30592-m.30975 Stramenopila Chrysophyceae
Exanthemachrysis gayraliae MMETSP1464 Transcript 17623-m.43285 Haptophyta Pavlovales
Amorphochlora amoebiformis MMETSP0042 Transcript 34918-m.39320 Rhizaria Chlorarachniophyceae
Bigelowiella natans JGI V11 92600 Rhizaria Chlorarachniophyceae
Chlorarachnion reptans MMETSP0109 Transcript 23443-m.39314 Rhizaria Chlorarachniophyceae
Chattonella subsalsa MMETSP0948 Transcript 23108-m.20318 Stramenopila Raphidophyceae
Vaucheria litorea MMETSP0946 Transcript 16608-m.15552 Stramenopila Xanthophyceae
Chrysoreinhardia MMETSP1166 Transcript 32667-m.77310 Stramenopila Pelagophyceae
Aureococcus anophagefferens EGB06789 Stramenopila Pelagophyceae
Pelagomonas calceolata MMETSP0888 Transcript 1986-m.4140 Stramenopila Pelagophyceae

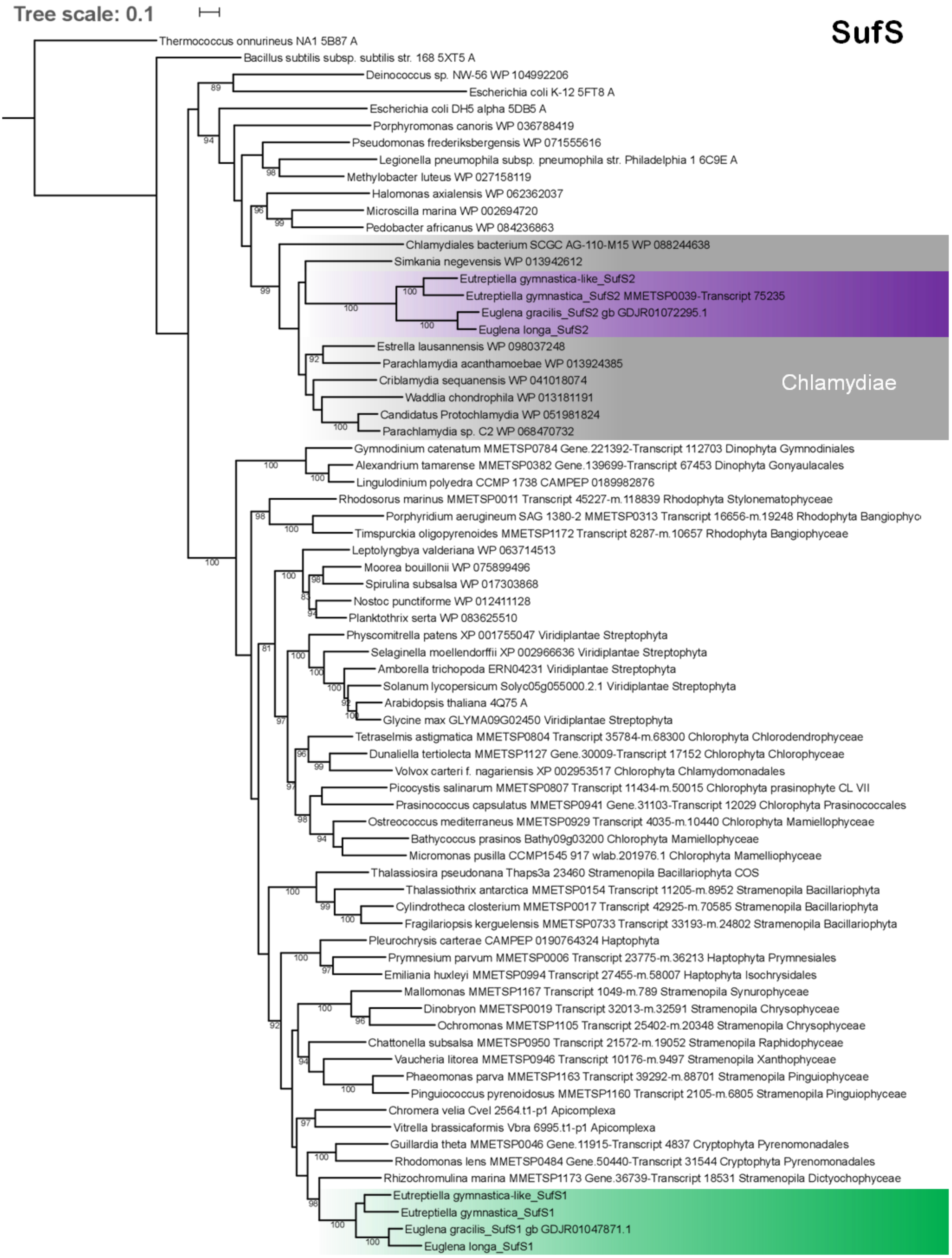
Tree scale: 0.1
SufS
Thermococcus onnurineus NA1 5B87 A
Bacillus subtilis subsp. subtilis str. 168 5XT5 A
Deinococcus sp. NW-56 WP 104992206
Escherichia coli K-12 5FT8 A
Escherichia coli DH5 alpha 5DB5 A
Porphyromonas canoris WP 036788419
Pseudomonas frederiksbergensis WP 071555616
Legionella pneumophila subsp. pneumophila str. Philadelphia 1 6C9E A
Methylobacter luteus WP 027158119
Halomonas axialensis WP 062362037
Microscilla marina WP 002694720
Pedobacter africanus WP 084236863
Chlamydiales bacterium SCGC AG-110-M15 WP 088244638
Simkania negevensis WP 013942612
Eutreptiella gymnastica-like_SufS2
Eutreptiella gymnastica_SufS2 MMETSP0039-Transcript 75235
Euglena gracilis_SufS2 gb GDJR01072295.1
Euglena longa_SufS2
Estrella lausannensis WP 098037248
Parachlamydia acanthamoebae WP 013924385
Criblamydia sequanensis WP 041018074
Waddlia chondrophila WP 013181191
Candidatus Protochlamydia WP 051981824
Parachlamydia sp. C2 WP 068470732
Chlamydiae
Gymnodinium catenatum MMETSP0784 Gene.221392-Transcript 112703 Dinophyta Gymnodiniales
Alexandrium tamarense MMETSP0382 Gene.139699-Transcript 67453 Dinophyta Gonyaulacales
Lingulodinium polyedra CCMP 1738 CAMPEP 0189982876
Rhodosorus marinus MMETSP0011 Transcript 45227-m.118839 Rhodophyta Stylonematophyceae
Porphyridium aerugineum SAG 1380-2 MMETSP0313 Transcript 16656-m.19248 Rhodophyta Bangiophyc
Timspurckia oligopyrenoides MMETSP1172 Transcript 8287-m.10657 Rhodophyta Bangiophyceae
Leptolyngbya valderiana WP 063714513
Moorea bouillonii WP 075899496
Spirulina subsalsa WP 017303868
Nostoc punctiforme WP 012411128
Planktothrix serta WP 083625510
Physcomitrella patens XP 001755047 Viridiplantae Streptophyta
Selaginella moellendorffii XP 002966636 Viridiplantae Streptophyta
Amborella trichopoda ERN04231 Viridiplantae Streptophyta
Solanum lycopersicum Solyc05g055000.2.1 Viridiplantae Streptophyta
Arabidopsis thaliana 4Q75 A
Glycine max GLYMA09G02450 Viridiplantae Streptophyta
Tetraselmis astigmatica MMETSP0804 Transcript 35784-m.68300 Chlorophyta Chlorodendrophyceae
Dunaliella tertiolecta MMETSP1127 Gene.30009-Transcript 17152 Chlorophyta Chlorophyceae
Volvox carteri f. nagariensis XP 002953517 Chlorophyta Chlamydomonadales
Picocystis salinarum MMETSP0807 Transcript 11434-m.50015 Chlorophyta prasinophyte CL VII
Prasinococcus capsulatus MMETSP0941 Gene.31103-Transcript 12029 Chlorophyta Prasinococcales
Ostreococcus mediterraneus MMETSP0929 Transcript 4035-m.10440 Chlorophyta Mamiellophyceae
Bathycoccus prasinos Bathy09g03200 Chlorophyta Mamiellophyceae
Micromonas pusilla CCMP1545 917 wlab.201976.1 Chlorophyta Mamelliophyceae
Thalassiosira pseudonana Thaps3a 23460 Stramenopila Bacillariophyta COS
Thalassiothrix antarctica MMETSP0154 Transcript 11205-m.8952 Stramenopila Bacillariophyta
Cylindrotheca closterium MMETSP0017 Transcript 42925-m.70585 Stramenopila Bacillariophyta
Fragilariopsis kerguelensis MMETSP0733 Transcript 33193-m.24802 Stramenopila Bacillariophyta
Pleurochrysis carterae CAMPEP 0190764324 Haptophyta
Prymnesium parvum MMETSP0006 Transcript 23775-m.36213 Haptophyta Prymnesiales
Emiliania huxleyi MMETSP0994 Transcript 27455-m.58007 Haptophyta Isochrysidales
Mallomonas MMETSP1167 Transcript 1049-m.789 Stramenopila Synurophyceae
Dinobryon MMETSP0019 Transcript 32013-m.32591 Stramenopila Chrysophyceae
Ochromonas MMETSP1105 Transcript 25402-m.20348 Stramenopila Chrysophyceae
Chattonella subsalsa MMETSP0950 Transcript 21572-m.19052 Stramenopila Raphidophyceae
Vaucheria litorea MMETSP0946 Transcript 10176-m.9497 Stramenopila Xanthophyceae
Phaeomonas parva MMETSP1163 Transcript 39292-m.88701 Stramenopila Pinguiophyceae
Pinguiococcus pyrenoidosus MMETSP1160 Transcript 2105-m.6805 Stramenopila Pinguiophyceae
Chromera velia Cvel 2564.t1-p1 Apicomplexa
Vitrella brassicaformis Vbra 6995.t1-p1 Apicomplexa
Guillardia theta MMETSP0046 Gene.11915-Transcript 4837 Cryptophyta Pyrenomonadales
Rhodomonas lens MMETSP0484 Gene.50440-Transcript 31544 Cryptophyta Pyrenomonadales
Rhizochromulina marina MMETSP1173 Gene.36739-Transcript 18531 Stramenopila Dictyochophyceae
Eutreptiella gymnastica-like_SufS1
Eutreptiella gymnastica_SufS1
Euglena gracilis_SufS1 gb GDJR01047871.1
Euglena longa_SufS1

S9: Phylogenetic tree showing position of plastidial ferredoxin among chlamydiae.

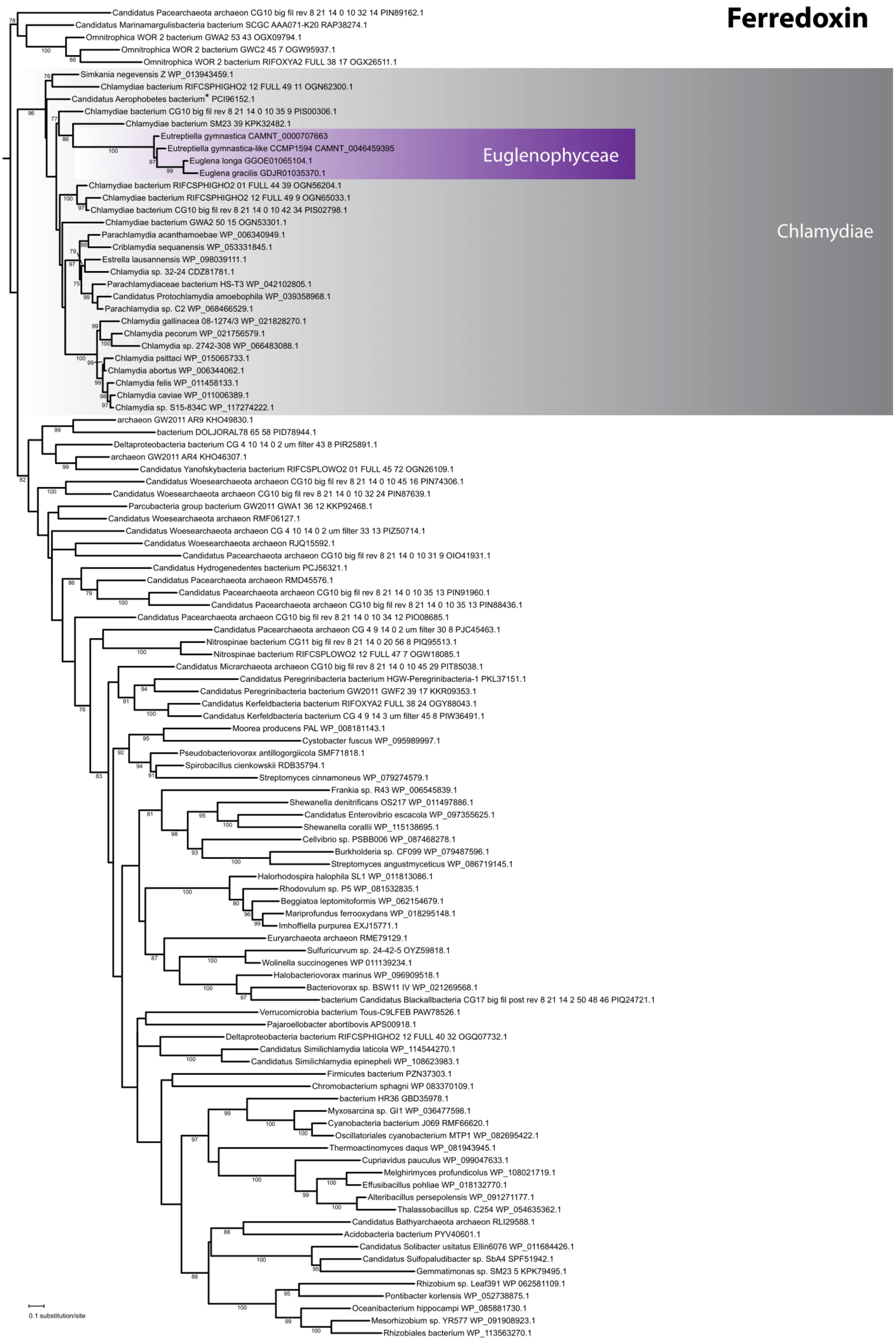

S10: Phylogenetic trees showing positions of sulfite reductase subunits. While the subunit alpha (CysJ) sits inside chlamydiae, the subunit beta (CysI), which is also of putative prokaryotic evolutionary origin, is sister to spirochaetes.

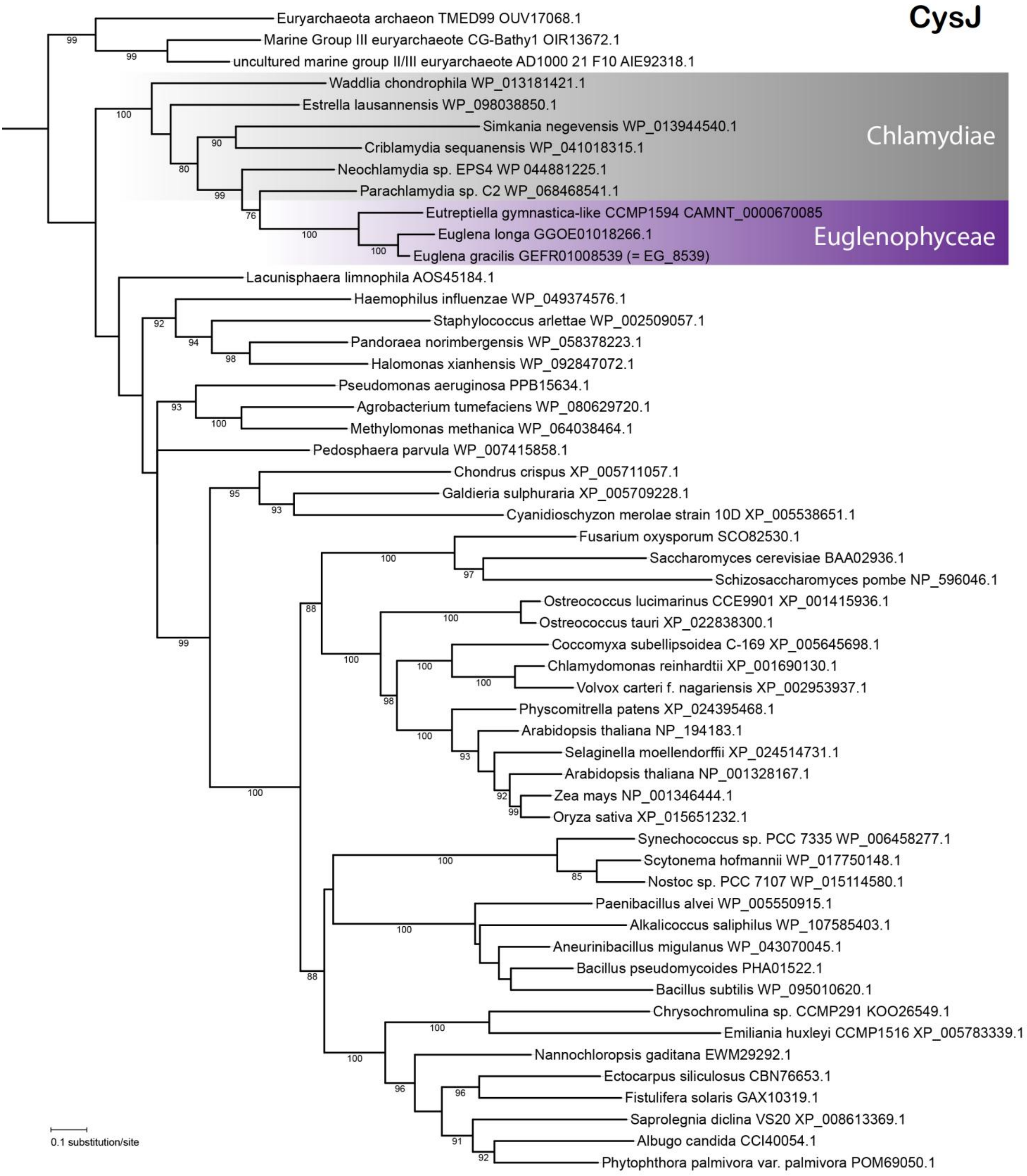

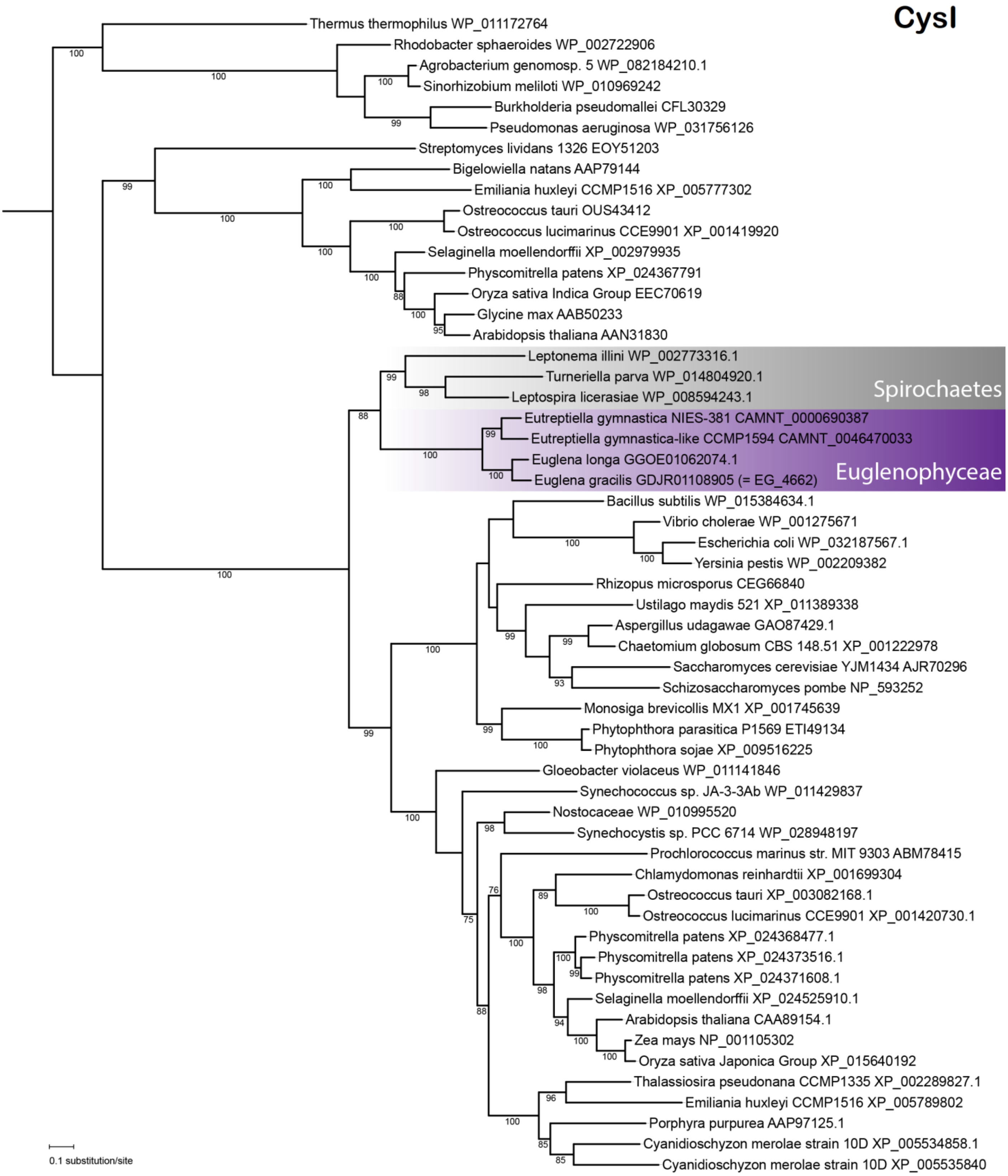
CysI
Thermus thermophilus WP_011172764
Rhodobacter sphaeroides WP_002722906
Agrobacterium genomosp. 5 WP_082184210.1
Sinorhizobium meliloti WP_010969242
Burkholderia pseudomallei CFL30329
Pseudomonas aeruginosa WP_031756126
Streptomyces lividans 1326 EOY51203
Bigelowiella natans AAP79144
Emiliania huxleyi CCMP1516 XP_005777302
Ostreococcus tauri OUS43412
Ostreococcus lucimarinus CCE9901 XP_001419920
Selaginella moellendorffii XP_002979935
Physcomitrella patens XP_024367791
Oryza sativa Indica Group EEC70619
Glycine max AAB50233
Arabidopsis thaliana AAN31830
Leptonema illini WP_002773316.1
Turneriella parva WP_014804920.1
Leptospira licerasiae WP_008594243.1
Spirochaetes
Eutreptiella gymnastica NIES-381 CAMNT_0000690387
Eutreptiella gymnastica-like CCMP1594 CAMNT_0046470033
Euglena longa GGOE01062074.1
Euglena gracilis GDJR01108905 (= EG_4662)
Euglenophyceae
Bacillus subtilis WP_015384634.1
Vibrio cholerae WP_001275671
Escherichia coli WP_032187567.1
Yersinia pestis WP_002209382
Rhizopus microsporus CEG66840
Ustilago maydis 521 XP_011389338
Aspergillus udagawae GAO87429.1
Chaetomium globosum CBS 148.51 XP_001222978
Saccharomyces cerevisiae YJM1434 AJR70296
Schizosaccharomyces pombe NP_593252
Monosiga brevicollis MX1 XP_001745639
Phytophthora parasitica P1569 ETI49134
Phytophthora sojae XP_009516225
Gloeobacter violaceus WP_011141846
Synechococcus sp. JA-3-3Ab WP_011429837
Nostocaceae WP_010995520
Synechocystis sp. PCC 6714 WP_028948197
Prochlorococcus marinus str. MIT 9303 ABM78415
Chlamydomonas reinhardtii XP_001699304
Ostreococcus tauri XP_003082168.1
Ostreococcus lucimarinus CCE9901 XP_001420730.1
Physcomitrella patens XP_024368477.1
Physcomitrella patens XP_024373516.1
Physcomitrella patens XP_024371608.1
Selaginella moellendorffii XP_024525910.1
Arabidopsis thaliana CAA89154.1
Zea mays NP_001105302
Oryza sativa Japonica Group XP_015640192
Thalassiosira pseudonana CCMP1335 XP_002289827.1
Emiliania huxleyi CCMP1516 XP_005789802
Porphyra purpurea AAP97125.1
Cyanidioschyzon merolae strain 10D XP_005534858.1
Cyanidioschyzon merolae strain 10D XP_005535840
0.1 substitution/site

S11: MAFFT alignment of the conserved region (positions 310-650 of the fasta alignment derlins.fasta provided in supplementary files) of Der-like plastid proteins of *E. gracilis*, *E. longa*, *Eutreptiella gymnastica*, and *Eutreptiella sp.*, reffered to as DerL1 and DerL2, host (ER) and symbiont (SELMA) specific Der1 proteins of *Guillardia theta*, *Phaeodactylum tricornutum*, and *Plasmodium falciparum* (as identified in Sommer et al. 2007), Der of a red alga *Gracilariopsis chorda*, and three human derlins and a human RHBDL4 rhomboid-like protease. In comparison to the more canonical derlins, the euglenophyte homologs contain approximately 200 aa long N-terminal extension not shown in this view. Conserved positions are highlighted in light and dark grey.

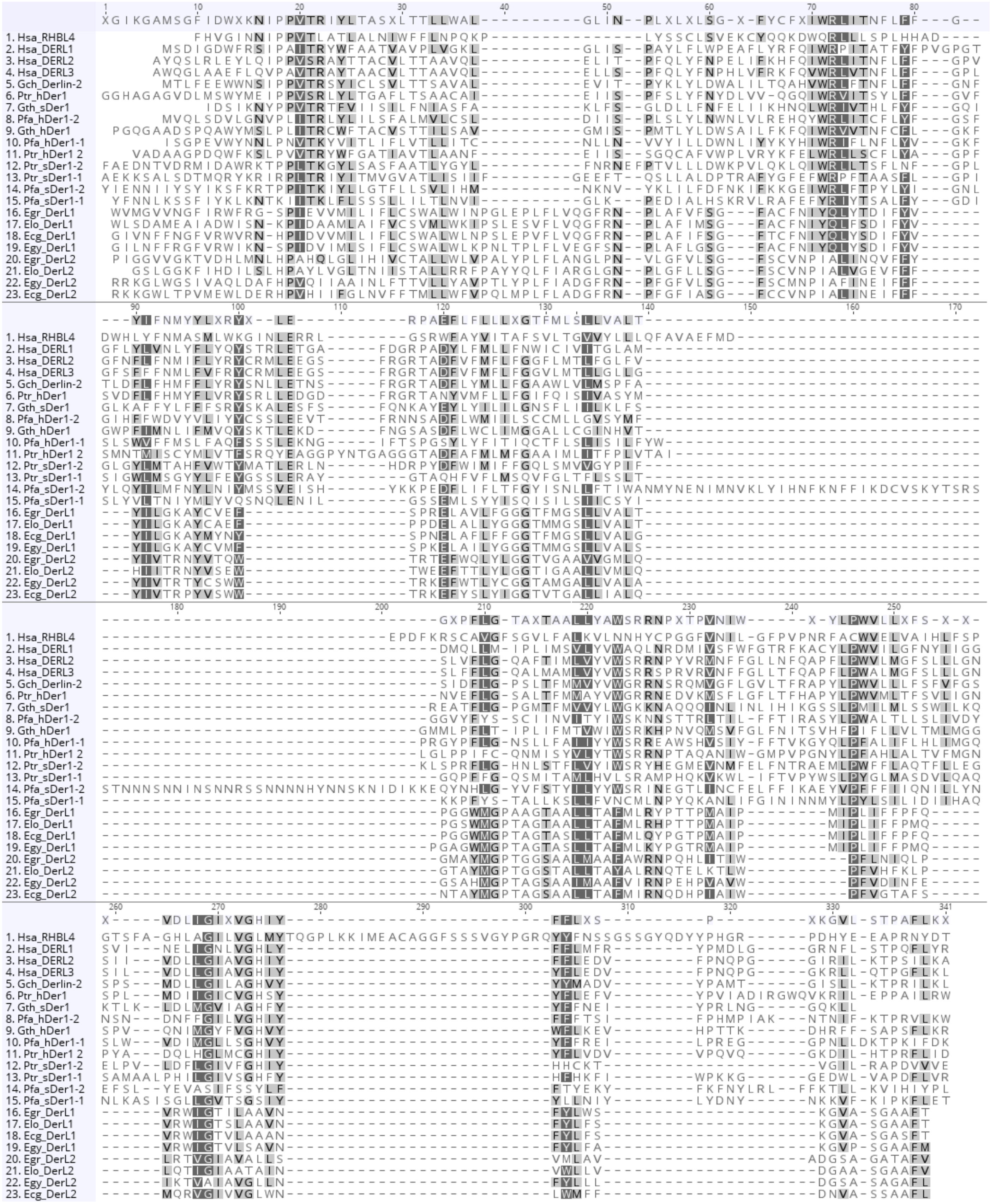

**SAMPLE**

| ALL | chisq sum | p-val chisq sum | norm chisq sum | p-val norm chisq sum | CLASS I | chisq sum | p-val chisq sum | norm chisq sum | p-val norm chisq sum | CLASS II | chisq sum | p-val chisq sum | norm chisq sum | p-val norm chisq sum | no class | chisq sum | p-val chisq sum | norm chisq sum | p-val norm chisq sum |
|---|---|---|---|---|---|---|---|---|---|---|---|---|---|---|---|---|---|---|---|
| R | 175,086 | 1 | 53,1541 | 0,00605 | R | 68,4805 | 1 | 5,5466 | 0,67413 | R | 76,4798 | 1 | 44,0918 | 0,00019 | R | 30,1258 | 0,99969 | 3,51573 | 0,65261 |
| H | 196,786 | 1 | 100,417 | 1,59E-07 | H | 103,726 | 0,99999 | 62,2853 | 2,04E-06 | H | 68,4743 | 1 | 32,7265 | 0,00485 | H | 24,5851 | 0,99999 | 5,40481 | 0,48533 |
| K | 295,634 | 0,99906 | -165,47 | 0 | K | 165,277 | 0,66994 | -125,249 | 0 | K | 97,0751 | 0,99779 | -41,4843 | 0,00045 | K | 33,2821 | 0,99854 | 1,26311 | 0,87152 |
| D | 270,683 | 0,99999 | -168,934 | 0 | D | 146,398 | 0,93696 | -118,104 | 0 | D | 74,2336 | 1 | -38,8899 | 0,00101 | D | 50,0515 | 0,84044 | -11,9406 | 0,1263 |
| E | 331,85 | 0,94694 | -150,778 | 6,88E-15 | E | 181,6 | 0,33097 | -127,026 | 0 | E | 91,39 | 0,9995 | -18,2782 | 0,1224 | E | 58,8598 | 0,55389 | -5,47392 | 0,48339 |
| S | 401,66 | 0,16465 | 206,083 | 0 | S | 232,689 | 0,00196 | 132,049 | 0 | S | 124,561 | 0,82088 | 57,6567 | 1,10E-06 | S | 44,4105 | 0,94553 | 16,377 | 0,03601 |
| T | 220,009 | 1 | 90,9184 | 2,67E-06 | T | 139,829 | 0,97327 | 71,7089 | 5,44E-08 | T | 49,0246 | 1 | 13,2613 | 0,26238 | T | 31,1553 | 0,99947 | 5,94825 | 0,4463 |
| N | 127,755 | 1 | -42,4131 | 0,0283 | N | 62,6668 | 1 | -23,0973 | 0,07908 | N | 42,1703 | 1 | -23,2282 | 0,04963 | N | 22,9175 | 1 | 3,91239 | 0,61642 |
| Q | 171,625 | 1 | 31,1712 | 0,10747 | Q | 109,334 | 0,99997 | 28,1758 | 0,03268 | Q | 44,4741 | 1 | 2,79381 | 0,81334 | Q | 17,8174 | 1 | 0,20158 | 0,97941 |
| A | 550,223 | 8,93E-09 | 207,823 | 0 | A | 322,218 | 6,09E-11 | 130,78 | 0 | A | 141,913 | 0,43895 | 50,5491 | 1,94E-05 | A | 86,0922 | 0,01889 | 26,4941 | 0,00069 |
| I | 198,963 | 1 | -116,336 | 1,79E-09 | I | 94,6478 | 1 | -60,7521 | 4,11E-06 | I | 75,0626 | 1 | -47,7654 | 5,09E-05 | I | 29,2522 | 0,99981 | -7,81817 | 0,31682 |
| L | 193,851 | 1 | -87,2043 | 6,69E-06 | L | 84,6792 | 1 | -43,2203 | 0,00105 | L | 69,5416 | 1 | -25,8399 | 0,02897 | L | 39,6304 | 0,98461 | -18,1441 | 0,02017 |
| V | 197,749 | 1 | -44,6475 | 0,02113 | V | 77,5955 | 1 | 4,64804 | 0,72456 | V | 80,1055 | 0,99999 | -36,5342 | 0,00202 | V | 40,0481 | 0,98254 | -12,7614 | 0,10227 |
| F | 208,752 | 1 | -134,126 | 4,32E-12 | F | 101,429 | 1 | -64,428 | 1,04E-06 | F | 84,0705 | 0,99995 | -54,9889 | 3,36E-06 | F | 23,253 | 1 | -14,7091 | 0,05966 |
| W | 78,9335 | 1 | -0,15093 | 0,9937 | W | 33,1091 | 1 | -0,69202 | 0,95755 | W | 32,2305 | 1 | -2,38569 | 0,83849 | W | 13,5939 | 1 | 2,92677 | 0,70318 |
| Y | 131,868 | 1 | -101,205 | 1,43E-07 | Y | 60,5727 | 1 | -56,6235 | 1,58E-05 | Y | 41,1337 | 1 | -31,9114 | 0,0066 | Y | 30,1611 | 0,99954 | -12,6704 | 0,10189 |
| M | 106,69 | 1 | -74,8029 | 9,86E-05 | M | 37,7605 | 1 | -27,7899 | 0,03357 | M | 45,2825 | 1 | -36,2452 | 0,00196 | M | 23,6471 | 1 | -10,7678 | 0,168 |
| C | 69,7508 | 1 | -45,7416 | 0,01298 | C | 22,9606 | 1 | -27,2205 | 0,03354 | C | 27,8388 | 1 | -12,5845 | 0,24666 | C | 18,9514 | 1 | -5,93654 | 0,43168 |
| G | 203,731 | 1 | -55,8315 | 0,00394 | G | 97,7821 | 1 | -32,365 | 0,01414 | G | 64,0528 | 1 | -21,6138 | 0,06775 | G | 41,8965 | 0,97065 | -1,85269 | 0,81249 |
| P | 794,957 | 0 | 331,381 | 0 | P | 425,55 | 0 | 193,94 | 0 | P | 300,299 | 1,00E-13 | 122,179 | 0 | P | 69,1068 | 0,22258 | 15,2626 | 0,05068 |

**CONTROL**

| ALL | chisq sum | p-val chisq sum | norm chisq sum | p-val norm chisq sum | CLASS I | chisq sum | p-val chisq sum | norm chisq sum | p-val norm chisq sum | CLASS II | chisq sum | p-val chisq sum | norm chisq sum | p-val norm chisq sum | no class | chisq sum | p-val chisq sum | norm chisq sum | p-val norm chisq sum |
|---|---|---|---|---|---|---|---|---|---|---|---|---|---|---|---|---|---|---|---|
| R | 177,641 | 1 | -13,522 | 0,48501 | R | 27,4672 | 0,99931 | 9,04872 | 0,22242 | R | 42,351 | 0,99967 | -1,7648 | 0,84162 | R | 107,823 | 1 | -20,8059 | 0,18107 |
| H | 104,238 | 1 | -16,265 | 0,40095 | H | 30,0401 | 0,99756 | 5,29145 | 0,47554 | H | 11,4542 | 1 | -0,46625 | 0,9579 | H | 62,7435 | 1 | -21,0902 | 0,17519 |
| K | 150,159 | 1 | -13,8913 | 0,47079 | K | 16,3777 | 1 | -13,4226 | 0,06776 | K | 22,1346 | 1 | -1,06981 | 0,90297 | K | 111,647 | 1 | 0,60112 | 0,96905 |
| D | 139,253 | 1 | 5,40623 | 0,78011 | D | 25,0269 | 0,99983 | -10,3959 | 0,16098 | D | 24,2952 | 1 | -3,77318 | 0,66921 | D | 89,9305 | 1 | 19,5753 | 0,20827 |
| E | 207,541 | 1 | -42,7289 | 0,02735 | E | 23,9834 | 0,99991 | -15,9465 | 0,03154 | E | 25,6805 | 1 | 1,61052 | 0,8553 | E | 157,877 | 0,99999 | -28,3929 | 0,06798 |
| S | 278,938 | 0,99994 | 62,0444 | 0,00136 | S | 29,437 | 0,99815 | 4,80842 | 0,51675 | S | 40,2188 | 0,99988 | 1,16952 | 0,89465 | S | 209,282 | 0,93691 | 56,0665 | 0,00031 |
| T | 145,228 | 1 | 13,1402 | 0,49742 | T | 25,8714 | 0,99972 | 6,54869 | 0,37722 | T | 16,4064 | 1 | 6,0564 | 0,49287 | T | 102,95 | 1 | 0,53511 | 0,97256 |
| N | 101,854 | 1 | -5,51324 | 0,7747 | N | 16,039 | 1 | -2,09914 | 0,77514 | N | 14,2222 | 1 | 3,48531 | 0,69123 | N | 71,5924 | 1 | -6,89941 | 0,65606 |
| Q | 127,543 | 1 | -19,6212 | 0,31095 | Q | 12,2343 | 1 | -6,35625 | 0,3914 | Q | 24,9884 | 1 | 4,05136 | 0,64643 | Q | 90,3204 | 1 | -17,3163 | 0,26565 |
| A | 275,145 | 0,99997 | 5,67527 | 0,76947 | A | 56,5231 | 0,41786 | 10,0478 | 0,17547 | A | 36,8012 | 0,99998 | 8,98436 | 0,30902 | A | 181,821 | 0,99852 | -13,3569 | 0,39055 |
| I | 131,641 | 1 | -22,2844 | 0,2492 | I | 24,5822 | 0,99987 | -10,6552 | 0,15079 | I | 21,1974 | 1 | 0,31489 | 0,97156 | I | 85,8615 | 1 | -11,9441 | 0,44166 |
| L | 197,538 | 1 | -28,5685 | 0,14014 | L | 22,4132 | 0,99997 | -2,84671 | 0,70109 | L | 34,1793 | 1 | -10,7669 | 0,2228 | L | 140,945 | 1 | -14,9549 | 0,33638 |
| V | 164,256 | 1 | -13,4159 | 0,48844 | V | 23,582 | 0,99993 | -4,91324 | 0,50765 | V | 35,4584 | 0,99999 | 2,2388 | 0,79989 | V | 105,216 | 1 | -10,7415 | 0,48989 |
| F | 145,28 | 1 | -9,19004 | 0,63464 | F | 20,2589 | 1 | -4,13733 | 0,57693 | F | 24,7953 | 1 | -14,6199 | 0,09785 | F | 100,225 | 1 | 9,5672 | 0,53771 |
| W | 81,3488 | 1 | 9,13389 | 0,63305 | W | 14,2621 | 1 | 6,06124 | 0,41376 | W | 11,9919 | 1 | -2,84992 | 0,74535 | W | 55,0948 | 1 | 5,92257 | 0,69863 |
| Y | 137,628 | 1 | -6,54392 | 0,73405 | Y | 19,4967 | 1 | -14,9079 | 0,04441 | Y | 19,1168 | 1 | -2,46614 | 0,77868 | Y | 99,0148 | 1 | 10,8301 | 0,48359 |
| M | 85,777 | 1 | -19,9739 | 0,29778 | M | 17,7007 | 1 | 0,66401 | 0,928 | M | 14,7746 | 1 | -5,19602 | 0,55376 | M | 53,3018 | 1 | -15,4419 | 0,31583 |
| C | 104,193 | 1 | -10,0069 | 0,59993 | C | 12,039 | 1 | -6,06273 | 0,40935 | C | 16,4919 | 1 | -0,77839 | 0,92932 | C | 75,6619 | 1 | -3,16583 | 0,8357 |
| G | 178,568 | 1 | 2,84623 | 0,88315 | G | 27,4145 | 0,99933 | 11,4667 | 0,12206 | G | 21,0289 | 1 | -0,84 | 0,92423 | G | 130,125 | 1 | -7,78046 | 0,61697 |
| P | 427,888 | 0,03065 | 21,1201 | 0,27543 | P | 67,3212 | 0,12313 | 22,6067 | 0,0023 | P | 63,2568 | 0,88685 | 3,17688 | 0,71906 | P | 297,31 | 0,00879 | -4,66352 | 0,76434 |

S12: Putative transit peptide region amino acid composition analysis results: The predicted "TP" region was compared to the predicted mature chain of the same sequence. This was performed a) for both experimental and control protein sets (two rows of tables), b*)* for the whole sets of 375 proteins regardless of their classification as well as for each subset representing a class of pre-proteins (four columns of tables), and c) for each amino acid (twenty rows in each table). $\chi^2$ sums and their respective *p*-values, as well as normalized $\chi^2$ sums (sum of $\chi^2$ with plus or minus sign depending on the positive or negative value of its residual) and their *p*-values are shown in each table. *P*-values lower than 0.01 are colored in yellow, *p*-values lower than 0.00001 are colored in light orange.

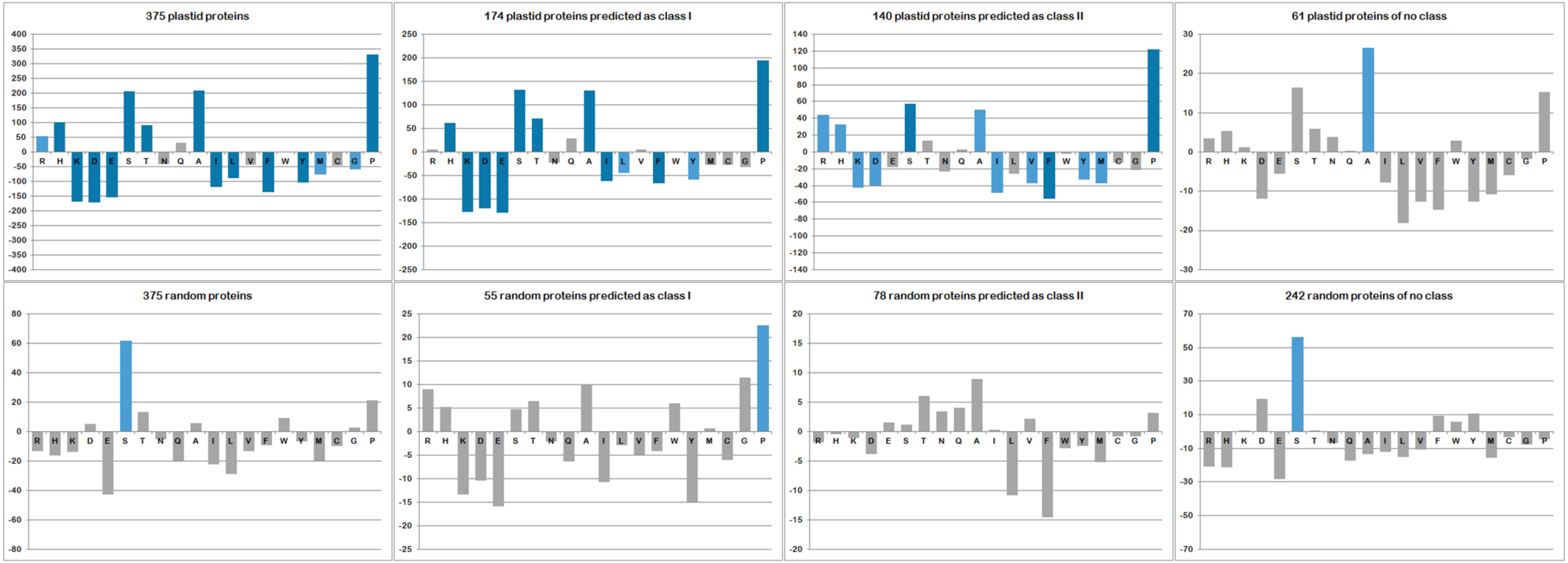


S13: Graphical representation of normalized $\chi^2$ sums for each amino acid in the TP vs. mature chain comparison in the two sets: positive or negative value reflects whether the amino acid frequency in the “TP” region is higher or lower than expected, colored bars represent statistically significant differencies with $p<0.01$ (lighter blue) and $p<0.00001$ (darker blue).
